# Behavioral and neurochemical effects of novel N-Benzyl-2-phenylethylamine derivatives in adult zebrafish

**DOI:** 10.1101/2022.01.19.476767

**Authors:** Konstantin A. Demin, Olga V. Kupriyanova, Vadim A. Shevyrin, Ksenia A. Derzhavina, Nataliya A. Krotova, Nikita P. Ilyin, Tatiana O. Kolesnikova, David S. Galstyan, Iurii M. Kositsyn, Abubakar-Askhab S. Khaybaev, Maria V. Seredinskaya, Yaroslav Dubrovskii, Raziya G. Sadykova, Maria O. Nerush, Mikael S. Mor, Elena V. Petersen, Tatyana Strekalova, Evgeniya V. Efimova, Dmitrii V. Bozhko, Vladislav O. Myrov, Sofia M. Kolchanova, Aleksander I. Polovian, Georgii K. Galumov, Allan V. Kalueff

## Abstract

Serotonergic hallucinogenic drugs potently affect human brain and behavior, and have recently emerged as potentially promising agents in psychopharmacotherapy. Complementing rodent studies, zebrafish (*Danio rerio*) is a powerful animal model for screening neuroactive drugs, including serotonergic agents. Here, we test ten different N-Benzyl-2-phenylethylamine (NBPEA) derivatives with the 2,4- and 3,4-dimethoxy substitutions in the phenethylamine moiety and the - OCH3, -OCF3, -F, -Cl and -Br substitutions in the *ortho* position of phenyl ring of *N*-benzyl fragment, assessing their behavioral and neurochemical effects in adult zebrafish. Overall, substitutions in *N*-benzyl fragment primarily affected zebrafish locomotion, and in phenethylamine moiety - anxiety-like behavior, also modulating brain serotonin and/or dopamine turnover. We also identified several behavioral clusters, including anxiogenic/hypolocomotor (24H-NBF, 24H-NBOMe and 34H-NBF), behaviorally inert (34H-NBBr, 34H-NBCl and 34H- NBOMe), anxiogenic/hallucinogenic-like (24H-NBBr, 24H-NBCl and 24H-NBOMe(F)), and anxiolytic/hallucinogenic-like (34H-NBOMe(F)) agents. The 24H-NBOMe(F) and 34H-NBOMe(F) also reduced despair-like behavior in zebrafish. The artificial intelligence-driven phenotyping supports association of multiple compounds with NMDA antagonists and/or MDMA, supporting their potential hallucinogenic-like properties, as well as other valuable psychoactive effects. *In silico* functional molecular activity modelling also supports existing of similarities between studied NBPEAs drugs, MDMA, and ketamine. Functional analysis implicates potential involvement of serotonin release stimulating activity, calcium channel (voltage-sensitive) activity, some serotonin receptors activity and variety of psychiatric and neurologic disorders treatments activities. Overall, we report potent neuroactive properties of several novel synthetic *N*-benzylphenylethylamines in an *in vivo* vertebrate model system (zebrafish), raising the possibility of their potential use in clinical practice.

## Introduction

Serotonergic psychedelic drugs potently affect human brain and behavior, evoking pronounced hallucinogenic states, including sensory alterations, ego dissolution and mystical/spiritual-like experiences^1–4^. These drugs have recently emerged as potentially promising compounds for translational medicine and pharmacotherapy^2, 5–7^. For example, psilocybin improves mood and anxiety in terminal cancer patients^8^ and reduces symptoms of treatment-resistant depression^9^. Mood improvement also occurs following lysergic acid diethylamide (LSD) and psilocybin therapy^10–12^, together with MDMA-assisted psychotherapy for post-traumatic stress disorder (PTSD)^13^. Paralleling human studies, experimental animal models have long been used to study psychedelic drugs, showing improved cognitive abilities, reduced depression-like behavior and increased cortical plasticity by psilocybin^8^, and potentiated neurogenesis and synaptogenesis in rat cultured neurons and in Drosophila larvae by LSD^14^.

The *N*-(2-methoxy)benzyl derivatives of the phenethylamines (NBOMes) represent a new family of synthetic hallucinogenic drugs, whose member, 25I-NBOMe, has been extensively studied for its robust 5HT_2A_ activity ^15–17^. Alternative phenethylamines (2C family), such as NBOMes, represent designer drugs that have methoxy groups in the 2 and 5 positions of the phenyl ring of the phenethylamine moiety, with various lipophilic groups at position 4^18^. The “2C” group of psychedelic phenethylamines, first described by the Shulgins^19^, includes the two carbons between the phenyl ring and the amino group C^20^. NBOMes, as derivatives of the 2C family, exert LSD-like central nervous system (CNS) effects clinically^21^, including hallucinations, euphoria, visual and sensory effects, mystical experiences and altered consciousness^21, 22^. NBOMes are considerably toxic, and their intake evokes severe negative side effects, including mortality^23^.

Major CNS effects of classical serotonergic hallucinogens (e.g., LSD and psilocybin) and NBOMes involve agonism at serotonin 5-HT_2A_ receptors^24^ that are strongly implicated in various mental processes, including working memory and affective (e.g., depression) or psychotic (e.g., schizophrenia) deficits^25, 26^. Modulating 5-HT_2A_ receptors may also be effective for various neurological disorders, including absence seizures, Tourette syndrome, autism and Alzheimer’s disease (AD)^20, 27–30^.

Highly efficient and selective 5-HT_2A_ agonists, NBOMes are 1000-fold more selective to these (vs. 5-HT_1A_) receptors^31–35^. They also display high affinity to 5-HT_2C_ receptors, α1 adrenergic receptors and histamine H1 receptor, but low affinity to dopamine D1-3 and trace amine (TAAR1) receptors and monoamine (dopamine, serotonin and norepinephrine) transporters DAT, SERT and NET^32, 34, 35^.

In addition to monoaminergic system, NBOMes also modulate the glutamatergic system^36^, as some of them (e.g., 25I-NBOMe) elevate extracellularglutamate in rat frontal cortex^37^, nucleus accumbens (NAc) shell and medial prefrontal cortex (mPFC)^36^. In rodents, the 5HT_2A_ receptor activation induces characteristic ‘head twitch’ response (HTR), a rapid paroxysmal head rotation^38–40^ widely used as a behavioral marker for hallucinogens (e.g., LSD or psilocybin) in rodents^38, 41^.

In line with this, 25C-NBOMe, 25I-NBOMe and 25B-NBOMe all induce overt HTR responses in rodents^34, 37, 42, 43^. Unlike NBOMes, classical 2C compounds display relatively low 5HT_2A_ affinity, with potential partial agonism or antagonism^44, 45^, despite typical hallucinogenic profile in both clinical^46^ and animal^34^ studies. Collectively, this calls for further studiesof neurobehavioral effects of serotonergic psychedelics and their derivatives, in order to better understand their mechanisms of action and potential to treat neuropsychiatric disorders.

Complementing rodent studies, zebrafish (*Danio rerio*) is a powerful in vivo model system for CNS drug screening^47, 48^ and deliriants^49^. Zebrafish models are particularly suitable for such screening due to high genetic and physiological homology to humans, easy husbandry, low cost, fast reproduction, and sensitivity to a wide range of psychoactive drugs^50, 51^, including serotonergic psychedelics^52, 53^ glutamatergic dissociatives^54^ and cholinergic deliriants^49^.

Capitalizing on zebrafish as a powerful screening platform, here we tested ten different N-Benzyl-2-phenylethylamine (NBPEA) derivatives with the 2,4- and 3,4-dimethoxy substitutions in the phenethylamine moiety and the -OCH_3_, -OCF_3_, -F, -Cl and -Br substitutions in the *ortho* position of phenyl ring of benzyl fragment, assessing their behavioral and neurochemical effects in adult zebrafish. The studied compounds represented 5-HT_2A_ receptor agonists not currently controlled by law, and included *N*-(2-methoxybenzyl)-2-(2,4-dimethoxyphenyl)ethanamine (24H-NBOMe) (**1**), *N*-(2-trifluoromethoxybenzyl)-2-(2,4-dimethoxyphenyl)ethanamine (24H-NBOMe(F) (**2**) *N*-(2-fluorobenzyl)-2-(2,4-dimethoxyphenyl)ethanamine (24H-NBF) (**3**), *N*-(2-chlorobenzyl)-2-(2,4-dimethoxyphenyl)ethanamine (24H-NBCl) (**4**), *N*-(2-bromobenzyl)-2-(2,4-dimethoxyphenyl)ethanamine (24H-NBBr) (**5**), *N*-(2-methoxybenzyl)-2-(3,4-dimethoxyphenyl)ethanamine (34H-NBOMe) (**6**), *N*-(2-trifluoromethoxybenzyl)-2-(3,4-dimethoxyphenyl)ethanamine (34H-NBOMe(F) (**7**) *N*-(2-fluorobenzyl)-2-(3,4-dimethoxyphenyl)ethanamine (34H-NBF) (**8**), *N*-(2-chlorobenzyl)-2-(3,4-dimethoxyphenyl)ethanamine (34H-NBCl) (**9**), *N*-(2-bromobenzyl)-2-(3,4-dimethoxyphenyl)ethanamine (34H-NBBr) (**10**) (**Fig. 1**).

**Figure 1.**
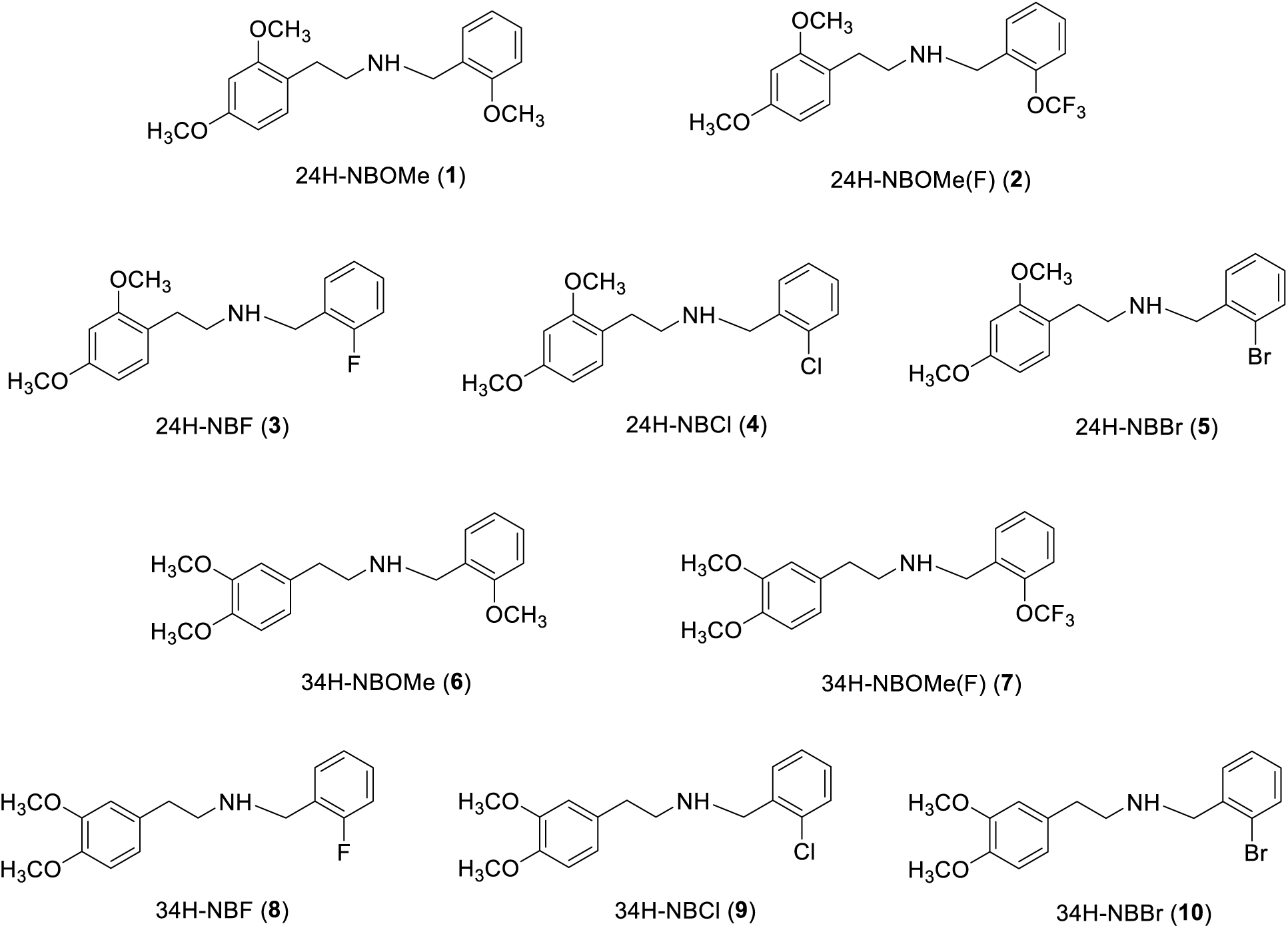
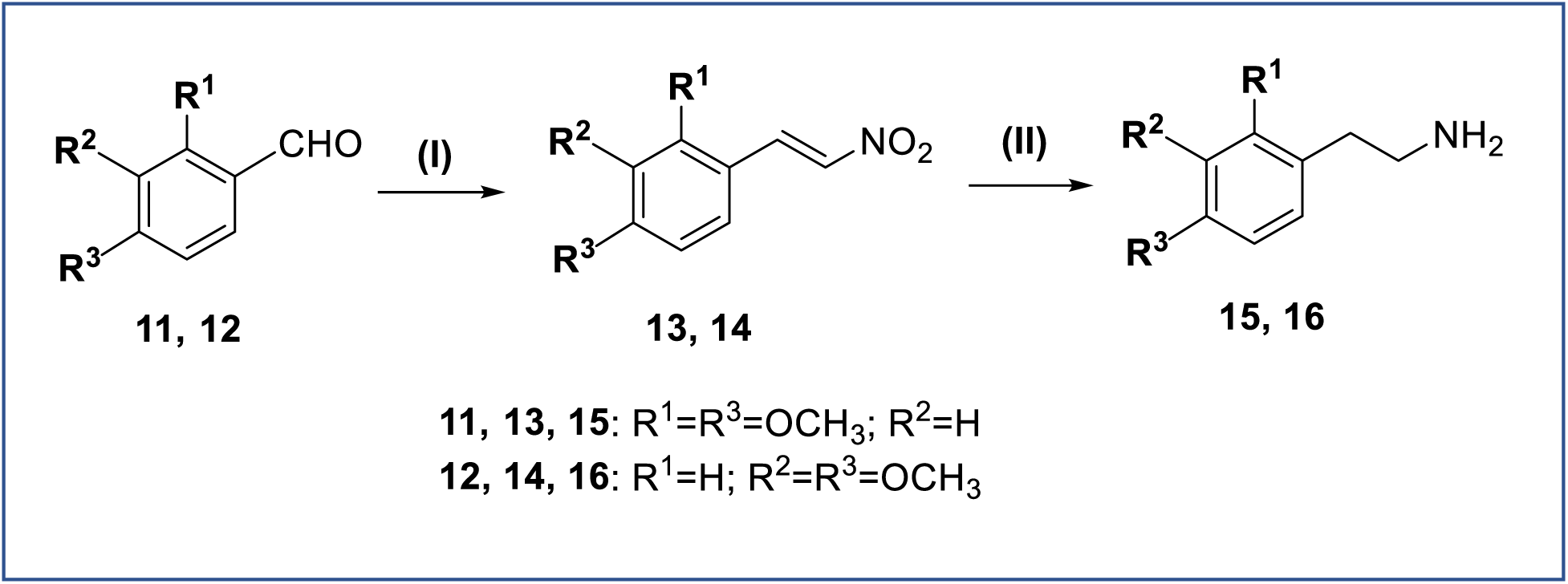
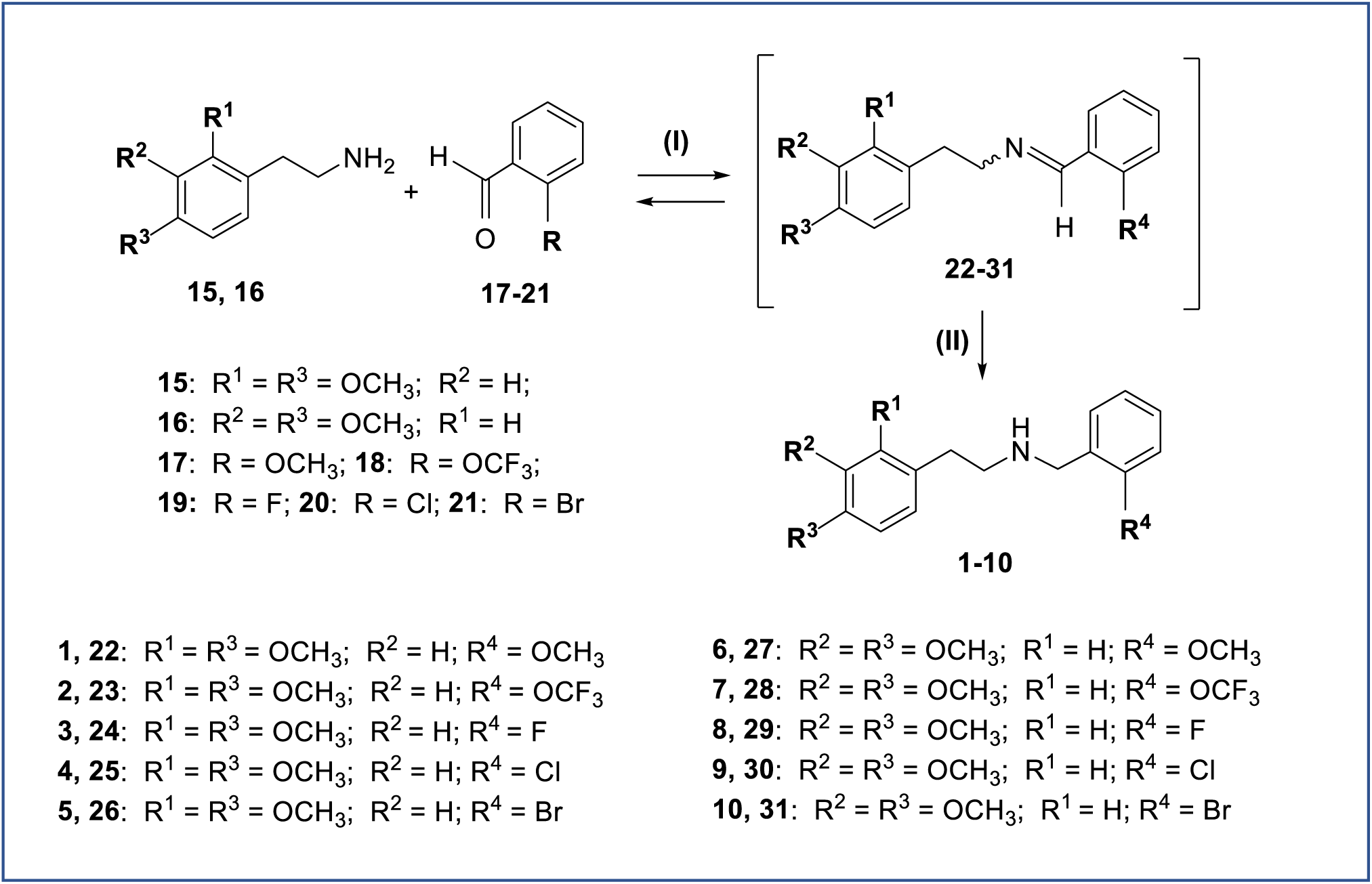
Structures of compounds **1**–**10** investigated in the present study. Top inset: Synthesis of compounds **13, 14** and **15, 16**. Reagents and conditions: (**I**) Henry reaction – CH_3_NO_2_, NH_4_OAc, reflux 100°C (4 h); (**II**) reduction – LiAlH_4_, THF, 0°C. Bottom inset: Synthesis of compounds **1-10** via reductive amination. Reagents and conditions: (**I**) Condensation – CH_2_Cl_2_, 10°C (4-5 h); (**II**) reduction – NaBH_4_, MeOH, 0°C.

For compounds **1, 2, 6, 7** their chemical synthesis and analytical data have already been reported^55^. The structures of compounds **2-5 and 7-10** were confirmed by ^1^H NMR, ^13^C NMR and ^19^F NMR (for compounds **2, 3, 7, 8** only) spectroscopy, gas chromatography with mass spectrometric detection (GC-MS), ultra-high-performance liquid chromatography with high resolution mass spectrometric detection (UHPLC-HRMS), elemental analysis data (except **2, 3, 7, 8**), and melting point determination. The present study assessed potential as CNS therapies of two groups of 24-and 34-dimethoxy-N-Benzyl-2-phenylethylamine (NBPEA), with various substituents (-OCH_3_, OCF_3_, -F, -Cl and -Br) at the *ortho* position of phenyl ring of *N*-benzyl fragment, focusing on their neurobehavioral and neurochemical effects in adult zebrafish.

## Results and discussion

### Chemistry

Compounds **2-5**, **7-10** were synthesized similarly to^55^, involving synthesis of 2,4-dimethoxy-*β*-nitrostyrene (**13)** and 3,4-dimethoxy-*β*-nitrostyrene (**14)** via Henry reaction^56^ with subsequent reduction to substituted phenethylamines **15, 16** (Fig. 1). Final synthesis of compounds **1-10** involved reductive *N*-amination through the formation of the corresponding intermediate imine products (**22-31,** Fig. 1).

### Behavioral studies

Tested compounds exerted significant concentration- and drug-specific effects on zebrafish behaviors in zebrafish novel tank test of anxiety and activity. The 24H-NBOMe exposure exerted significant effects on fish behavior, concentration-dependently reducing both the time spent in top and the number of top entries at 10 and 20 mg/L (Fig. 2 and Supplementary Table S1). 24H-NBF affected only top entries, concentration-dependently reducing it at 20 mg/L (Supplementary Table S1). 24H-NBCl reduced top dwelling behavior in zebrafish as well, concentration-dependently reducing time in top and the number of top entries at 10 and 20 mg/L (Fig. 2 and Supplementary Table S1). The 20 mg/L 24H-NBCl group displayed significantly more horizontal shuttling behavior (see the Methods section for details) vs. controls (Fig. 3 and Supplementary Table S1). The 24H-NBBr-treated fish showed significant drug effect on velocity, rising it at 10 mg/L and decreasing top entries at 10 and 20 mg/L (Supplementary Table S1). There was also a significant effect for horizontal shuttling behavior, as the drug increased it at both 10 and 20 mg/L vs. controls (Fig. 3 and Supplementary Table S1).

**Figure 2.**
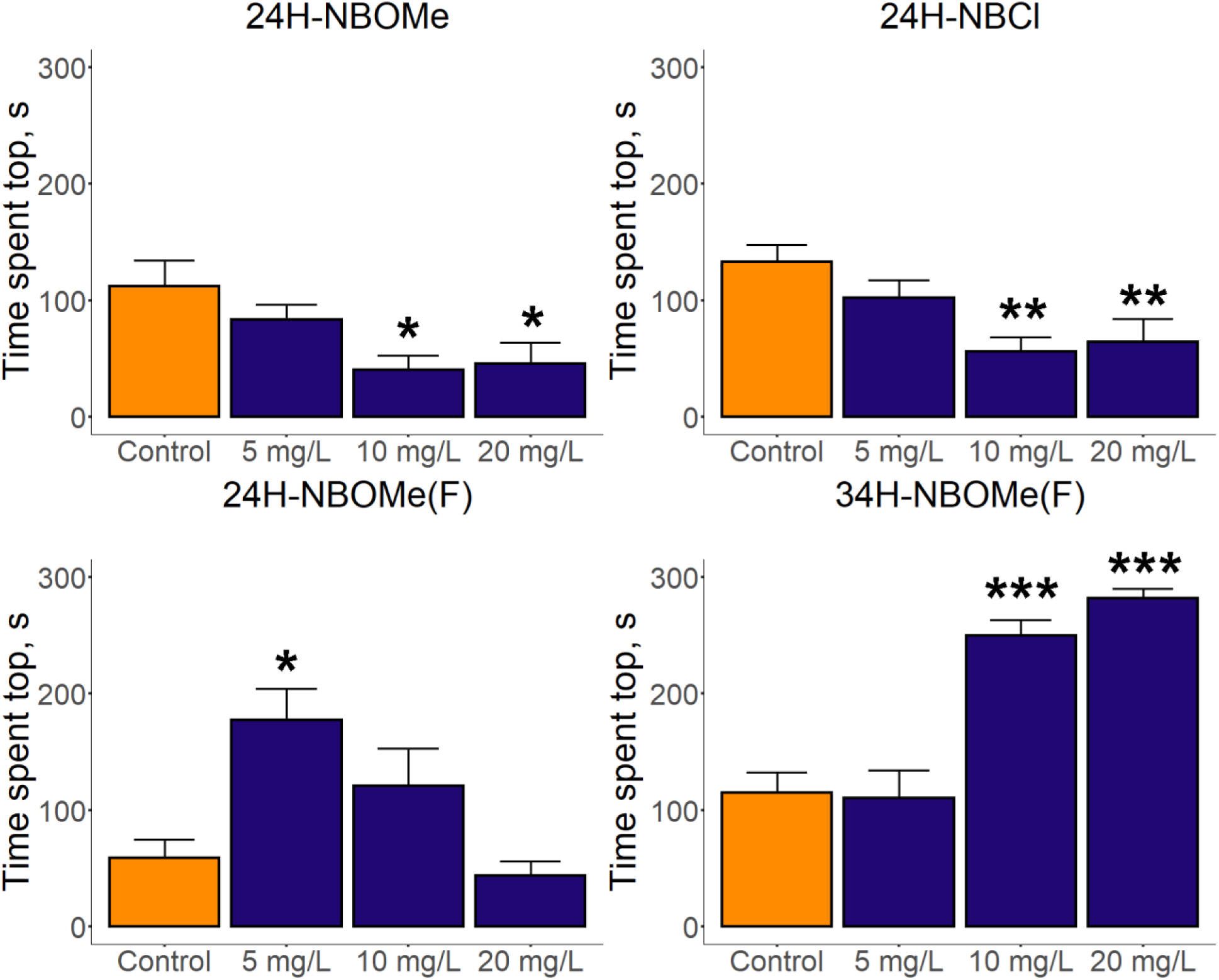
Behavioral effects an anxiety-related time in top behavior assessed in the novel tank test (NTT) induced by selected N-Benzyl-2-phenylethylamine (NBPEA) derivatives. Increase in time spent top corresponds to an anxiolytic effect. Data is represented as mean ± S.E.M. (n=15-17). *p<0.05, **p<0.01, ***p<0.001 vs. control, post-hoc Dunn’s test for significant Kruskal-Wallis data. Graphs were constructed using the ggplot2 R package^135^, also see Supplementary Table S1 for statistical details.

**Figure 3.**
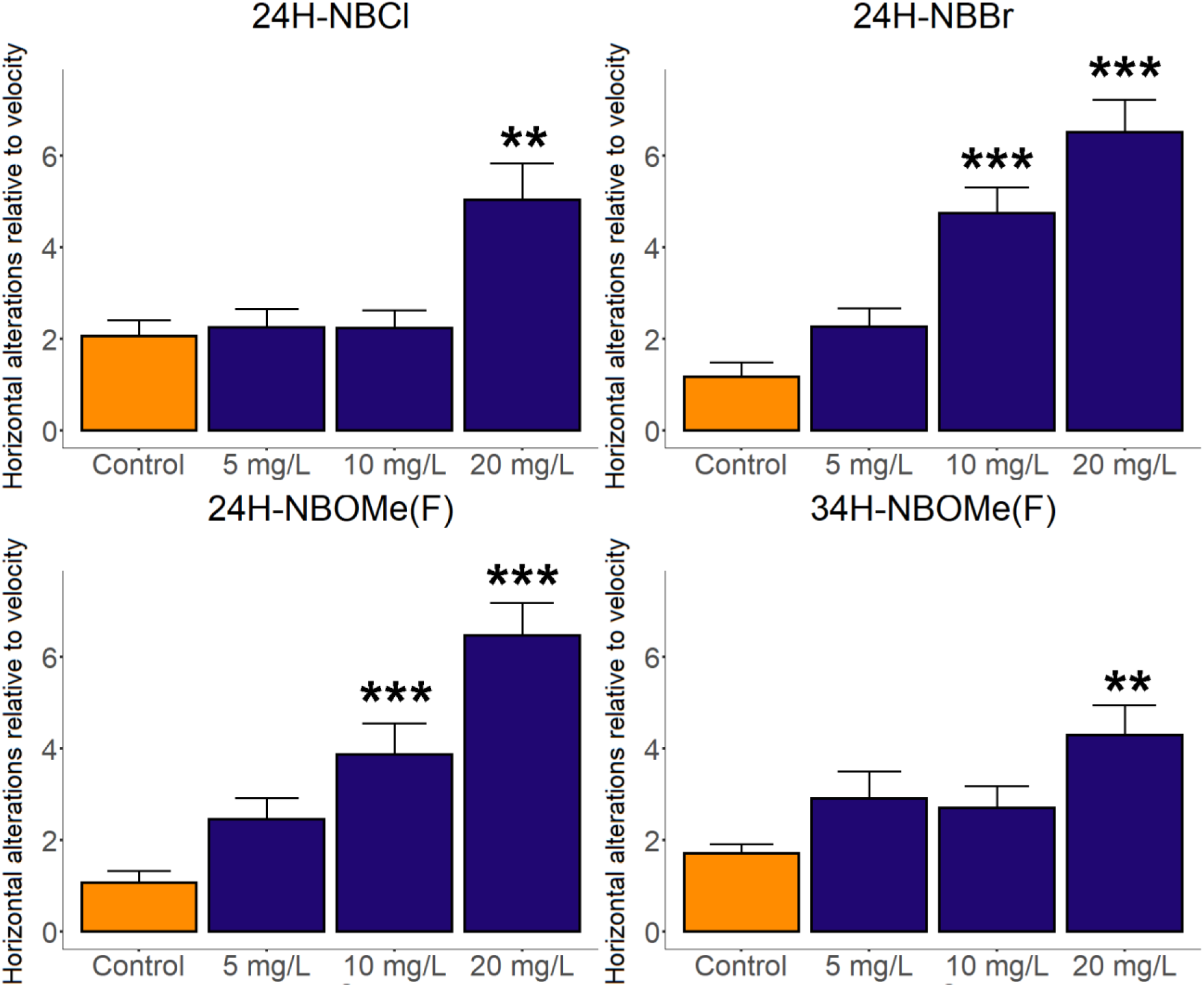
Behavioral effects observed in the novel tank test induced by selected N-Benzyl-2-phenylethylamine (NBPEA) derivatives, assessed as horizontal ‘shuttling’ (normalized horizontal transitions) calculated as total left-to-right or right-to-left horizontal transitions divided by distance traveled. Increase in shuttling behavior corresponds to potential increase in hallucinogenic-like properties. Data are presented as mean ± S.E.M. (n=15-17). *p<0.05, **p<0.01, ***p<0.001 vs. control, post-hoc Dunn’s test for significant Kruskal-Wallis data. Graphs were constructed using the ggplot2 R package^135^, also see Supplementary Table S1 for summary of statistical details.

24H-NBOMe(F) increased zebrafish velocity (at 5 and 10 mg/L) and time in top (at 5 mg/L), reduced the number of top entries at 10 mg/L and the latency to top at 5 mg/L, and promoted horizontal shuttling behavior at 5 and 10 mg/L (Fig. 2-3 and Supplementary Table S1). The 34H-NBF exposure only decreased the latency to top at 10 mg/L, and 34H-NBCl and 34H-NBBr evoked more top entries at 20 mg/L (Supplementary Table S1). 34H-NBOMe(F) increased time in top and reduced the number of top entries and the latency to enter the top at 10 and 20 mg/L (Fig. 2 and Supplementary Table S1). Furthermore, exposure to 34H-NBOMe(F) increased horizontal shuttling behavior at 20 mg/L, whereas 34H-NBOMe was generally behaviorally inactive at all concentrations tested (Fig. 3, Supplementary Table S1).

Generalized linear models (GZLM) analyzed the role of individual substitutions in *N*-benzyl fragment and in phenethylamine moiety, as well as their interactions, at the most behaviorally active concentration (20 mg/L) across all compounds (Fig. 5 and 6, Supplementary Tables S4-S8). Significant Wald test effect was found for substitutions in the *N*-benzyl fragment for velocity, with significant post-hoc differences between -F vs. -OMe(F), -Br or -Cl substitutions (Fig. 5, Supplementary Tables S5 and S7). Horizontal shuttling behavior yielded significant substitution effects in both the phenethylamine moiety and the *N*-benzyl fragment (Fig. 5, Supplementary Tables S5-7). Time spent in top yielded significant Wald test effect for substitutions in the phenethylamine moiety, with post-hoc test showing significant differences between 24H- and 34H-derivatives (Fig. 5, Supplementary Tables S5 and S6). Significant Wald test effects were also found for substitutions in the phenethylamine moiety for the latency to enter the top (24H- vs. 34H-), and the number of top entries for substitutions both in the phenethylamine moiety and the *N*-benzyl fragment as well as their interaction (Fig. 5, Supplementary Tables S4-S8). Finally, in the zebrafish tail immobilization (ZTI) test of despair-like behavior, 34H-NBOMe(F) evoked strong behavioral effects, as assessed both by more distance travelled and time spent mobile vs. control, whereas 24H-NBOMe(F) had less pronounced antidepressant-like effect with longer time spent mobile (Fig. 7, Supplementary Table S9).

**Figure 5.**
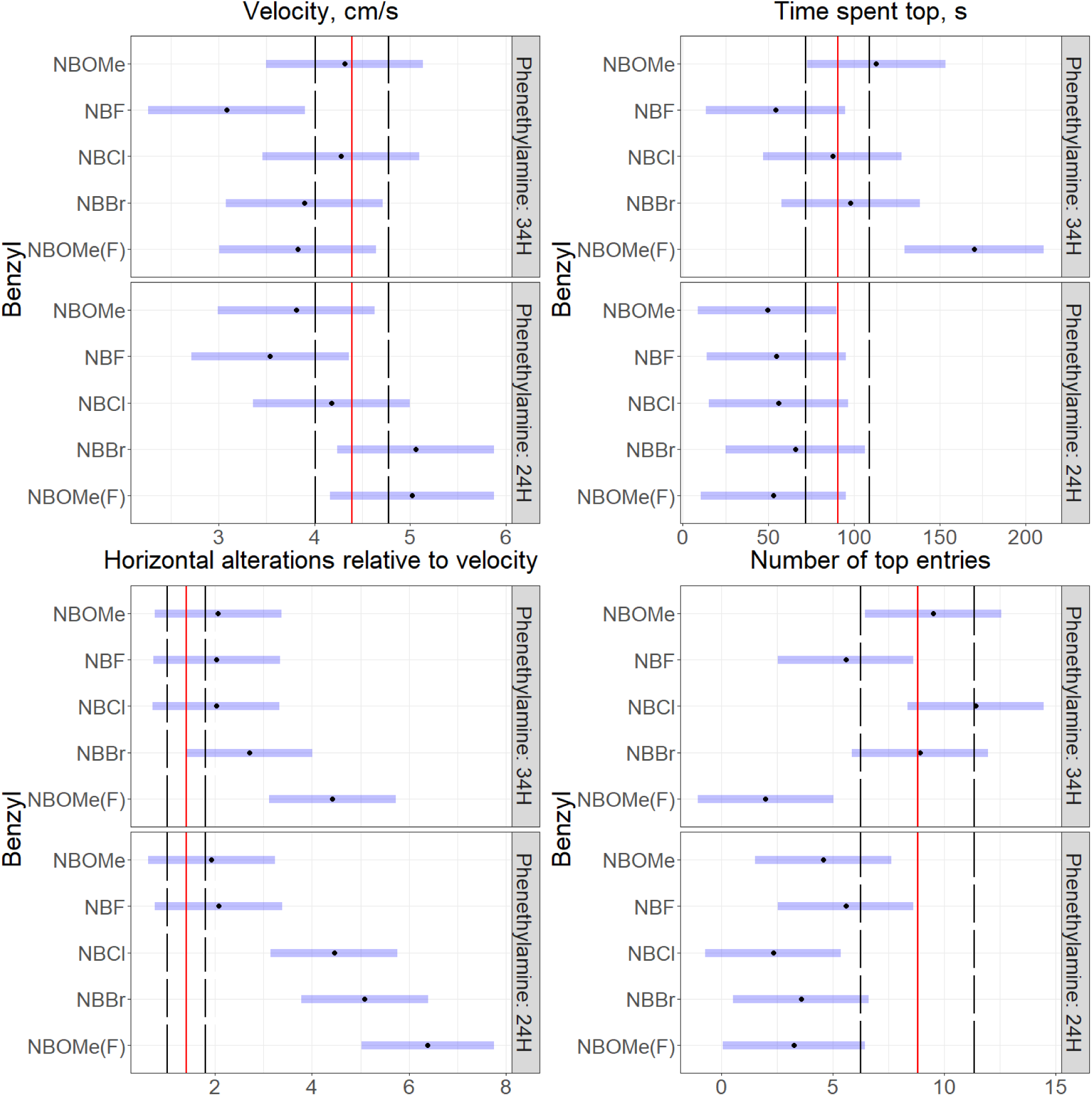
Summary of GZLM analyses (factors benzyl, and phenethylamine derivatives, and their interaction) comparing behavior of all synthesized N-Benzyl-2-phenylethylamine (NBPEA) compounds at the same 20 mg/L concentration in the novel tank test. Data are presented as estimated marginal means (emmeans)^136^. Increase in time spent top corresponds to an anxiolytic effect. Increase in shuttling behavior corresponds to potential increase in hallucinogenic-like properties. Representative control, that was excluded from statistical analysis is shown graphically as mean (red vertical line) ± S.E.M. (black dotted lines; n = 8-10). Graphs were constructed using the ggplot2 R package^135^, also see Supplementary Tables S4-8 for summary of statistical details.

**Figure 6.**
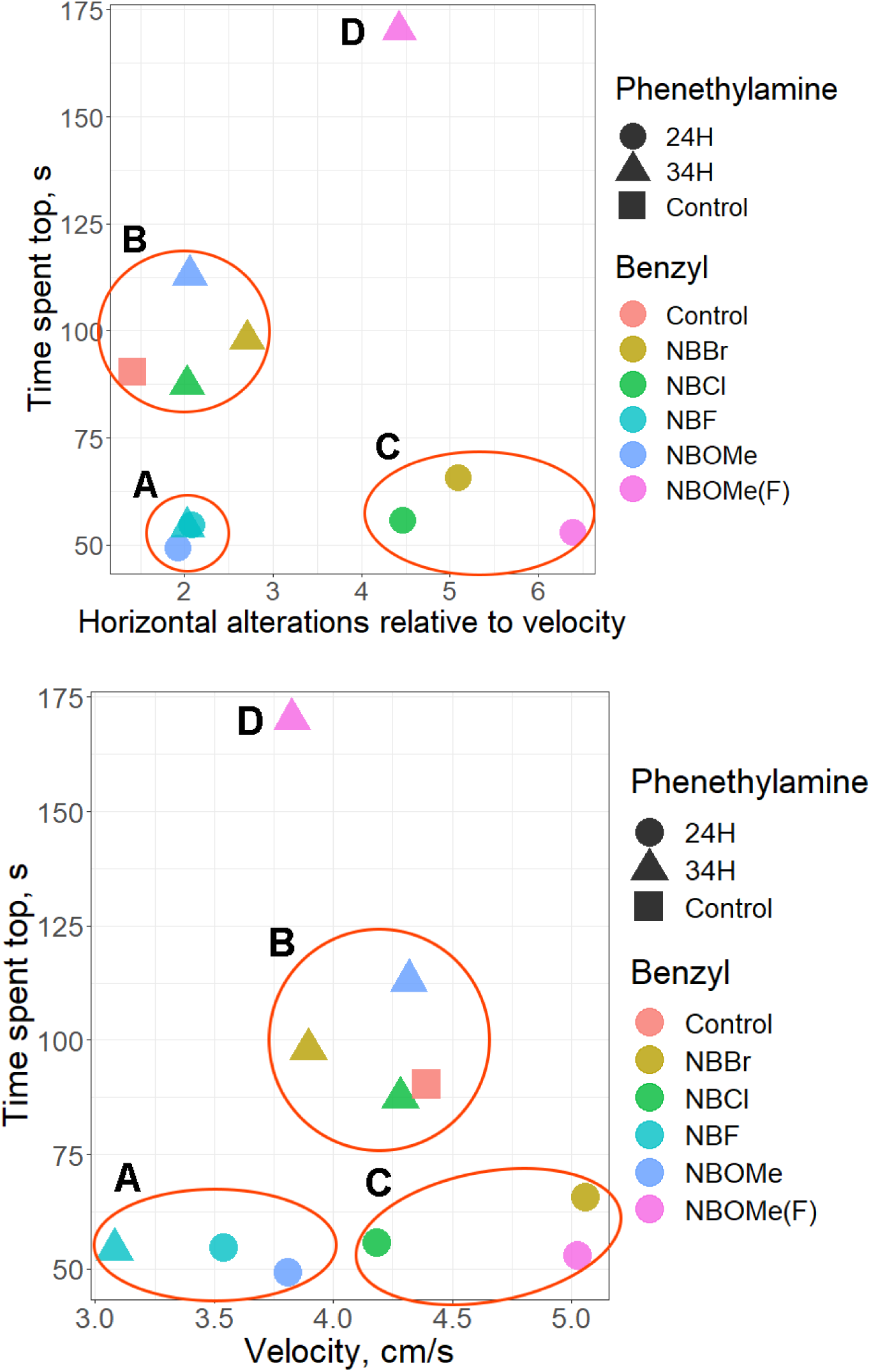
The 3D map of group means behavioral endpoints observed in GZLM analyses comparing all synthesized N-Benzyl-2-phenylethylamine (NBPEA) compounds in the similar 20 mg/L concentration in the novel tank test. Data are presented as mean (n=8-10). 4 clusters can be easily distinguished that is proven by k-means analysis (Supplementary Table S10). Graphs were constructed using the ggplot2 R package^135^, also see Supplementary Tables S4-S8 for statistical details and https://www.stressandbehavior.com/derriatives-testing-result for the htlm 3D plot version of standardized values used for k-means analysis. Increase in time spent top corresponds to an anxiolytic effect. Increase in shuttling behavior corresponds to potential increase in hallucinogenic-like properties. A – anxiogenic and hypolocomotor cluster (24H-NBF, 24H-NBOMe and 34H-NBF), B – control-like cluster (Control, 34H-NBBr, 34H-NBCl, 34H-NBOMe), C – anxiogenic and hallucinogenic-like cluster (24H-NBBr, 24H-NBCl, 24H-NBOMe(F)), D – highly anxiolytic and hallucinogenic cluster (34H-NBOMe(F)).

**Figure 7.**
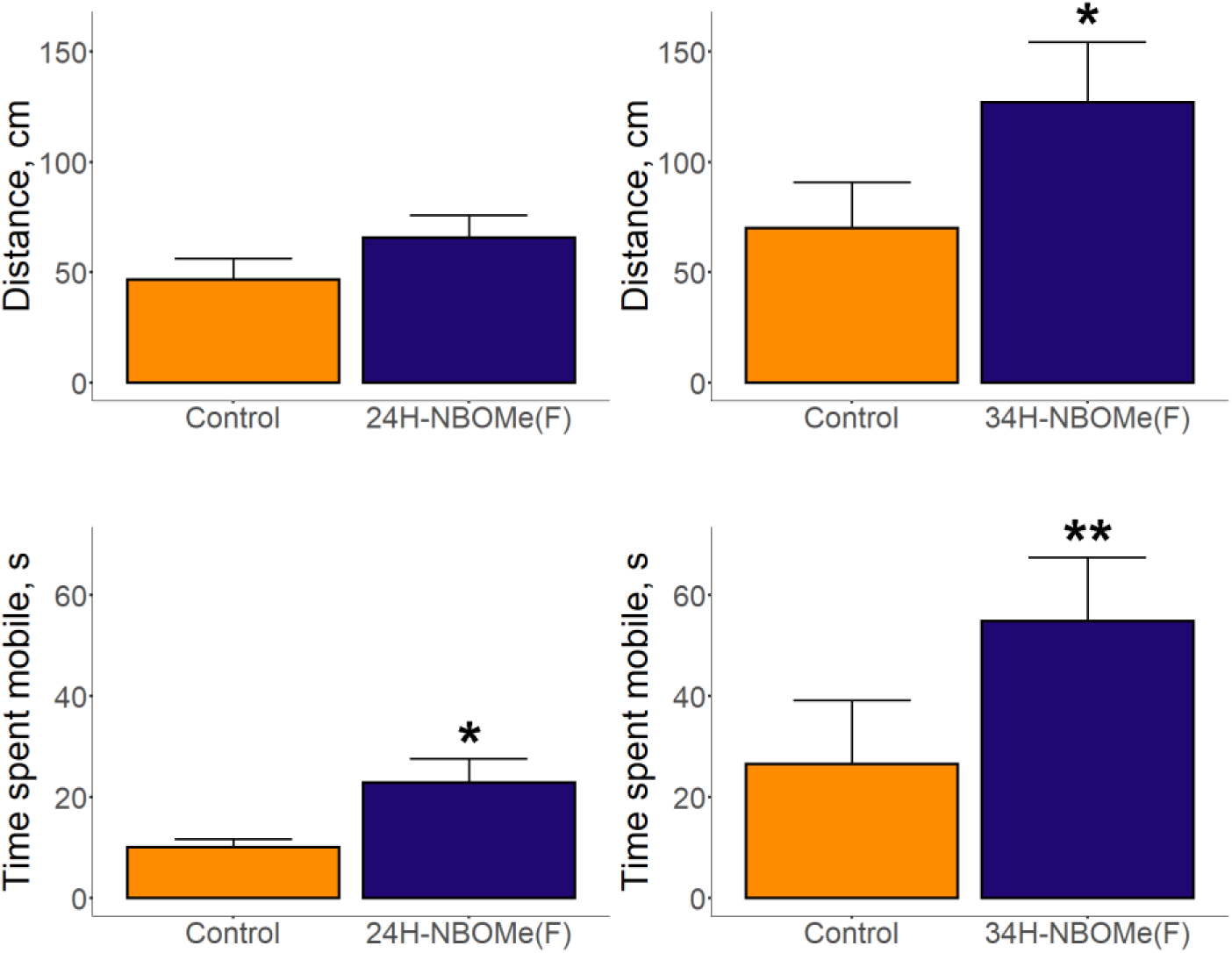
Behavioral effects in the zebrafish tail immobilization (ZTI) test assessing despair-like behavior induced by anxiolytic N-Benzyl-2-phenylethylamine (NBPEA) derivatives for distance traveled and time spent mobile. Increase in test activity corresponds to antidepressant-like drug effect. Data is represented as mean ± S.E.M. (n=13-15). *p<0.05, **p<0.01, ***p<0.001 vs. control, U-test. Graphs were constructed using the ggplot2 R package^135^, also see Supplementary Table S9 for statistical details.

### Neurochemical effects

Neurochemical analyses revealed significant drug effects, with lower serotonin levels in adult zebrafish brain for 24H-NBF, 24H-NBBr, 24H-NBOMe(F) and 34H-NBOMe(F) vs. control groups (see Supplementary Table S3 for details of post-hoc Dunn’s test for significant KW data). Furthermore, 24H-NBF, 24H-NBBr, 24H-NBOMe(F), 34H-NBCl and 34H-NBOMe(F) all increased serotonin turnover (the 5-HIAA/serotonin ratio) vs. controls (Fig. 4 and Supplementary Table S3). Similarly, 24H-NBBr, 24H-NBOMe(F), 34H-NBCl and 34H-NBOMe(F), but not 24H-NBF, reduced dopamine levels and increased dopamine turnover (the DOPAC/dopamine ratio) vs. control group (Fig. 4 and Supplementary Table S3). The HVA/dopamine ratio increased after 24H-NBOMe(F), 34H-NBCl and 34H-NBOMe(F) exposure vs. controls (Supplementary Table S3).

**Figure 4.**
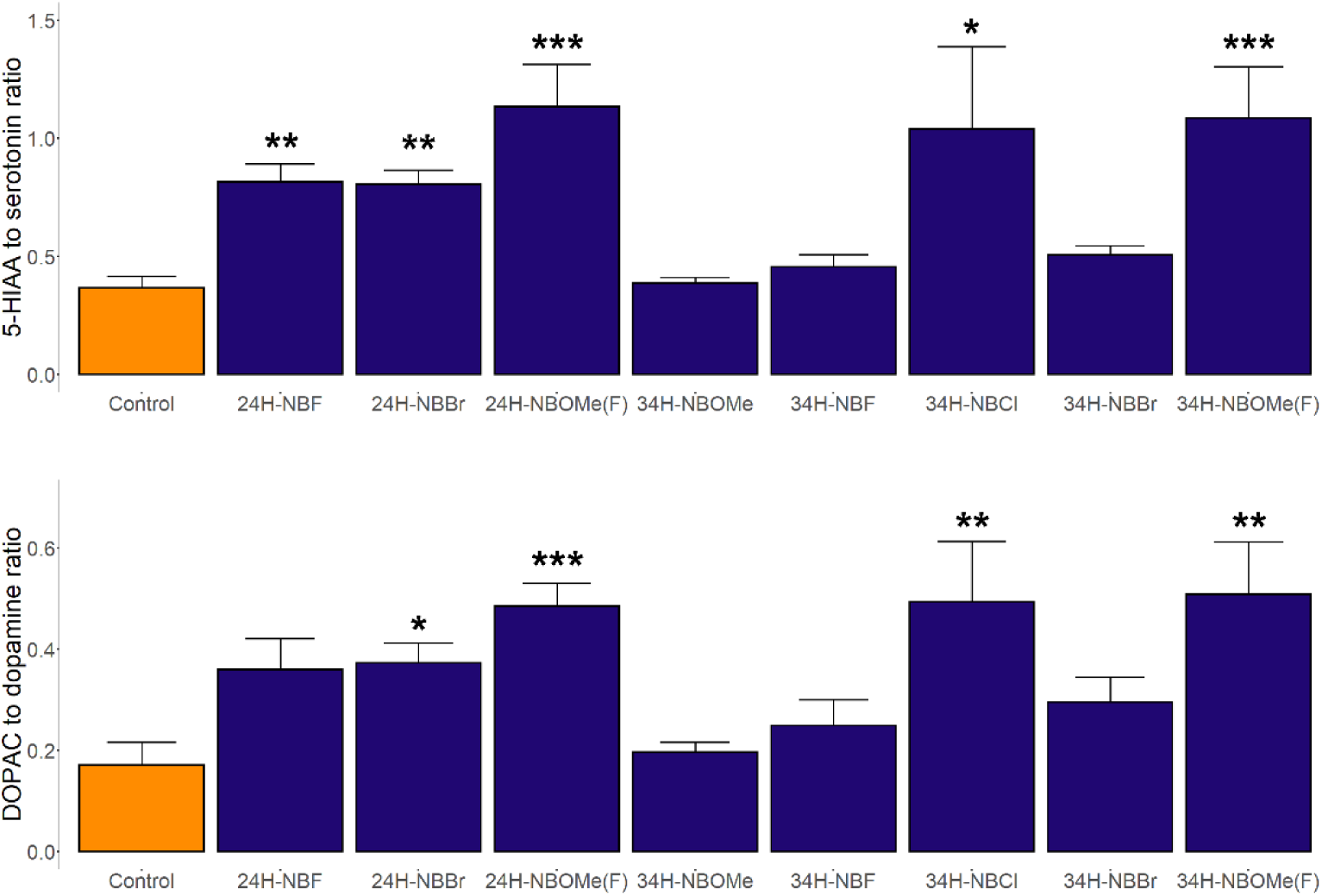
Alterations in serotonin and dopamine metabolism in adult zebrafish brain induced by selected N-Benzyl-2-phenylethylamine (NBPEA) derivatives. Data are represented as mean ± S.E.M. (n = 8-10). *p<0.05, **p<0.01, ***p<0.001 vs. control, post-hoc Dunn’s test for significant Kruskal-Wallis data. Graphs were constructed using the ggplot2 R package^135^, also see Supplementary Table S3 for statistical details.

Finally, our UHPLC-HRMS analyses detected overt N-Benzyl-2-phenylethylamines (NBPEAs) peaks in zebrafish brain samples for all derivatives tested here, indicating that all drugs reached zebrafish CNS at 20 mg/L, the systemic concentration which also showed behavioral and neurochemical effects in the present study (Fig. 8 and Table 1).

**Figure 8.**
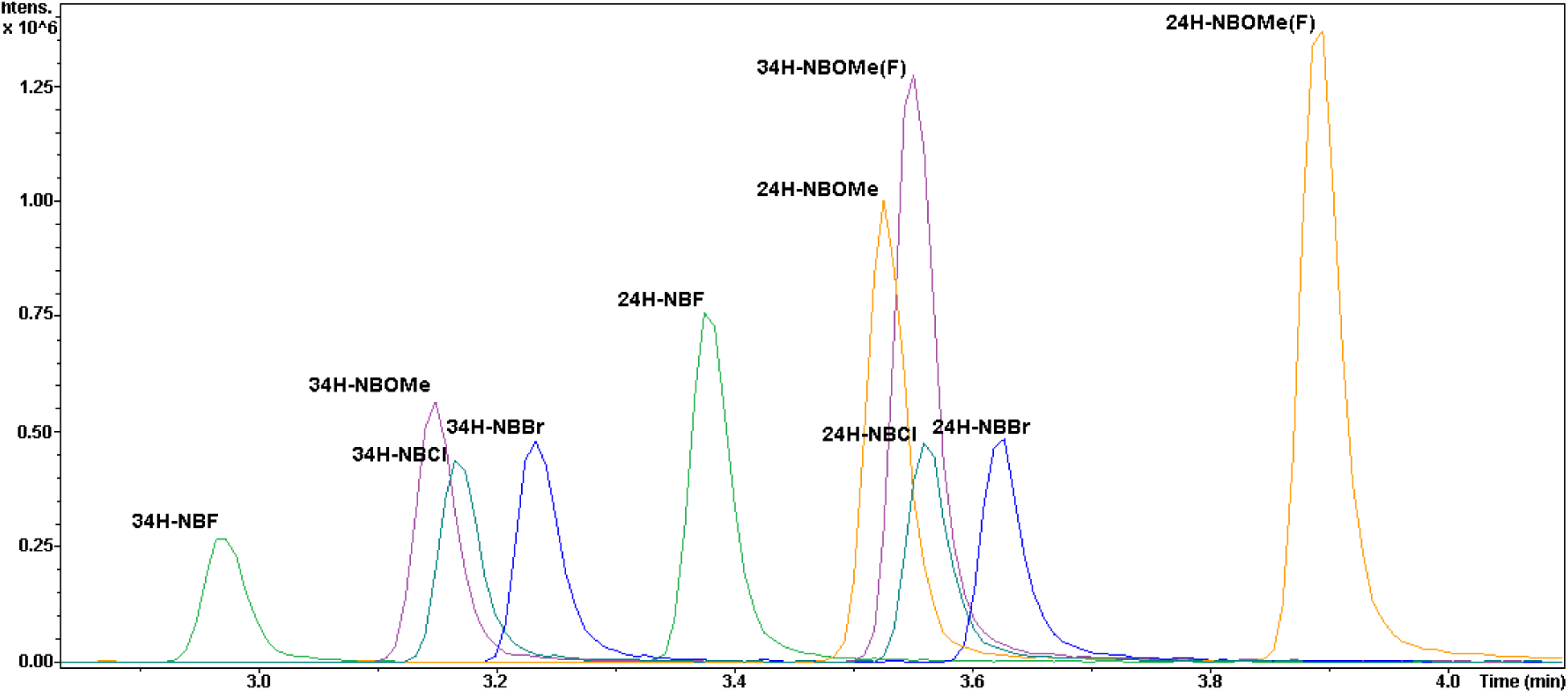
A representative extracted ion chromatogram of mix of all studied compounds (Table 1) using the UHPLC-HRMS method.

**Table 1.**
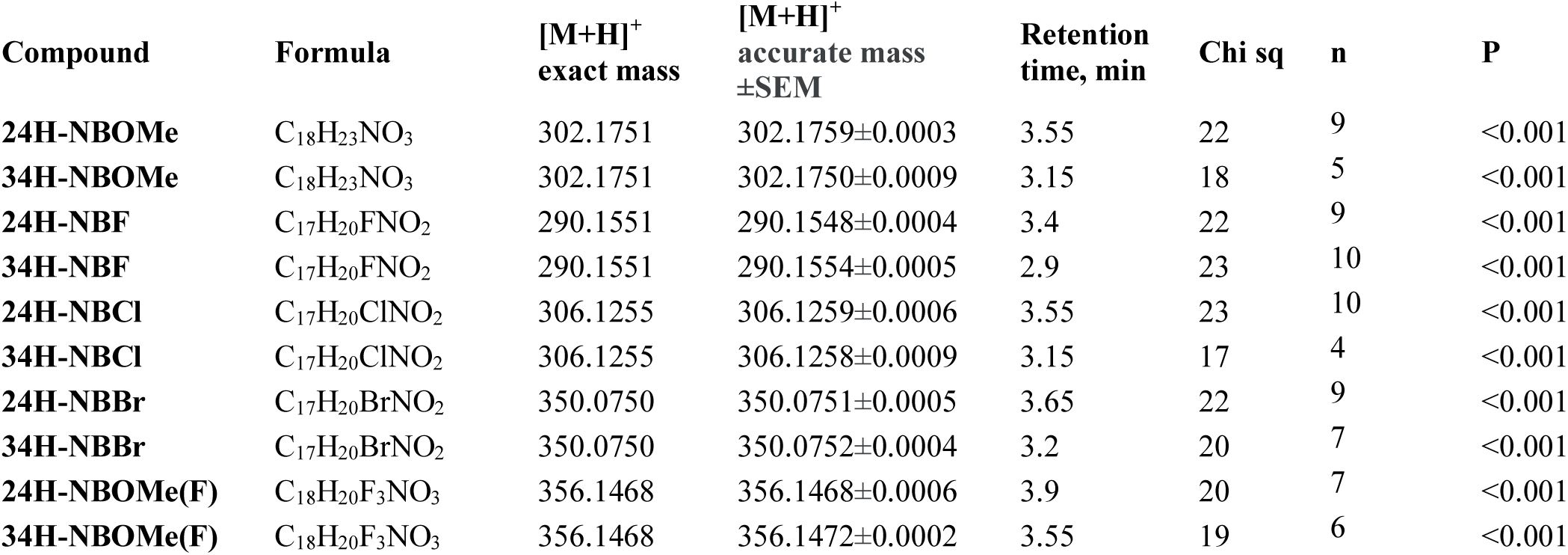
Qualitative analyses of the presence of 10 tested N-Benzyl-2-phenylethylamine (NBPEA) derivatives in zebrafish brain samples, detected by the UHPLC-HRMS (n=4-13; p<0.001 all groups vs. control, Chi-square test) following their acute 20-min exposure to 20 mg/L.

### Artificial intelligence-driven psychopharmacological profiling

In the first experiment, only the initial set of 10 NBPEA-drugs was used to assess the drug and concentration effects compared to the control groups using the artificial intelligence *in silico* analysis of behavioral data with neural networks. All NBPEAs in the concentration of 5 mg/L had little or no effect on the adult zebrafish behavior in the Novel Tank Test. Drugs (Table 2) have formed 4 concentration-dependent effect clusters consisting of: (A) Drugs that does not show any significant effect in any concentration studied (24H-NBOMe, 34H-NBOMe); (B) Drugs that were effective only in concentration of 20 mg/L (24H-NBF, 34H-NBF), (C) Drugs with the similar effects for 10 and 20 mg/L that are different from the control groups (24H-NBCol, 34H-NBCl, 24H-NBBr, 34H-NBBr, 34H-NBOMe(F)); (D) The drug in which 10 and 20 mg/L concentrations were different from both the control group and each other (24H-NBOMe(F)).

**Table 2.** Results of NBPEAs concentrations effects comparisons using artificial-intelligence driven phenotyping approach.

| Drug name | Accuracy baseline | Group |  |  |  |
| --- | --- | --- | --- | --- | --- |
|  |  | Control | 5 mg/L | 10 mg/L | 20 mg/L |
| **24H-NBOMe** | 0.25 | 5 mg/L<br>20 mg/L | 5 mg/L<br>Control | 10 mg/L<br>20 mg/L | 10 mg/L<br>20 mg/L |
| **34H-NBOMe** | 0.26 | 10 mg/L<br>20 mg/L<br>Control | 10 mg/L<br>20 mg/L | 10 mg/L<br>Control | Control |
| **24H-NBF** | 0.26 | 5 mg/L<br>10 mg/L<br>Control | 10 mg/L | 10 mg/L<br>Control | 10 mg/L<br>20 mg/L |
| **34H-NBF** | 0.25 | 10 mg/L<br>Control | 5 mg/L<br>20 mg/L | 5 mg/L<br>Control | 20 mg/L |
| **24H-NBCl** | 0.26 | Control | 10 mg/L<br>Control | 10 mg/L<br>20 mg/L | 10 mg/L<br>20 mg/L |
| **34H-NBCl** | 0.26 | Control | 20 mg/L<br>Control | 10 mg/L<br>20 mg/L | 20 mg/L |
| **24H-NBBR** | 0.26 | Control | Control | 10 mg/L<br>20 mg/L | 10 mg/L<br>20 mg/L |
| **34H-NBBR** | 0.25 | 5 mg/L<br>Control | 10 mg/L<br>Control | 20 mg/L | 20 mg/L |
| **24H-NBOMe(F)** | 0.26 | Control | 5 mg/L<br>10 mg/L<br>Control | 10 mg/L | 20 mg/L |
| **34H-NBOMe(F)** | 0.26 | 5 mg/L<br>Control | Control | 5 mg/L<br>20 mg/L | 20 mg/L |

In the second experiment, the comparison of the 10 NBPEA -drugs to the reference with known psychopharmacological profile were performed. Model training showed overall similar to the experiment 1 results in terms of concentrations effect predictions. The suitable accuracy of prediction of any NBPEA drugs as the reference compound was found only for the 3,4-Methylenedioxymethamphetamine (MDMA), ketamine and Phencyclidine (PCP), but not for lysergic acid diethylamide (LSD) or tetrahydrocannabinol (THC). In the case of LSD, no comparison could be made due to the incompatibility of the behavior of control groups between NBPEAs and LSD data. In the case of THC none of the studied NBPEA compounds were found to be similar to psychopharmacological profile of THC. 24H-NBOMe, 24H-NBF, 24H-NBCl, 34H-NBCl and 34H-NBBr were predicted with higher accuracy than the baseline (0.27) as a ketamine in concentration 40 mg/L (Table 3, Figure 9). 34H-NBOMe, 24H-NBCl, 24H-NBBr, 34H-NBBr, 24H-NBOMe(F) and 34H-NBOMe(F) were predicted with higher accuracy than the baseline (0.28) as MDMA in concentrations 80 and 120 mg/L (Table 3, Figure 9). 24H-NBOMe, 24H-NBF, 24H-NBCl, 34H-NBCl and 34H-NBOMe(F) were predicted with higher accuracy than the baseline (0.51) as PCP (Table 3, Figure 9).

**Figure 9.**
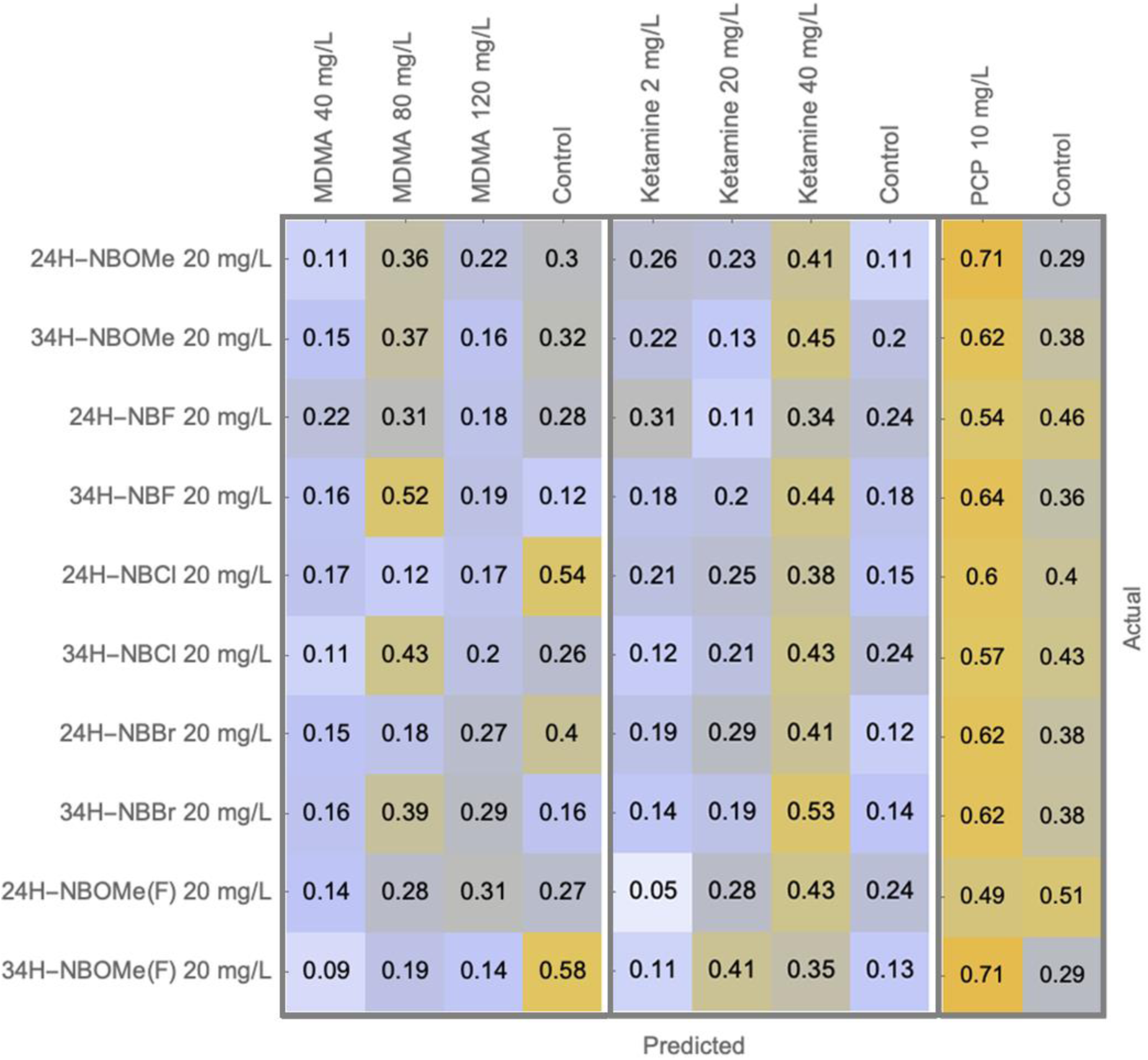
The results of artificial intelligence-driven psychopharmacological profiling investigating similarities between initially studied NBPEA derivatives and MDMA, ketamine or PCP. The data are represented as prediction accuracy/probability. The corresponding prediction accuracy of studied compounds was compared to the baseline prediction accuracy for the reference compounds and if it was higher the results were considered significant. The baseline prediction was 0.28 for MDMA, 0.27 for ketamine and 0.51 for PCP.

**Table 3.** Comparison of the main *in vivo* experiment with reference dataset including MDMA, PCP and ketamine. Bold font – Control group from the initial drug set predicted as a reference control group. Reference data can be used for comparison. Normal font – Control group from the initial drug set predicted as a reference control group and remaining drug groups. Empty cells – Comparison not allowed since control group from the initial drug set was not predicted as a reference control group or since control groups don’t match, thus reference data cannot be used for comparison. NS – no significant changes. No compounds were similar to ketamine 2 or 20 mg/L and MDMA 2 mg/L, thus they were excluded from the table.

| Drug name | Ketamine 40 mg/L | MDMA 80 mg/L | MDMA 120 mg/L | PCP 10 mg/L |
| --- | --- | --- | --- | --- |
| **24H-NBOMe** | Match |  |  | **Match** |
| **34H-NBOMe** |  | Match | NS |  |
| **24H-NBF** | **Match** |  |  | **Match** |
| **34H-NBF** |  |  |  |  |
| **24H-NBCl** | Match | **Match** | NS | **Match** |
| **34H-NBCl** | **Match** |  |  | **Match** |
| **24H-NBBr** |  | Match | Match |  |
| **34H-NBBr** | Match | Match | Match |  |
| **24H-NBOMe(F)** |  | Match | Match |  |
| **34H-NBOMe(F)** |  | Match |  | **Match** |

### In silico molecular modeling

Overall, *in silico* prediction successfully clustered studied compounds in terms of their molecular activities. Both hierarchical clustering and principal component analyses (PCA) support AI-based psychopharmacological phenotyping results (Figures 10-11). All 10 studied NBPEA compounds were well grouped together, forming tight clusters of 2 compounds with similar benzyl but not different phenethylamine substitutions in both analyses. Then, the closest compound to the NBPEAs was ketamine, followed by MDMA. Both THC and LSD were further away, whereas their range was determined mostly by different principal components in PCA. Using hierarchical clustering on activities we also proposed 10 different groups of activities that were differentially affected with different compounds combinations (Figures 10-11). The first is a cluster of activities primarily presented in all drugs except LSD and consisting of seemingly unrelated activities including *acetylcholine neuromuscular blocking agent*, *membrane integrity agonist* and *MAP kinase stimulant*. The second cluster was presented in all NBPEAs, but partially presented in ketamine, MDMA and THC (not LSD), consisting for example of *sodium channel blocker*, *JAK2 expression inhibitor* and *chlordecone reductase inhibitor*. The third cluster was activities represented in all NBPEAs and highly represented in MDMA, but only partially in other reference compounds, consisting of *caspase 3 stimulant*, *nicotinic α6β3β4α5 receptor antagonist*, *spasmolytic*. The next cluster consisted only from *5 hydroxytryptamine release* stimulant activity and was presented in all NBPEAs, both LSD and MDMA, only slightly in THC and not presented in ketamine. The fifth cluster was presented on all NBPEAs, ketamine and MDMA, but only partially in other drugs, *consisting of calcium channel (voltage-sensitive) activator*, *phobic disorders treatment*, *antineurotic*, etc. Other clusters had only sporadic association with NBPEAs. The sixth cluster is associated primarily with THC activity and only partially with MDMA and ketamine, whereas among NBPEAs are mostly methoxy and -Cl substituted compounds. This cluster consist of large number of cytochromes (CYP) related activities, *membrane permeability inhibitor*, *muscle relaxant* and other activities. The seven cluster was represented in MDMA, partially in ketamine and LSD and in methoxy derivatives of NBPEAs, consisting of *leukopoiesis stimulant*, *neurotransmitter uptake inhibitor* and *methylamine-glutamate N-methyltransferase inhibitor*. The eight cluster is associated primarily with ketamine, with slight effect in LSD, consisting of *analgesic*, *glycosylphosphatidylinositol phospholipase D*, and *CYP2A8 substrate* activities. The ninth cluster belongs mostly to LSD activities with only sporadic presentation of other compounds. The most prominent similarity among NBPEAs is existing of relatively low probability of 5 *hydroxytryptamine 7L antagonist* activity, as well as 5 *hydroxytryptamine 2A antagonist* activity for methoxy, F and 24H-NBOMe(F) derivatives. Note that while antagonist serotonergic activity is predicted in NBPEAs, it is similarly predicted in LSD, probably supporting existence of interactions between ligands and receptors, but not their type. Finally, the tenth cluster was associated with LSD, THC, MDMA, and 34H-NBOMe(F) activities, sporadically represented in other compounds and consisting of *nootropic*, *antisecretoric*, *CYP3A*, *CYP3A4*, and *CYP3A5* activities. Overall molecular fingerprints clustering closely resembling functional activity and behavoral analyses results, supporting association of ketamine and PCP with NBPEA-derriatives (Figure 12).

**Figure 10.**
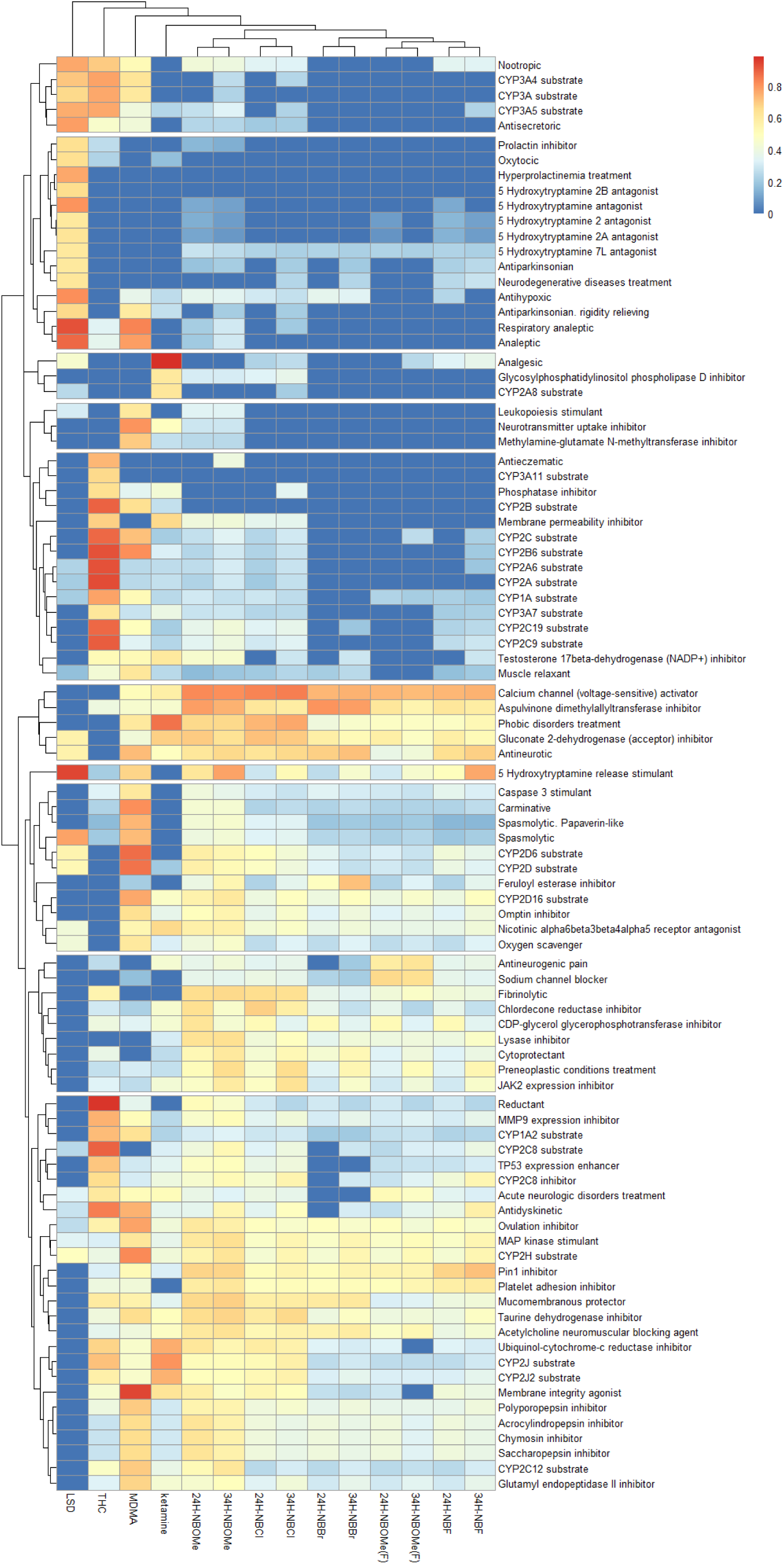
Heatmap representing probability of compounds activities (Pa, %) assessed using the PASSonline software ^126^ with hierarchical clustering.

**Figure 11.**
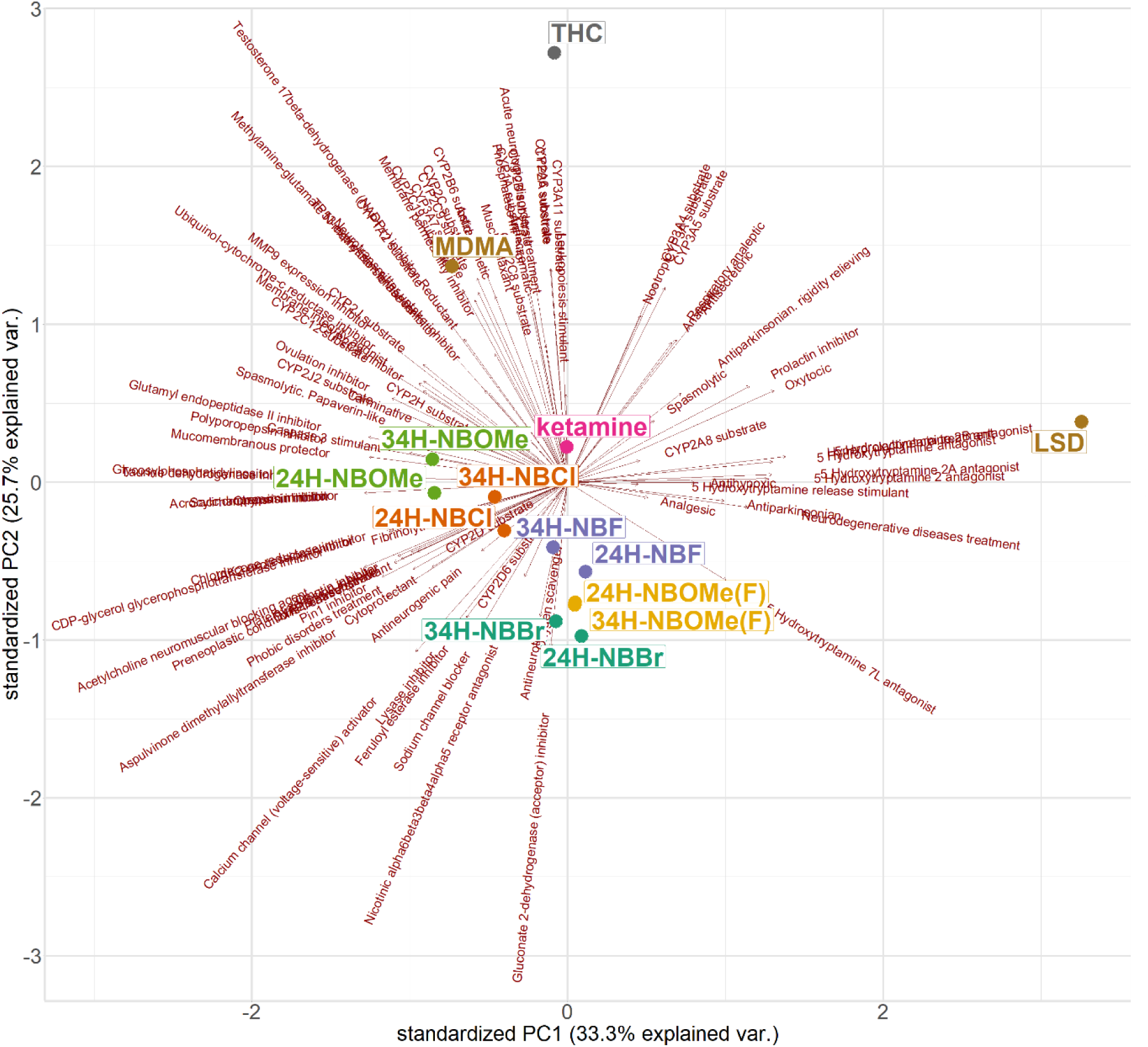
The principal component analysis results represented using the principal components with the most % of explained variance based on the compounds probability of activity. The arrows correspond to an activity loading.

**Figure 12.**
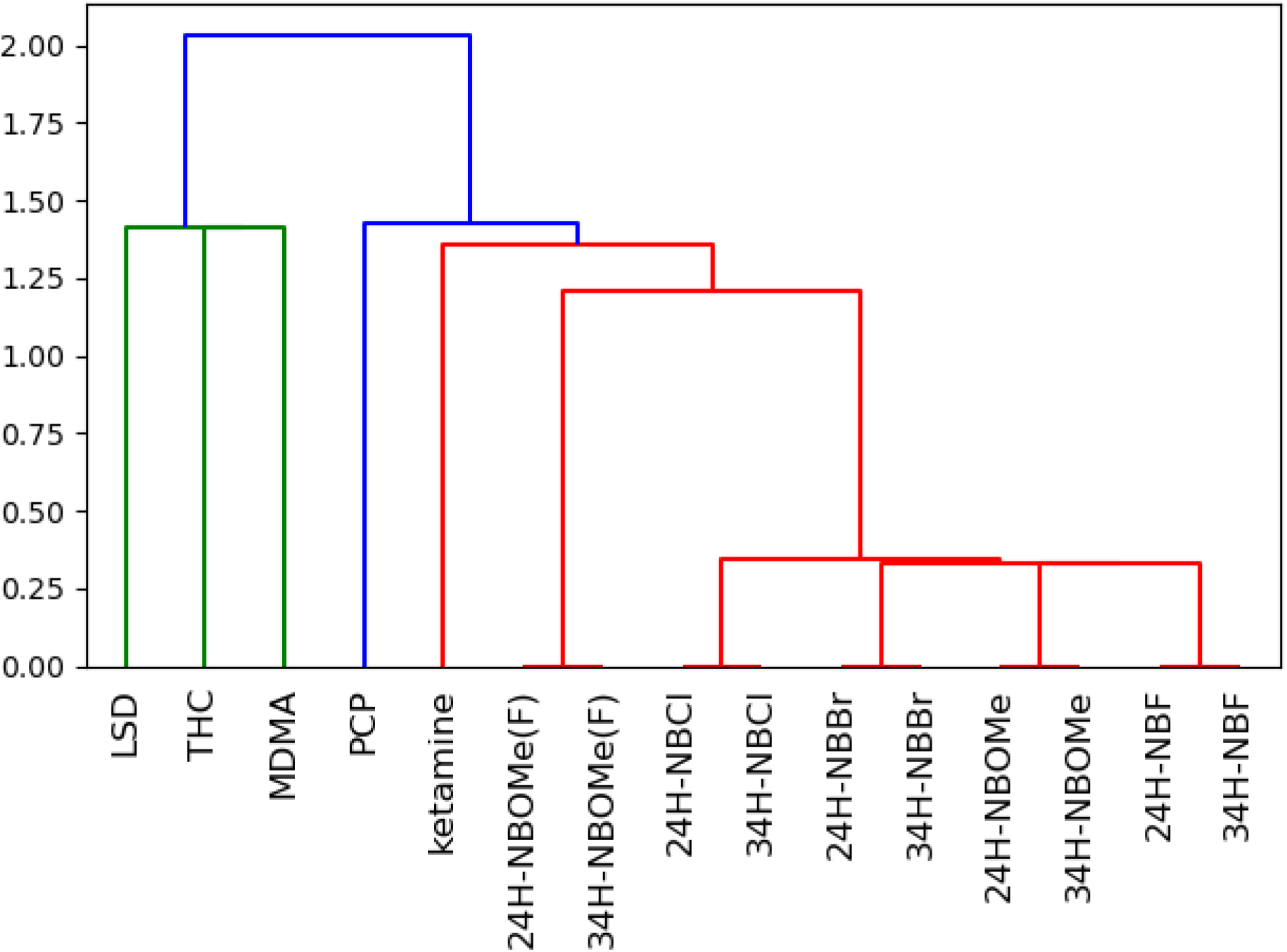
The results of NxN clustering of fingerprints obtained using Open Babel FP4^133^ fingerprinting method, based on 307 key functional substructure SMILES (Simplified Molecular Input Line Entry System) arbitrary target specification (SMARTS) patterns.

## Discussion

This study is the first report utilizing zebrafish for an in-depth, systematical neurobehavioral and neurochemical screening of a battery of novel N-Benzyl-2-phenylethylamines (NBPEAs). To the best of our knowledge, there were only two prior NBOMe studies in zebrafish, both focusing on their toxicity for adult^57^ and larval fish^58^. Here, for the first time, we demonstrate CNS effects of a battery of novel compounds, potentially relevant to antidepressant-, anxiolytic-, hallucinogenic- and psychostimulant-like activity in this model. We also show that a wide range of radical substitutions can dramatically affect behavioral profiles of these neuroactive drugs.

Indeed, mounting biological evidence shows that some *N*-benzyl-substituted phenethylamines can exhibit valuable pharmacological properties and, therefore, may have promising medical applications^59, 60^, including treating various mental disorders ^61^. However, data are lacking on positional isomers of 25X-NBOMes at the position of two methoxy groups in the phenylethyl moiety, especially for substituents different from the methoxy group in the *ortho* position of the phenyl ring of *N*-benzyl fragment. Notably, the addition of a polar methoxy or hydroxyl group to the *ortho* position of the phenyl ring of *N*-benzyl fragment (e.g., for 24H-NBOMe and 24H-NBOH) further increases their affinity for the rat 5-HT_2A_ receptor^62^, raising the possibility that these drugs may exert enhanced neurotropic activity.

Several positional isomers of 25H-NBOMe that differ in the mutual arrangement of two methoxy groups in the phenylethylamine moiety exhibit high affinity for this receptor (assessed *in vitro* for binding at human 5-HT_2A_)^63^. At the same time, their higher potency (EC50) is due to the obligatory presence of one of the two methoxy groups at position 2 of the phenylethylamine moiety, and the presence of a methoxy group at position 3 significantly reduces potency (EC50) to activate this receptor in vitro. In contrast, the 2,4 isomer (24H-NBOMe) showed the highest potency (EC50) and efficacy at 5-HT_2A_ receptors of all tested isomers, and the 3,4 isomer (34H-NBOMe) had lower potency (EC50) with a relatively high efficacy. Similarly, 25H-NBF was reported to have high efficiency, closely resembling LSD and 34H-NBF had slightly reduced efficiency compared to LSD and 25H-NBF ^64^.

Given these *in vitro* data, the present study in zebrafish tested compounds with 2,4- and 3,4-arrangement of two methoxy groups. For the *ortho* position of the phenyl ring of *N*-benzyl moiety, in addition to the traditional methoxy group (NBOMe), we also evaluated -OCF_3_, -F, -Cl and -Br substituents, testing adult zebrafish in the NTT, chosen here as one of the most commonly used tests of zebrafish anxiety and activity ^65–68^, highly sensitive to hallucinogens^52, 54^. This test is based on exposure to novel environment, assessing anxiety-like behavior through diving (the time spent top vs. bottom behavior), locomotor activity (e.g., velocity or distance traveled) and hallucinogenic-like behavior (e.g., horizonal mobility)^52, 54, 65–68^.

Overall, our NTT analyses revealed several common patterns between the compounds, clustered based on their specific profiles (Fig. 6 and Supplementary Table S10) into anxiogenic/hypolocomotor (24H-NBF, 24H-NBOMe and 34H-NBF), behaviorally inert (34H-NBBr, 34H-NBCl and 34H-NBOMe), anxiogenic/hallucinogenic-like (24H-NBBr, 24H-NBCl and 24H-NBOMe(F)), and anxiolytic/hallucinogenic-like (34H-NBOMe(F)) subgroups.

The two-factorial NTT analysis of ring substitutions further shed light on potential mechanisms underlying their behavioral effects (Fig. 5 and Supplementary Tables S4-S8). For example, anxiety-like behavior was affected by substituting in the phenethylamine moiety, showing for 34H-derivatives more anxiolysis than for 24H-derivatives (Fig. 5 and Supplementary Tables S4-S5), whereas velocity was most affected by substitutions in the phenyl ring of *N*-benzyl fragment, since NBOMe(F), NBBr and NBCl all caused hyperlocomotor profile vs. NBF (Fig. 5 and Supplementary Tables S4-S7).

Finally, both substitutions in the phenethylamine moiety and in the *N*-benzyl fragment (but not their interaction) modulate zebrafish hallucinogenic-like behavior, with substitutions in the *N*-benzyl fragment having superior effects compared to substitutions in the phenethylamine moiety (Fig. 5 and Supplementary Tables S4-S6). The 24H-derivatives increased hallucinogenic-like properties compared to 34H-, whereas -NBOMe(F) increased it vs. -NBOMe, -NBF, -NBCl or - NBBr, and -NBBr increased it vs. NBF and NBOMe. Systematically characterizing behavioral effects of multiple chemical substitutions in NBOMe-like psychotropic drugs in zebrafish, such analyses suggest that such novel agents may exert similar behavior/structure-specific psychotropic activity in humans and rodents as well.

To compare obtained results with some well-studied reference compounds, here we performed artificial-driven phenotyping approach recently proposed by our team ^69^. Overall, the study implicates the similarity of studied NBPEA compounds to three classical drugs with hallucinogenic effects – ketamine, PCP and MDMA. However, these compounds belong to different classes of hallucinogenic compounds, that are supposed to act differently in the CNS. Ketamine and PCP are antiglutamergic dissociatives acting primarily through NMDA receptor antagonism whereas MDMA is substituted methylenedioxyphenethylamine belonging to class with primarily empathogen-entactogen effect and acting as a serotonin releasing agent. Unfortunately, the proposed method was unable to compare NBPEAs effects to LSD, due to the low correspondence between corresponding control groups. Whereas no similarities were observed between NBPEAs and THC, as expected.

Overall, the neural networks behavioral analysis showed that NBPEAs differ in their impact. Some drugs show concentration-dependent effects, e.g. concentrations of 10 mg/L do not show considerable effect for 24H-NBOMe, 34H-NBOMe, 24H-NBF, 34H-NBF. For all drugs dosage of 5 mg shows the most inconsiderable effect, and the dosage of 20 mg show the most considerable effect, supporting existence of dose-dependent effects and original behavioral study results. The obtained AI psychopharmacological phenotyping results support hallucinogenic-like nature of multiple compounds. However, the specific similarities are quite surprising giving the high selectivity of 25-NBOMes family to 5HT2A (but not NMDA) receptor and lack of clear MDMA-like activity. Furthermore, while the shuttling behavior observed here partially corresponds to the circling behavior reported for ketamine and PCP in zebrafish earlier ^54, 70, 71^, it is a rather distinct phenotype. Ketamine and PCP induce in zebrafish tight circling behavior with the approximate diameter from 5 cm or 2 fish lengths ^54, 70, 71^, whereas shuttling correspond to the circles with diameter of a whole 20 cm length of the novel tank test apparatus. Similarly, no clear-cut similarities in behavioral phenotypes were observed compared to zebrafish MDMA studies ^72^, since it is mostly associated with anxiolytic effects that were observed only in 34H-NBOMe(F) here using other methods. Furthermore, ketamine (but not PCP) is also primarily anxiolytic in zebrafish ^71^. Overall, 4 types of NBPEA drugs may be considered as a result of AI-phenotyping analysis. Compounds that resemble only NMDA (ketamine or PCP) associated activity – 24H-NBOMe, 24H-NBF and 34H-NBCl, compounds that have only MDMA-like activity – 24H-NBBr, 24H-NBOMe(F), 34H-NBOMe and compounds that have both activities – 24H-NBCl, 34H-NBBr, 34H-NBOMe(F). Note that many drugs that were behaviorally inert in other studies had some similarities with reference compounds here and only 34H-NBF is considered not active in any concentration studied. Overall, if these clusters would be mapped on the Figure 6, they would correspond primarily to velocity or to a lesser extent to shuttling behavior, with NMDA being the slowest and the less shuttling, MDMA the fastest and the combination would be in the middle. However, no clear-cut conclusions can be made from this approach. The AI-based phenotyping used here suggests close association of the studied compounds to different classes of hallucinogenic compounds necessitating further studies.

In addition to testing anxiety and locomotion, we also assessed zebrafish despair-like behavior, for the first time applying ZTI test to novel N-Benzyl-2-phenylethylamine (NBPEA) compounds. Briefly, in this test the zebrafish caudal half was immobilized for 5 min (using the wet sponge) with the rest body freely hanging vertically in a small beaker ^73^. Interestingly, 24H- and 34H-NBOMe(F) increased mobility (distance traveled), as well as serotonin and dopamine metabolism, similarly to acute SSRIs effects in zebrafish^73, 74^. As classical NBPEAs also exhibit relatively potent SERT, DAT and NET-inhibiting properties in vitro^33^, this further implies their potential modulation of monoamine reuptake in the present study (Fig. 4).

Except three neurochemically inert compounds (34H-NBOMe, 34H-NBF and 34H-NBBr), most compounds studied here shared a relatively similar neurochemical profile, increasing serotonin turnover (the 5-HIAA/serotonin ratio for 24H-NBF, 24H-NBBr, 24H-NBOMe(F), 34H-NBCl and 34H-NBOMe(F)) and dopamine turnover (the DOPAC/dopamine ratio, 24H-NBBr, 24H-NBOMe(F), 34H-NBCl and 34H-NBOMe(F)). While only 24H-NBF increased only serotonin turnover, three compounds increased the HVA/dopamine ratio (24H-NBOMe(F), 24H-NBCl and 34H-NBOMe(F)). As altered turnover is commonly seen in rodents and zebrafish treated by various serotonergic and dopaminergic drugs^74–77^, our results generally support serotonergic action of 24H-NBF, 24H-NBBr, 24H-NBOMe(F), 34H-NBCl and 34H-NBOMe(F), and dopaminergic action of 24H-NBBr, 24H-NBOMe(F), 34H-NBCl and 34H-NBOMe(F) in adult zebrafish.

The cheminformatics functional analysis study supports results of AI-based phenotyping highlighting close similarities in functional activity of studied NBPEAs derivatives with ketamine and MDMA, but not LSD and THC. Importantly, the study suggests that compounds have serotonin releasing activity, that may explain increase in serotonin turnover rates observed here. 24H-NBOMe, 34H-NBOMe, 24H-NBOMe(F), 24H-NBF, and 34H-NBF are also suggested to exhibit 5HT2A antagonist activity with low prediction value. However, LSD also has been predicted to have this activity despite 5HT2A agonist activity reported in molecular studies^78^. Probably, such activity should be considered as existence of interaction, but with no strict agonistic/antagonistic relationship. All NBPEAs shown 5HT-7 receptor activity, similarly to LSD, that was also supported by affinity studies (assessed using PDSP Ki database https://pdsp.unc.edu/databases/kidb.php ^79^). Four compounds (24H-NBF, 34H-NBF, 34H-NBBr, 34H-NBCl) also had neurodegenerative diseases treatment activity suggesting their further study in this role. The most pronounced NBPEA associated activity is a cluster of *calcium channel voltage sensitive activator, antineurotic, phobic disorders treatment,* etc. The cluster supports existence of valuable CNS effects since voltage sensitive calcium channels are implicated in psychiatric diseases^80–84^ as well as associated with receptors activity^85, 86^. Finally, no glutamatergic activity was observed in NBPEAs, but 24H-NBCl, 24H-NBCl, 34H-NBOMe, 24H-NBF, and 34H-NBF have probability of analgesic functional activity, that is the most certain predictor for ketamine. Interestingly, both MDMA, ketamine and all NBPEAs have probability of antagonistic activity for nicotinic α6β3β4α5 receptor, suggesting existence of offside effects. Also, some hematopoiesis offside effects are suggested including JAK2 expression inhibition and platelet adhesion inhibition that also should be considered in other studies. Similarly to functional assay, molecular fingerprints supports association of NBPEAs with ketamine and MDMA.

Nevertheless, the present study has several limitations. First, it did not assess lethal concentration and/or potential toxic effects of these drugs. Some NBOMes are highly lethal at high concentration^87, 88^, which can complicate their future application in translational biomedical and preclinical research. At the same time, studied compounds seemed to be rather safe at acute concentrations tested in zebrafish here, since no overt toxic effects (e.g., ataxia, incoordination or seizures) were observed in the present study, hence meriting further cross-species analyses in other models beyond zebrafish. Furthermore, sex and strain differences were also not examined in the present study, albeit being important factors in both clinical and preclinical neurobiological and toxicological assays^89–91^. Since maximum concentration studied here was 20 mg/L, it is also possible that some behaviorally inert compounds are in fact psychoactive in higher concentrations. The study assessed only some behavioral domains (e.g., anxiety, locomotion and despair-like behavior), and further studies are needed to explore full behavioral profiles of these drugs. Finally, while the study utilizes cheminformatics methods, no direct docking to suggested targets was performed, necessitate further studies. However, in fact, we believe that the docking should be performed before compounds synthesis as an investigative approach aimed to target the most affined molecules, whereas this study utilizes another strategy where fine-tuned small changes in derivative suggested to produce distinct (probably activating another pathways or receptors), but not more selective/affined effects on target of reference proteins. Thus, the idea of behavioral screening prior to affinity screening here is beneficial to properly estimate potential key proteins involved in compounds activity for further analysis. We have also performed reverse *in silico* functional analysis that were juxtaposed to *in vivo* studies and allowed better understanding of observed effects and may be used for molecular affinity analyses or pharmacological and physiological investigations. For AI behavioral analysis potential differences in experimental protocols of the present and reference *in vivo* experiments could introduce discrepancies in drugs comparison, e.g., the control group for the 34H-NBF trial did not match with any control group from the reference study. For some trained models, accuracy for control groups matching was below the baseline.

## Conclusions

Assessing neuroactive properties of ten novel synthetic N-Benzyl-2-phenylethylamine (NBPEA) in zebrafish, we showed that all drugs readily crossed the blood-brain barrier, reaching detectable levels in the brain, underlying their behavioral and neurochemical effects observed here. Our analyses also revealed some potential general patterns of structure-activity relationships of these compounds. For example, 2,4-dimethoxy derivatives were more potent in exerting hallucinogenic-like effects on zebrafish behavior, inducing marked changes in serotonin and dopamine turnover. In contrast, most of 3,4-dimethoxy derivatives were behaviorally and neurochemically inert, except for 34H-NBOMe(F), that was highly anxiolytic and hallucinogenic, albeit at concentrations significantly higher than its 2,4-dimethoxy isomer, 24H-NBOMe(F). These findings also fit well with recent pilot *in vitro* data demonstrating higher 5HT_2A_ receptor activation by 2,4- than 3,4-dimethoxylated NBOMe and NBF derivatives^63, 64^.

In addition to positional isomers, some correlations between the structure and activity can be found for various modifications of the *N*-benzyl fragment. For example, among all tested *ortho* substitutions in the phenyl ring, the most prominent behavioral effect was noted for trifluoromethoxy derivatives. Fish that were exposed to 24H- and 34H-NBOMe(F) showed clear hallucinogenic-like phenotype in the NTT, paralleled by markedly increased serotonin and dopamine turnover. In contrast, compounds with classical -OMe substitution in the benzyl ring failed to induce behavioral changes (e.g., 34H-NBOMe) or induced anxiety-like and hypolocomotor phenotype (e.g., 24H-NBOMe) accompanied by unaltered brain monoamine metabolism. Finally, among halogenated compounds, hallucinogenic-like behavioral effects at chosen concentrations were induced only by Cl- and Br-substituted compounds with 2,4-dimethoxylated phenethylamine moiety, whilst their 3,4-dimethoxylated isomers were behaviorally inert. The artificial intelligence-driven phenotyping revealed association of 24H-NBOMe, 24H-NBF and 34H-NBCl with ketamine and PCP behavioral effects, 24H-NBBr, 24H-NBOMe(F) and 34H-NBOMe with MDMA effects and 24H-NBCl, 34H-NBBr, 34H-NBOMe(F) with both MDMA and NMDA antagonists supporting potential hallucinogenic-like phenotype for the studied compounds. Furthermore, *in silico* functional molecular activity modelling also supports existing of similarity between studied NBPEAs drugs, MDMA, and ketamine. Functional analysis implicates potential involvement of serotonin release stimulating activity, calcium channel (voltage-sensitive) activity, some serotonin receptors activity and variety of psychiatric and neurologic disorders treatments activities. Both behavioral (NTT and ZTI) and neurochemical effects induced by trifluoromethoxy substituted derivatives may provide new insights into structure-activity relationships of N-Benzyl-2-phenylethylamines (NBPEAs). Finally, our study strongly supports the growing utility of zebrafish CNS screens for drug development and discovery^92, 93^.

## Methods

### Materials, solvents, and reagents

The following starting materials and reagents were used for the synthesis of compounds **1– 10**: 2,4-dimethoxybenzaldehyde (98%); 3,4-dimethoxybenzaldehyde (99%); 2– fluorobenzaldehyde (97%); 2–chlorobenzaldehyde (97%); 2–bromobenzaldehyde (98%); 2– (trifluoro) methoxybenzaldehyde (96%) all from Alfa Aesar (Kandel, Germany); 2-methoxybenzaldehyde (98%); nitromethane (96%); lithium aluminum hydride (95%); sodium borohydride (99%)—all from Acros Organics, Fair Lawn, NJ, USA; and other reagents: glacial acetic acid (≥99%, Russian GOST 61-75), hydrochloric acid (≥99%, Russian GOST 3118-77), sodium chloride (≥99%, Russian GOST 4233-77), ammonium acetate (≥98.5%, Russian GOST 3117-78), sodium hydroxide (≥99%, Russian GOST 4328-77) all from Reachem (Moscow, Russia), and anhydrous magnesium sulfate (≥99.0%, Bioshop, Canada) purchased commercially and used without further purification. Solvent tetrahydrofuran (99.8%, Tathimprodukt, Kazan, Russia) was dried and purified using standard procedures ^94^. Solvents - methylene chloride (≥99%, TU 2631-019-44493179-98), acetone (≥99%, TU 2633-018-44493179-98), propanol-2 (≥99%, TU 2632-181-44493179-14) all from EKOS-1 (Moscow, Russia); diethyl ether (≥99, TU 2600-001-45286126-11, Medhimprom, Moscow, Russia) and methanol (≥99%, Russian GOST 6995-77, Vekton, St. Petersburg, Russia), were purchased commercially and used without further purification. Flash chromatography was carried out using high-purity grade Merck silica gel (9385, 60 Å, 230-400 mesh).

### NMR spectroscopy

^1^H, ^13^C NMR and ^19^F NMR spectra were acquired in CD_3_OD at 400, 101 and 376 MHz, respectively, using a Avance II spectrometer (Bruker, Switzerland). For ^1^H and ^13^C spectra, signals of residual protons and carbons from the solvent were used as internal standards. To record the ^19^F spectra, trichlorofluoromethane was used as an internal standard. For NMR spectroscopy, CD_3_OD of isotopic purity ≥99.5% (Russian TU 95-669-79, FSUE RSC “Applied Chemistry,” St. Petersburg, Russia) was used.

### Melting point

The melting points of compounds **1–10** were obtained using a Stuart SMP 30 (Bibby Scientific, Ltd., Stone, UK) digital melting temperature measuring instrument at a rate of 1°C/min.

### Gas chromatography–mass spectrometry (GC–MS)

GC-MS analysis was performed using an Agilent 7890B/5977A system (Agilent Technologies, USA) with a HP–5ms (30.0 m × 0.25 mm × 0.25 μm; 19091S-433; Agilent) capillary column. The oven temperature was maintained at 70°C for 1.0 min and then programmed at 15°C/min to 295°C, which was maintained for 15 min. The injector temperature and the interface temperature were 280°C and 290°C, respectively. Helium with flow rate of 1.0 mL/min was used as carrier gas. For control in the formation of compounds **1–10** and their imines **22-31**, 1 μL of the reaction mixture solution was injected into the chromatograph injector without preliminary sample preparation.

### Ultra-high-performance liquid chromatography–high resolution mass spectrometry (UHPLC–HRMS)

High resolution mass spectra of compounds were recorded using a Agilent 1290 Infinity II UHPLC system with a tandem quadrupole time-of-flight (Q-TOF) accurate mass detector Agilent 6545 Q-TOF LC–MS. The Q-TOF instrument was operated with an electrospray ion (ESI) source in positive ion mode. The mobile phase was prepared from 0.1% (vol.) aqueous formic acid (solvent A) and 0.1% (vol.) formic acid in acetonitrile (solvent B), with a linear gradient of 5% to 100% solvent B at 5 minand a flow rate of 0.4 mL/min. UHPLC analysis was performed on a Zorbax Eclipse Plus C18 RRHD (2.1 mm × 50 mm × 1.8 μm) column with additional 5 mm guard column. Acetonitrile (HPLC-gradient grade, Panreac, Barcelona, Spain), water (for GC, HPLC, and spectrophotometry grade, Honeywell, Burdick, and Jackson, Muskegon, USA) and formic acid (≥98.0%) from Sigma-Aldrich (Steinheim, Germany) were used for UHPLC-HRMS. A 5 μg/mL solution of compounds **1–10** in water was prepared.

To examine the ability of synthesized N-Benzyl-2-phenylethylamine (NBPEA) derivatives to enter the brain, we performed their mass spectrometry utilizing brain samples collected from drug-exposed zebrafish, as explained above. Chromatographic separation was undertaken using an Elute UHPLC (Bruker Daltonics GmbH, Bremen, Germany) system, equipped with a Millipore Chromolith Performance/PR-18e, C18 analytical column (Merck, Darmstadt, Germany) with a Chromolith® RP-18 endcapped 5-3 guard cartridges (Merck, Darmstadt, Germany), operated under a flow rate of 0.3 mL/min. The mobile phase was prepared from two solutions, A and B, where A was 0.1% formic acid (v/v) in water and B was 0.1% formic acid (v/v) in acetonitrile (all compounds were purchased at Sigma-Aldrich, St. Louis, MO, US). Elution gradient program for the analysis included a linear gradient of 5% to 95% solvent B at 7 min kept 95% B for 1 min. The initial mobile phase composition was held for 2 min between runs to re-equilibrate the system. The injection volume was 2 μL. Mass spectra were obtained on a Maxis Impact Q-TOF mass spectrometer equipped with an ESI source (Bruker Daltonics GmbH, Bremen, Germany), operated in positive ion mode. Mass calibration was performed using 0.1 M sodium formate in 50:50 water:acetonitrile (vol/vol) solution. Acquisition parameters were electrospray voltage of 4.5 kV; Mass Range from 150 to 1000 m/z; Spectra rate 2 Hz; Save Line and Profile Spectra; End Plate Offset 500 V, Capillary 4500 V, Nebulizer 29 psi, Dry Gas 6.0 L/min and Dry Temperature 220°C.

### Synthetic Procedures

### Synthesis of N-(2-benzylsubstituted)-2-dimethoxyphenylethanamines (1–10)

Products **1, 6** were obtained from their corresponding substituted phenethylamines **15, 16** (1.0 g, 5.52 mmol) and 2-methoxybenzaldehyde (0.864 g, 6.35 mmol), products **2, 7** from substituted phenethylamines **15, 16** (1.0 g, 5.52 mmol) and 2-(trifluoromethoxy)benzaldehyde (1.207 g, 6.35 mmol), products **3, 8** from **15, 16** (1.0 g, 5.52 mml) and 2-fluorobenzaldehyde (0.788 g, 6.35 mmol), products **4, 9** from **15, 16** (1.0 g, 5.52 mmol) and 2-chlorobenzaldehyde (0.893 g, 6.35 mmol), products **5, 10** from **15, 16** (1.0 g, 5.52 mmol) and 2-bromobenzaldehyde (1.175 g, 6.35 mmol).

#### General procedure

A solution of substituted benzaldehyde in CH_2_C_l2_ (40 ml) was added dropwise to a solution of substituted phenethylamine in CH_2_Cl_2_ (40 ml) with cooling to 0°C and stirring for 4-5 h. The completeness of the reaction and identification of the formed imines **22-31** were monitored using GC-MS. The solvent was then evaporated, and the residue dissolved in 80 ml absolute methanol, followed by the addition of NaBH_4_ (0.839 g, 22.08 mmol) over 1 h. After completion, the mix was quenched with distilled H_2_O (20 ml), the organic phase was evaporated, and then 3M aqueous NaOH was added. Products-bases **1-10** were extracted with CH_2_Cl_2_, washed with water and dried. Then 5 ml of propan-2-ol saturated with 5M hydrochloric acid was added to obtain hydrochlorides, which were washed with diethyl ether and dried. Compounds **1-10** were purified by flash chromatography on high-purity grade silica gel (9385, 60 Å, 230-400 mesh) from Merck (Germany), producing hydrochlorides in the form of white powders.

### Analytical data

**24H-NBOMe(F) imine (23).** MS (EI, 70 eV), *m/z* (%): 353 (M^+•^), 151 (100), 121 (26), 175 (18), 152 (11), 91 (11), 77 (10), 78 (7), 202 (6), 109 (6), 353 (5). **24H-NBF imine (24).** MS (EI, 70 eV), *m/z* (%): 287 (M^+•^), 151 (100), 109 (28), 121 (28), 91 (12), 152 (10), 136 (9), 77 (8), 108 (7), 78 (7), 107 (7). **24H-NBCl imine (25).** MS (EI, 70 eV), *m/z* (%): 303 (M^+•^), 151 (100), 121 (25), 125 (18), 152 (16), 91 (11), 89 (9), 77 (8), 127 (6), 78 (6), 90 (4). **24H-NBBr imine (26).** MS (EI, 70 eV), *m/z* (%): 347/349 (M^+•^), 151 (100), 121 (27), 152 (15), 91 (11), 169 (11), 171 (11), 89 (10), 90 (9), 77 (8), 78 (5). **34H-NBOMe(F) imine (28).** MS (EI, 70 eV), *m/z* (%): 353 (M^+•^), 151 (100), 175 (46), 202 (35), 353 (24), 152 (13), 107 (11), 109 (9), 77 (7), 78 (7), 106 (6). **34H-NBF imine (29).** MS (EI, 70 eV), *m/z* (%): 287 (M^+•^), 151 (100), 109 (51), 136 (35), 287 (18), 107 (15), 152 (10), 108 (8), 135 (7), 106 (6), 78 (6). **34H-NBCl imine (30).** MS (EI, 70 eV), *m/z* (%): 303 (M^+•^), 151 (100), 125 (38), 152 (33), 303 (14), 127 (13), 89 (12), 107 (11), 154 (8), 90 (7), 77 (7). **34H-NBBr imine (31).** MS (EI, 70 eV), *m/z* (%): 347/349 (M^+•^), 151 (100), 169 (23), 171 (23), 196 (20), 198 (19), 90 (13), 347 (3), 349 (13), 152 (12), 89 (11). ***N*-(2-Trifluoromethoxybenzyl)-2-(2,4-dimethoxyphenyl)ethanamine hydrochloride (24H-NBOMe(F), (2)).** White powder, yield 94%; ^1^H NMR (CD_3_OD) *δ*, ppm: 2.98-3.05 (m, 2H), 3.21-3.29 (m, 2H), 3.80 (s, 3H), 3.84 (s, 3H), 4.36 (s, 2H), 6.51 (dd, ^3^*J*=8.4 Hz / ^4^*J*=2.4 Hz, 1H), 6.57 (d, ^4^*J*=2.4 Hz, 1H), 7.14 (d, ^3^*J*=8.4 Hz, 1H), 7.43-7.53 (m, 2H), 7.61 (td, ^3^*J*=7.8 Hz / ^4^*J*=1.6 Hz, 1H), 7.74 (dd, ^3^*J*=7.6 Hz / ^4^*J*=2.4 Hz, 1H); ^13^C NMR (CD3OD) *δ*, ppm: 27.94, 46.10, 48.85, 55.89, 55.92, 99.63, 105.98, 117.65, 121.62 (d, ^4^*J_C-F_* = 1.2 Hz), 121.89 (d, ^1^*J_C-F_* = 258.9 Hz), 125.43 (d, ^3^*J_C-F_* = 62.7 Hz), 128.90, 132.03, 132.80, 133.22, 149.13 (d, ^4^*J_C-F_* = 1.0 Hz), 159.88, 162.16; ^19^F NMR (CD3OD) *δ*, ppm: −58.67 (3F); HRMS (ESI, [M+H]^+^), *m/z*: accurate mass 356.1479, exact mass 356.1468, Δ = −2.95ppm, C_18_H_20_F_3_NO_3_; m.p. 92–94°C.

**2-(2,4-Dimethoxyphenyl)-*N*-(2-fluorobenzyl)ethanamine hydrochloride (24H-NBF, (3)).** White powder, yield 72%; ^1^H NMR (CD_3_OD) *δ*, ppm: 2.96-3.04 (m, 2H), 3.19-3.28 (m, 2H), 3.80 (s, 3H), 3.83 (s, 3H), 4.33 (s, 2H), 6.50 (dd, ^3^*J*=8.2 Hz / ^4^*J*=2.4 Hz, 1H), 6.56 (d, ^4^*J*=2.4 Hz, 1H), 7.13 (d, ^3^*J*=8.2 Hz, 1H), 7.22-7.35 (m, 2H), 7.48-7.56 (m, 1H), 7.62 (td, ^3^*J*=7.6 Hz / ^4^*J*=1.6 Hz, 1H); ^13^C NMR (CD_3_OD) *δ*, ppm: 27.91, 45.45 (d, ^3^*J_C-F_* = 4.0 Hz), 48.67, 55.90, 55.93, 99.62, 105.96, 116.95 (d, ^2^*J_C-F_* = 21.6 Hz), 117.73, 119.85 (d, ^2^*JC_-F_* = 14.8 Hz), 126.21 (d, ^3^*J_C-F_* = 3.7 Hz), 132.04, 133.27 (d, ^3^*J_C-F_* = 8.4 Hz), 133.42 (d, ^4^*J_C-F_* = 2.8 Hz), 159.90, 162.13, 162.72 (d, ^1^*J_C-F_* = 247.8 Hz); ^19^F NMR (CD3OD) *δ*, ppm: -118.31 (1F); HRMS (ESI, [M+H]^+^), *m/z*: accurate mass 290.1553, exact mass 290.1551, Δ = −0.79 ppm, C_17_H_20_FNO_2_; m.p. 123–124°C.

***N*-(2-Chlorobenzyl)-2-(2,4-dimethoxyphenyl)ethanamine hydrochloride (24H-NBCl, (4)).** White powder, yield 92%; ^1^H NMR (CD_3_OD) *δ*, ppm: 2.98-3.06 (m, 2H), 3.23-3.31 (m, 2H), 3.80 (s, 3H), 3.84 (s, 3H), 4.42 (s, 2H), 6.50 (dd, ^3^*J*=8.0 Hz / ^4^*J*=2.4 Hz, 1H), 6.56 (d, ^4^*J*=2.4 Hz, 1H), 7.15 (d, ^3^*J*=8.4 Hz, 1H), 7.41-7.51 (m, 2H), 7.56 (dd, ^3^*J*=7.6 Hz / ^4^*J*=2.0 Hz, 1H), 7.67 (dd, ^3^*J*=7.2 Hz / ^4^*J*=2.0 Hz, 1H); ^13^C NMR (CD3OD) *δ*, ppm: 27.93, 48.89, 49.26; 55.92, 55.97, 99.64, 105.99, 117.70, 129.03, 130.63, 131.22, 132.08, 132.56, 133.25, 135.83, 159.89, 162.14; HRMS (ESI, [M+H]^+^), *m/z*: accurate mass 306.1260, exact mass 306.1255, Δ = −1.43 ppm, C_17_H_20_ClNO_2_; elemental analysis calculated (%) for C_17_H_20_ClNO_2_·HCl: C 59.66, H 6.18, Cl 20.72, N 4.09; found: C 58.57, H 5.99, Cl 20.16, N 3.96; m.p. 105–107°C.

***N*-(2-Bromobenzyl)-2-(2,4-dimethoxyphenyl)ethanamine hydrochloride (24H-NBBr, (5)).** White powder, yield 94%; ^1^H NMR (CD_3_OD) *δ*, ppm: 2.98-3.08 (m, 2H), 3.26-3.32 (m, 2H), 3.80 (s, 3H), 3.84 (s, 3H), 4.42 (s, 2H), 6.51 (dd, ^3^*J*=8.0 Hz / ^4^*J*=2.4 Hz, 1H), 6.57 (d, ^4^*J*=2.0 Hz, 1H), 7.15 (d, ^3^*J*=8.4 Hz, 1H), 7.39 (td, ^3^*J*=7.4 Hz / ^4^*J*=1.6 Hz, 1), 7.50 (td, ^3^*J*=7.4 Hz / ^4^*J*=1.2 Hz, 1H), 7.67 (dd, ^3^*J*=7.6 Hz / ^4^*J*=2.4 Hz, 1H), 7.73 (dd, ^3^*J*=8.0 Hz / ^4^*J*=1.2 Hz, 1H); ^13^C NMR (CD_3_OD) *δ*, ppm: 27.96, 48.94, 51.77; 55.92, 55.99, 99.65, 106.00, 117.69, 125.93, 129.64, 132.09, 132.43, 132.67, 133.12, 134.63, 159.89, 162.15; HRMS (ESI, [M+H]^+^), *m/z*: accurate mass 350.0756/352.0739, exact mass 350.0750/352.0731, Δ = −1.76/−2.11 ppm, C_17_H_20_BrNO_2_; elemental analysis calculated (%) for C_17_H_20_BrNO_2_·HCl: C 52.80, H 5.47, Br 20.66, Cl 9.17, N 3.62; found: C 52.72, H 5.39, Br 19.99, Cl 8.88, N 3.50; m.p. 125–126°C.

***N*-(2-Trifluoromethoxybenzyl)-2-(3,4-dimethoxyphenyl)ethanamine hydrochloride (34H-NBOMe(F), (7)).** White powder, yield 77%; ^1^H NMR (CD_3_OD) *δ*, ppm: ^1^H NMR (CD_3_OD) *δ*, ppm: 3.01-3.08 (m, 2H), 3.29-3.38 (m, 2H), 3.82 (s, 3H), 3.85 (s, 3H), 4.37 (s, 2H), 6.86 (dd, ^3^*J*=8.0 Hz / ^4^*J*=2.0 Hz, 1H), 6.90-6.97 (m, 2H), 7.43-7.53 (m, 2H), 7.57-7.66 (m, 1H), 7.77 (dd, ^3^*J*=7.6 Hz / ^4^*J*=1.6 Hz, 1H); ^13^C NMR (CD_3_OD) *δ*, ppm: 32.79, 46.26, 50.32; 56.58 (2C), 113.49, 113.75, 121.62 (d, ^4^*J_C-F_* = 1.3 Hz), 121.90 (d, ^1^*J_C-F_* = 258.8 Hz), 122.23, 125.46 (d, ^3^*J_C-F_* = 57.7 Hz), 128.90, 130.46, 132.81, 133.31, 149.11 (d, ^4^*J_C-F_* = 1.2 Hz), 149.82, 150.86; ^19^F NMR (CD_3_OD) *δ*, ppm: −58.69 (3F); HRMS (ESI, [M+H]^+^), *m/z*: accurate mass 356.1476, exact mass 356.1468, Δ = −2.23 ppm, C_18_H_20_F_3_NO_3_; m.p. 78–80°C.

***N*-(2-Fluorobenzyl)-2-(3,4-dimethoxyphenyl)ethanamine hydrochloride (34H-NBF, (8)).** White powder, yield 82%; ^1^H NMR (CD_3_OD) *δ*, ppm: 2.99-3.09 (m, 2H), 3.28-3.38 (m, 2H), 3.82 (s, 3H), 3.85 (s, 3H), 4.35 (s, 2H), 6.86 (d, ^3^*J*=8.0 Hz, 1H), 6.89-6.97 (m, 2H), 7.22-7.34 (m, 2H), 7.48-7.57 (m, 1H), 7.65 (td, ^3^*J*=7.6 Hz / ^4^*J*=1.6 Hz, 1H); ^13^C NMR (CD3OD) *δ*, ppm: 32.76, 45.59 (d, ^3^*J_C-F_* = 4.0 Hz), 50.13; 56.60 (2C), 113.50, 113.80, 116.96 (d, ^2^*J_C-F_* = 21.5 Hz), 119.84 (d, ^2^*J_C-F_* = 14.8 Hz), 122.25, 126.23 (d, ^3^*J_C-F_* = 3.7 Hz), 130.50, 133.32 (d, ^3^*J_C-F_* = 8.4 Hz), 133.50 (d, ^4^*J_C-F_* = 2.7 Hz), 149.79, 150.84, 162.71 (d, ^1^*J_C-F_* = 247.7 Hz); ^19^F NMR (CD_3_OD) *δ*, ppm: -118.04 (1F); HRMS (ESI, [M+H]^+^), *m/z*: accurate mass 290.1553, exact mass 290.1551, Δ = −0.87 ppm, C_17_H_20_FNO_2_; m.p. 163–165°C.

***N*-(2-Chlorobenzyl)-2-(3,4-dimethoxyphenyl)ethanamine hydrochloride (34H-NBCl, (9)).** White powder, yield 89%; ^1^H NMR (CD_3_OD) *δ*, ppm: 3.02-3.10 (m, 2H), 3.31-3.41 (m, 2H), 3.82 (s, 3H), 3.85 (s, 3H), 4.44 (s, 2H), 6.87 (d, ^3^*J*=8.4 Hz, 1H), 6.90-6.98 (m, 2H), 7.41-7.51 (m, 2H), 7.53-7.59 (m, 1H), 7.70 (dd, ^3^*J*=7.0 Hz / ^4^*J*=1.6 Hz, 1H); ^13^C NMR (CD_3_OD) *δ*, ppm: 32.75, 49.42, 50.33; 56.62 (2C), 113.52, 113.82, 122.27, 129.04, 130.48, 130.62, 131.23, 132.61, 133.35, 135.84, 149.82, 150.86; HRMS (ESI, [M+H]^+^), *m/z*: accurate mass 306.1259, exact mass 306.1255, Δ = −1.06 ppm, C_17_H_20_ClNO_2_; elemental analysis calculated (%) for C_17_H_20_ClNO_2_·HCl: C 59.66, H 6.18, Cl 20.72, N 4.09; found: C 58.87, H 6.12, Cl 20.35, N 3.88; m.p. 166–168°C.

***N*-(2-Bromobenzyl)-2-(3,4-dimethoxyphenyl)ethanamine hydrochloride (34H-NBBr, (10)).** White powder, yield 91%; ^1^H NMR (CD3OD) *δ*, ppm: 3.03-3.10 (m, 2H), 3.34-3.42 (m, 2H), 3.82 (s, 3H), 3.85 (s, 3H), 4.44 (s, 2H), 6.87 (dd, ^3^*J*=8.0 Hz / ^4^*J*=1.6 Hz, 1H), 6.90-6.98 (m, 2H), 7.39 (td, ^3^*J*=7.8 Hz / ^4^*J*=1.6 Hz, 1H), 7.49 (td, ^3^*J*=7.4 Hz / ^4^*J*=0.8 Hz, 1H), 7.72 (td, ^3^*J*=8.0 Hz / ^4^*J*=1.2 Hz, 2H); ^13^C NMR (CD_3_OD) *δ*, ppm: 32.77, 50.36, 51.92, 56.63 (2C), 113.53, 113.83, 122.29, 125.94, 129.65, 130.47, 132.41, 132.72, 133.24, 134.63, 149.82, 150.86; HRMS (ESI, [M+H]^+^), *m/z*: accurate mass 350.0755/352.0737, exact mass 350.0750/352.0731, Δ =−1.38/−1.46 ppm, C_17_H_20_BrNO_2_; elemental analysis calculated (%) for C_17_H_20_BrNO_2_·HCl: C 52.80, H 5.47, Br 20.66, Cl 9.17, N 3.62; found: C 52.06, H 5.19, Br 19.92, Cl 8.86, N 3.58; m.p. 163–165°C.

### Animals and housing

Adult, 3-4 months old wild-type short-fin experimentally naïve zebrafish (approximately 1:1 sex ratio) were obtained from a local distributor (Axolotl, Ltd., St. Petersburg, Russia) and housed for at least 2 weeks in standard conditions in groups of 10-15 fish in 4-L tanks (2.5-3.75 fish/L) at the Aquatic Facility of the Almazov National Medical Research Center (St. Petersburg, Russia) using the ZebTec Active Blue Stands with Water Treatment Unit (Tecniplast, West Chester, USA), with filtered water environment maintained at 27±0.5°C and pH 7.4. The illumination (950–960 lux) in the home tanks was provided by 18-W fluorescent light tubes with a 12:12 light:dark cycle. All fish were fed with small food pellets Neon Micro Granules for fish size 1-2 cm long (Dajana Pet, Bohuňovice, Czech Republic) twice a day according to the zebrafish feeding standards^95^.

All fish belonged to the same baseline population. All animals tested were included in final analyses, without removing outliers. All experiments were performed as planned, and all analyses and endpoints assessed were included without omission. Animal experiments were approved by the Institutional IACUC and fully adhered to National and Institutional guidelines and regulations. The study experimental design and its description here, as well as data analysis and presenting, adhered to the ARRIVE (Animal Research: Reporting of In Vivo Experiments) guidelines for reporting animal research and the PREPARE (Planning Research and Experimental Procedures on Animals: Recommendations for Excellence) guidelines for planning animal research and testing, see Ethical Confirmation statement for approval and ethical details of animals use in the research. All animals were allocated to the experimental groups randomly using a random number generator (https://www.random.org/).

The outbred short-fin strain selection for the present study was based on population validity considerations and their relevance for the present study^96^. Specifically, while genetically controlled inbred zebrafish strains may offer reproducible and more reliable systems for neurogenetics research, modeling CNS disorders (such as in the present study) involves mimicking ‘real’ human maladies that affect genetically heterogenous clinical populations. Thus, using outbred populations of zebrafish can represent a more populationally valid and translationally relevant approach for the purpose of this study. This selection also considered overt strain-specific peculiarities of zebrafish behaviors in different^97^ tests that can be mitigated by using wild-type outbred fish, and also paralleled recent rodent findings (noting no higher phenotypic trait variability in outbred (vs. inbred) mice and concluding that outbred strains may be better subjects for most biomedical experiments^98^).

### Behavioral testing

Behavioral testing of parallel zebrafish cohorts was performed on different days with individual control group for each compound. Behavioral analyses were performed between 11:00 am and 2:00 pm by individually exposing zebrafish to the NTT. Prior to behavioral testing, all fish (n=16-17) were transported from the holding room and acclimated to the testing room for 2 h. After this acclimation period, the fish were individually pre-exposed to 5, 10 or 20 mg/L of studied compounds or to drug-free water (control group) for 20 min. The concentrations were chosen based on our pilot studies with a wide concentration range of tested compounds, with a start concentration based on human recreational 25H-NBOMe doses, further increasing the concentration until most of compounds shown psychoactive effect. After behavioral testing, fish were returned to their respective home tanks in the aquatic housing system.

The NTT was chosen here as the most commonly used behavioral test that is sensitive to alterations in anxiety and locomotion in zebrafish^99, 100^, and was performed similarly to^101, 102^. The NTT apparatus consisted of a 2-L acrylic rectangular tank (20 height × 20 length × 5 width cm) filled with water up to 19 cm, and divided into two equal virtual horizontal portions, corresponding to top and bottom halves of the tank ^74, 77, 96, 102–104^, and two vertical portions, corresponding to left and right halves. Back and lateral sides of the tank were covered with nontransparent white covers (fixed to the outside walls), to increase contrast and reduce external visual clues during behavioral recording ^74, 77, 105, 106^. Trials were video-recorded using an SJ4000 action camera (SJCAM, Ltd., Shenzhen, China) at 60 frames/s ^105, 106^. We assessed the mean velocity (cm/s), the time spent in top (s), the number of top entries, the latency to enter top (s), and the horizontal shuttling behavior based on the center body position computation, using the EthoVision XT11.5 software (Noldus IT, Wageningen, Netherlands), as in^107, 108^. The horizontal ‘shuttling’ behavior was calculated as the normalized horizontal transitions (total left-to-right or right-to-left horizontal transitions) divided by distance traveled, to reflect specific drug-induced hallucinogenic-like activity, closely resembling circling behavior observed in zebrafish earlier ^70, 72^. The mean velocity in the test is associated with locomotor activity, similarly to the rodent tests, whereas top dwelling behavior corresponds to anxiolytic like behavior (the more time fish spent top, the less anxious it is) and is supported by the large amount of pharmacological, neurochemical, genetic and physiological studies^93, 108–110^.

Furthermore, to study effects of individual substitutions between all compounds we used Generalized Linear Models (GZLM) two-factorial analysis (see Statistical analysis and data handling for details), using substitutions in the phenethylamine moiety (24H and 34H) and *N*-benzyl fragment (-OMe, -F, -Cl, -Br, -OMe(F)) rings and their interactions as factors. The experiment was conducted similarly to the original NTT study, using 20 mg/L concentration for each compound (n=11-12) and done in parallel in one day to ensure lack of daytime effect. The control drug-free group was also done in parallel with other groups for the mean reference, but excluded from further statistical analysis to perform proper two-factorial analysis.

Since 24H- and 34H-NBOMe(F) had pronounced anxiolytic effect in the NTT (see results, Fig. 2 and Supplementary Table S1), it became logical to their potential antidepressant properties. Recently, we proposed the ZTI test as a tool to assess despair-like behavior in zebrafish, analogous to testing it in rodents^73^. The test is sensitive to antidepressant drugs and common stressors, but not to anxiolytic treatments, thus making it (unlike NTT) an efficient tool to distinguish anxiolytic vs. antidepressant activity^73^. Briefly, the fish caudal half was immobilized for 5 min using the wet viscose sponge (8 length x 4 height x 5 width, cm) cut in the middle with a sharp scalpel to a 2-cm depth from the bottom and attached to the top of the beaker using two additional 2-cm cuts of the sponge on the sides, to allow fixation by the beaker walls, with the cranial part of the fish body remaining freely hanging vertically in a small beaker, a 7×5.5-4.8 cm transparent plastic cup shaped as a truncated cone, filled with water (similar to ^73^). In the ZTI increase in activity corresponds to antidepressant-like activity and decrease in despair-like behavior, similarly to rodent tests. Fish were pre-exposed to 5 mg/L of 24H-NBOMe(F) (n=13) or 20 mg/L 34H-NBOMe(F) (n=15) water solution or drug-free water in the 0.5L beaker for 20 minutes, similar to the NTT study. The concentration was chosen as having most pronounced anxiolytic effects in the NTT. The distance travelled by the head (cm) and time spent mobile (s) were studied using the EthoVision XT11.5 software (Noldus IT, Wageningen, Netherlands).

### Brain neurochemistry

Since brain monoaminergic systems play a key role in NBOMe CNS effects^31–37^, here we studied changes in zebrafish whole-brain concentrations of norepinephrine (NE), serotonin (5-HT), dopamine (DA) and their metabolites 5-hydroxyindoleacetic acid (5-HIAA), 3,4-dihydroxyphenylacetic acid (DOPAC) and homovanillic acid (HVA) using the high-performance liquid chromatography (HPLC), similarly to ^73, 74, 77, 96, 102, 103^. As in the NTT and ZTI studies, fish (n=7-10) were pre-exposed to 20 mg/L of 24H-NBF, 34H-NBOMe, 34H-NBF, 34H-NBCl, 34H-NBBr, 34H-NBOMe, or to 10 mg/L 24H-NBBr, 24H-NBOMe(F) or drug-free water, in the 0.5L beaker for 20 min. The concentrations were chosen based as most behaviorally active in zebrafish tests. 24H-NBOMe and 24H-NBCl were excluded from neurochemical analysis due to pronounced anxiogenic effects in both NTT concentration-dependent and two-factorial studies, that reduces the potential of their use in clinics.

The fish were euthanized in ice-cold water immediately after drug exposure, decapitated following the cessation of opercular movements, and their brains dissected on ice and stored in liquid nitrogen for analyses, as in^73, 74, 77, 96, 102, 103^. On day of analyses, all samples were weighted and placed into 10 μL of ice-cold 0.1 M perchloric acid (Sigma Aldrich, St. Louis, MO, USA) solution with 100 ng/mL 3,4-dihydroxybenzylamin (DHBA, internal standard) per 1 mg of brain tissue for preservation of analytes in line with other studies^73, 74, 77, 96, 102, 103^. The samples were next sonicated for 10 s at half-power settings, cleared by centrifugation and filtered through a 0.22-μm Durapore-PVDF centrifuge filter (Merck Millipore, Billerica, MA, USA). HPLC was performed using a CA-5ODS column and a HTEC-500 chromatograph (Eicom, San Diego, CA, USA) with a carbon WE-3 G electrode using a +650 mV applied potential, similar to^73, 74, 77, 96, 102, 103^. Chromatography mobile phase consisted of 0.1 M phosphate buffer, 400 mg/L sodium octylsulphonate, 50 mg/L ethylenediaminetetraacetic acid (EDTA), 17-% methanol and was adjusted to pH 4.5 by phosphoric acid (all reagents were purchased from Sigma Aldrich, St. Louis, MO, USA), in line with ^73, 74, 77, 96, 102, 103^. The concentrations data were normalized using individual DHBA sample concentrations, and presented as pg/mg of brain tissue weight, similar to^73, 74, 77, 96, 102, 103^. Additionally, we also assessed monoamine turnover in the brain using the ratios of 5-HIAA to serotonin, DOPAC to dopamine, and HVA to dopamine.

### Statistical analyses and data handling

Statistical analysis was performed using the Statistics 10, R programming language and Excel software package. Behavioral data were analyzed using the Kruskal-Wallis (KW) test followed with Dunn’s post-hoc test (for significant KW data) for pairwise between group comparison. The sample size was chosen here based on previously published studies on zebrafish stress-related behavior, including own works^47, 73, 74, 111, 112^ and sample size estimation using the R package pwr2. Briefly, we used ss.1way function to determine the sample size, using 7 groups, 0.05 alpha level, 0.85 power level (0.15 type II error probability) and 0.4 effect size (based on out previous works^47, 73, 74, 111, 112^ and efficiency assumptions from pilot studies). As a result, we had n=15 for each group and used n=15-17 depending on the experiment. Mann-Whitney U-test was used to compare zebrafish behavior in experiments involving Zebrafish Tail Immobilization test.

The present study utilized Generalized Linear Models (GZLMs) to analyze effects of different substitutions in phenyl rings of synthesized N-Benzyl-2-phenylethylamine (NBPEA) on zebrafish behavior in the Novel Tank Test. GZLM is an effective method to analyze data with non-normal distribution since it is a generalization of regression methods that allows variables to have distributions other than a normal distribution ^113^. GZLM method is widely used in various fields^114–,116^, including zebrafish neuroscientific and neurobehavioral studies ^102, 117^. For behavioral and neurochemical analysis, we performed the Wald chi-square (χ2) analysis of variance (ANOVA, Type II; Fig. 5 and Supplementary Table S5) of GZLM (Supplementary Table S4), fits, followed by Tukey’s post-hoc testing for significant GZLM/Wald pair-wise comparison data, similarly to ^102^ (Fig. 5 and Supplementary Tables S6-8). GZLM is an effective method to analyze multifactorial and factors interaction data that provides robust and sensitive results both for most types of data ^114, 115, 118, 119^. To assess effects of substitutions on zebrafish behavior, substitutions in the phenethylamine moiety (24H or 34H), *N*-benzyl fragment (NBOMe, NBF, NBCl, NBBr and NBOMe(F)) and their interaction effects were used as predictors comparing all 10 derivatives. To choose optimal GZLM distribution and link functions (goodness of fit) for each endpoint, we compared (where applicable) the Akaike information criterion (AIC) levels ^120, 121^ of Gaussian distribution (identity link), Poisson distribution (with log link), Gamma distribution (inverse and log links) and Inverse Gaussian distribution (with inverse and log links), choosing the least AIC score (indicating the model most likely to be correct) ^122^. K-means clustering analyses (4 centers) was performed on standardized means of velocity, time spent top and horizontal alterations relative to velocity of all groups (including control reference group) in GZLM experiment (Fig. 7, Supplementary Table S10). Correlation analyses between horizontal (left-right) alterations and velocity or horizontal alterations relative to velocity vs. velocity in control groups among all NTT experiments were performed using Spearman rank correlation coefficient (Supplementary Table S2). All calculations were performed using the R software (R Core Team, New Zealand) ^123^.

Mass-spectrometry data were analyzed using DataAnalysis® and TASQ® Software (Bruker Daltonics GmbH, Bremen, Germany). Peaks with area higher than 400 000 units were considered as compound presence in a probe for further qualitative analysis using Chi-square, comparing the number of probes with studied compounds in experimental groups vs. control (the number of positive probes in control were 0 for any compound studied; Table 1). All experiments here were performed by blinded experimenters, using individual codes for fish/groups identification. Unless specified otherwise, all data is expressed as mean ± standard error of mean (SEM), and P<0.05 in all analyses.

### Artificial intelligence-driven psychopharmacological profiling

Finally, in addition to the classical statistical analyses aimed to access strict behavioral alterations, we also performed a machine learning approach for drugs profiling aimed to compare the results with other studied in zebrafish compounds. For these purposes we used the convolutional neural network (CNN) with ResNet34 architecture that were recently developed by our team ^69^. As was shown this approach is capable to fairly-well distinguish and cluster effects of different psychoactive substances in the zebrafish, including different hallucinogenic compounds such as ketamine, phencyclidine (PCP) and lysergic acid diethylamide (LSD), using only behavioral data ^69^. Here we trained the neural network on the raw fish coordinates tracks data in behavioral experiment assessed using Noldus Ethovision XT 11.5 similarly to other behavioral studies. Then each individual fish track was cut into pieces of 30 seconds length each and used for the training that was shown to be the most sensitive approach in the previous study ^69^. Briefly, the data was further processed using Python matplotlib library 3.5.2 ^124^ to generate track images with color corresponding to the fish velocity and using data augmentation procedure as in ^69^ to increase the size of training datasets.

We designed two different *in silico* experiments. The first aimed to compare effects in the initial 10 NBPEA-drugs (20 mg/L) to support the main findings of the article and the model validity. Here we studied how well different drugs, their concentrations and control groups can be distinguished and predicted. This data corresponding to the potential effect of this factors on the zebrafish behavior. The results were further validated using the same experiment on the data from the second study (with the same initial drugs).

The second study performed comparative analysis of the initial results with previously published by our group results, using LSD, ketamine, PCP, 3,4-methylenedioxymethamphetamine (MDMA) and tetrahydrocannabinol (THC) ^52, 54, 71, 72, 125^. The drugs were chosen based on the quality of prediction in the original article ^69^, the pilot *in silico* studies and the psychopharmacological profile observed in the main behavioral study. Here we trained neural networks on the old compounds data and used these neural networks on the initial set of drugs to find similarities. Many of these old compounds represent classical hallucinogens, thus providing valuable additional reference for behavioral estimation of hallucinogenic potential. The results of artificial intelligence analysis are represented as prediction accuracy/probability, unless specified otherwise. The corresponding prediction accuracy of studied compounds was compared to the baseline prediction accuracy for the reference compound and if it was higher the results were considered significant.

### In silico molecular modeling

To better understand observed behavioral effects, here we used predictive analyses of structure-activity relationship of the novel compounds using the PASSonline software ^126^. The PASSonline trained on Multilevel Neighborhoods of Atoms (MNA) structure descriptors for large database (>250000) of compounds with known functional activities and can estimate the probability of 4000 kinds of biological activities of compounds using structural formulas ^126^.

Average accuracy of prediction estimated in leave-one-out cross-validation procedure (each compound is excluded from the training set and its activity predicted based on SAR model obtained on the rest part of the training set) for the whole PASS training set is about 95% ^126^. This approach is widely used by pharmacologists and toxicologists to estimate potential drug effects ^127–129^. Here we evaluated the probability of studied 10 NBPEA compounds to have specific functional activity (Pa, %) *in vivo* also comparing these predictions to drugs studied in AI-driven phenotyping including ketamine, PCP, MDMA, LSD and THC. Probability “to be active” (Pa) is a qualitative parameter appraised the percent of probability belong to the class of chemical substance having similar biological activity i.e. activate similar receptors, transporters, and enzymes like substances with known activity^126^. Only activities that had summed probability higher than 60% for all compounds were used for further comparison, thus using only activities that had high prediction level or were represented in multiple compounds simultaneously, allowing better comparison. Due to the relatively high amount of determined activities for PCP comparing to other drugs, it was excluded from further analysis due to bad clustering both using principal components analysis (PCA) and hierarchical clustering (Supplementary Figures S29-S30). All activities that were not reported for a drug were considered as 0% probability for further analyses. Finally, using the remaining compound the hierarchical clustering calculation was conducted along with heatmap construction using pheatmap ^130^ package in R ^123^ and PCA plot was constructed using function prcomp and ggbiplot package for 2 top principal components explaining the most variance between the compounds (more than 50% together).

Finally, to support the *in-silico* activity clustering results we utilized molecular fingerprinting approach. Molecular fingerprints are widely and for a long time used in drug discovery and virtual compounds screening^131, 132^. Overall, they provide a virtual screening performance similar to other more complex methods, however are easier and faster to perform^131, 132^. Here we used Open Babel FP4^133^ fingerprinting method, based on 307 key functional substructure SMILES (Simplified Molecular Input Line Entry System) arbitrary target specification (SMARTS) patterns on the similar set of 10 NBPEA compounds and 5 reference compounds ketamine, PCP, MDMA, LSD and THC. We have further used NxN and Taylor-Butina clustering approaches on the evaluated FP4 fingerprints with thresholds = 0.5. All manipulations were performed using cheminformatics usegalaxy.org cloud (https://cheminformatics.usegalaxy.eu/)134.

## Acknowledgements

The laboratory is supported by SPSU state budgetary funds АААА-А19-119092790064-0 (Pure ID 73026081), and the budgetary state funding from the Ministry of Healthcare of Russian Federation (projects 121040200141-4 and 121031100292-2). AVK is the Chair of the International Zebrafish Neuroscience Research Consortium (ZNRC) and President of the International Stress and Behavior Society (ISBS, www.stress-and-behavior.com) that coordinated this collaborative multi-laboratory project. The consortium provided a collaborative idea exchange platform for this study. It is not considered as an affiliation, and did not fund the study. He is also supported by the Southwest University Zebrafish Platform Construction Fund. OVK is supported by the subsidy allocated to the Kazan Federal University for the state research assignment (project 0671-2020-0058); the research was performed using the equipment of Interdisciplinary Center for shared use of Kazan Federal University. The funders had no role in the design, analyses, and interpretation of the submitted study, or decision to publish.

## Additional information

The authors declare no conflicts of interest.

## Ethical Confirmation statements

Animal experiments were approved by IACUC of St. Petersburg State University and fully adhered to the National and Institutional guidelines and regulations on animal experimentation, as well as the 3Rs principles of humane animal experimentation.

## Data availability

The datasets generated and/or analyzed during the current study are available from the corresponding authors upon reasonable requests.

## CRediT authorship contribution statement

Konstantin A. Demin (Investigation) (Methodology) (Project administration) (Validation) (Writing - original draft) (Writing - review and editing), Olga V. Kupriyanova (Investigation) (Methodology) (Formal analysis) (Writing - Original Draft) (Writing - review and editing), Vadim A. Shevyrin (Investigation) (Methodology) (Formal analysis) (Writing - Original Draft) (Writing - review and editing), Ksenia A. Derzhavina (Investigation) (Formal analysis) (Writing - original draft) (Writing - review and editing), Nataliya A. Krotova (Investigation) (Formal analysis) (Writing - review and editing), Nikita P. Ilyin (Investigation) (Formal analysis) (Writing - review and editing), Tatiana O. Kolesnikova (Investigation) (Writing - Original Draft) (Writing - review and editing), David S. Galstyan (Investigation) (Writing - Original Draft) (Writing - review and editing), Iurii M. Kositsyn (Investigation) (Writing - Original Draft) (Writing - review and editing), Abubakar-Askhab S. Khaybaev (Investigation) (Writing - review and editing), Maria V. Seredinskaya (Investigation) (Writing - review and editing), Yaroslav Dubrovskii (Investigation) (Methodology) (Writing - review and editing), Raziya G. Sadykova (Investigation) (Writing - review and editing), Maria O. Nerush (Investigation) (Writing - review and editing), Mikael S. Mor (Investigation) (Writing - review and editing), Elena V. Petersen (Methodology) (Writing - review and editing), Tatyana Strekalova (Methodology) (Writing - review and editing), Evgeniya V. Efimova (Investigation) (Methodology) (Writing - review and editing), Dmitrii V. Bozhko (Investigation) (Methodology) (Writing - review and editing), Vladislav O. Myrov (Investigation) (Methodology) (Writing - review and editing), Sofia M. Kolchanova (Investigation) (Methodology) (Writing - review and editing), Aleksander I. Polovian (Investigation) (Methodology) (Writing - review and editing), Georgii K. Galumov (Investigation) (Methodology) (Writing - review and editing), and Allan V. Kalueff (Conceptualization) (Funding acquisition) (Methodology) (Project administration) (Resources) (Supervision) (Validation) (Writing - review and editing)

