## Supplementary Materials for "Behavioral and neurochemical effects of novel N-Benzyl-2-phenylethylamine derivatives in adult zebrafish"

**Data availability**

The datasets generated and/or analyzed in the present study are available from the original article, supplementary materials, or the corresponding author (upon reasonable requests, for use in collaborative research projects and/or for joint publications resulting from such projects).

**Supplementary Table S1.** Dose-dependent behavioral alterations of the adult zebrafish in the Novel Tank Test induced by *N*-benzylphenethylamine derivatives assessed using Kruskal-Wallis test. Data is represented as mean ± S.E.M. (n=15-17). *p<0.05, **p<0.01, ***p<0.001 vs. control, post-hoc Dunn's test for significant Kruskal-Wallis data. Also see Figure 2 and Figure 3 in the main text for graphical representations.

| Endpoint | Compound | KW H statistics | KW P | Control | 5 mg/L | 10 mg/L | 20 mg/L |
|---|---|---|---|---|---|---|---|
| **Velocity, cm/s** | **24H-NBOMe** | **9.86** | **0.0198** | 3.388±0.3314 | 3.831±0.1505 | 2.843±0.1931 | 3.321±0.2143 |
| | 24H-NBF | 1.903 | 0.5928 | 3.528±0.3101 | 3.168±0.3213 | 3.543±0.2374 | 3.869±0.348 |
| | **24H-NBCl** | **8.84** | **0.0315** | 4.911±0.3324 | 4.22±0.283 | 4.016±0.1763 | 4.976±0.3188 |
| | **24H-NBBr** | **10.04** | **0.0183** | 3.405±0.4154 | 3.828±0.3292 | **4.936±0.2703**** | 4.196±0.3921 |
| | **24H-NBOMe(F)** | **15.78** | **0.0013** | 3.133±0.2935 | **4.265±0.2257*** | **5.107±0.342***** | 4.147±0.2742 |
| | 34H-NBOMe | 1.641 | 0.6501 | 3.217±0.2959 | 3.579±0.1629 | 3.194±0.2925 | 3.35±0.2079 |
| | 34H-NBF | 2.01 | 0.5703 | 2.792±0.2756 | 3.29±0.2988 | 2.556±0.274 | 2.85±0.361 |
| | 34H-NBCl | 4.203 | 0.2404 | 4.06±0.2668 | 3.38±0.3551 | 3.185±0.3877 | 3.249±0.3157 |
| | 34H-NBBr | 2.105 | 0.5508 | 3.254±0.3627 | 3.17±0.3494 | 2.872±0.3487 | 2.661±0.2953 |
| | 34H-NBOMe(F) | 3.893 | 0.2732 | 3.758±0.2804 | 3.547±0.3239 | 4.179±0.2355 | 4.093±0.1968 |
| **Time spent top, s** | **24H-NBOMe** | **13.22** | **0.0042** | 112.3±21.93 | 83.7±12.96 | **40.13±12.59*** | **45.51±18.42*** |
| | 24H-NBF | 6.071 | 0.1082 | 149.2±18.77 | 168.3±20.17 | 140.7±24.37 | 95.19±17.95 |
| | **24H-NBCl** | **15.26** | **0.0016** | 133.2±14.33 | 102±15.61 | **56.37±12.01**** | **64.52±19.32**** |
| | 24H-NBBr | 2.372 | 0.499 | 82.37±14.12 | 90.63±19.13 | 63.12±20.87 | 68.73±20.33 |
| | **24H-NBOMe(F)** | **10.32** | **0.0161** | 58.9±15.79 | **177.4±26.98*** | 120.8±31.93 | 43.9±12.46 |
| | 34H-NBOMe | 1.229 | 0.746 | 92.47±20.75 | 120.2±20.13 | 89.74±18.75 | 97.08±20.24 |
| | 34H-NBF | 7.329 | 0.0621 | 95.41±19.09 | 153.1±20.28 | 166.3±20.49 | 157.7±20.6 |
| | 34H-NBCl | 3.562 | 0.3128 | 147.7±18.73 | 161.7±24.33 | 121.6±24.73 | 179.4±23.89 |
| | 34H-NBBr | 3.668 | 0.2996 | 115.5±19.95 | 133.3±14.87 | 158.4±21.96 | 172.9±25.35 |
| | **34H-NBOMe(F)** | **37.48** | **<0.0001** | 115.2±17.52 | 110.2±24.12 | **249.8±13.42***** | **281.7±8.045***** |
| **Number of top entries** | **24H-NBOMe** | **22.99** | **<0.0001** | 8.188±1.539 | 8.188±0.8525 | **2.188±0.7143**** | **2.938±0.7717*** |
| | **24H-NBF** | **12.34** | **0.0063** | 9.25±1.216 | 7.125±1.303 | 6.938±1.574 | **3.563±0.9308**** |
| | **24H-NBCl** | **32.14** | **<0.0001** | 16.94±1.697 | 10.53±1.493 | **5.118±1.221***** | **2.412±0.6363***** |
| | **24H-NBBr** | **20.03** | **0.0002** | 8.438±1.541 | 3.438±0.5475 | **2.313±0.5379**** | **1.375±0.482***** |
| | **24H-NBOMe(F)** | **9.994** | **0.0186** | 5.625±1.399 | **3.438±0.689*** | 1.25±0.2814 | 2.313±0.5222 |
| | 34H-NBOMe | 4.466 | 0.2154 | 4.625±0.9656 | 7.118±1.144 | 4.188±1.089 | 4.75±0.8539 |
| | 34H-NBF | 2.427 | 0.4886 | 6.25±1.286 | 9.5±1.751 | 7.188±1.145 | 6.563±1.777 |
| | 34H-NBCl | 7.708 | 0.0524 | 9.813±1.089 | 7.313±1.393 | 7.059±1.984 | 5.125±0.9953 |
| | **34H-NBBr** | **9.472** | **0.0236** | 7.875±1.36 | 7.5±1.012 | 5±1.208 | **3.313±0.7229*** |
| | **34H-NBOMe(F)** | **20.22** | **0.0002** | 10.63±1.455 | 5.563±1.372 | **3.188±0.9139**** | **2.5±0.9037***** |
| **Latency to enter top, s** | **24H-NBOMe** | **17.57** | **0.0005** | 9.819±4.938 | 60.71±18.87 | **163.2±31.6***** | **115.6±31.94*** |
| | 24H-NBF | 4.999 | 0.1719 | 32.43±19.72 | 9.171±7.56 | 38.58±17.47 | 96.87±30.94 |
| | 24H-NBCl | 3.725 | 0.2927 | 36.8±19.64 | 33.54±19.02 | 39.04±21.83 | 92.77±29.65 |
| | 24H-NBBr | 7.147 | 0.0674 | 69.34±25.47 | 42.68±20.97 | 105.9±29.74 | 136.3±31.06 |
| | **24H-NBOMe(F)** | **12.09** | **0.0071** | 100.4±24.74 | **21.61±12.22*** | 62.91±30.14 | 113.7±30.08 |
| | 34H-NBOMe | 3.697 | 0.2961 | 129±28.92 | 82.8±22.71 | 140.9±25.37 | 100.4±19.18 |
| | **34H-NBF** | **9.096** | **0.028** | 62.26±23.32 | 39.31±14.39 | **6.129±3.816*** | 31.45±11.95 |
| | 34H-NBCl | 6.851 | 0.0768 | 32.26±11.99 | 57.23±25.81 | 91.86±31.04 | 13.96±9.678 |
| | 34H-NBBr | 5.039 | 0.169 | 75.17±25.83 | 38.72±12.73 | 27.27±10.98 | 17.42±13.06 |
| | **34H-NBOMe(F)** | **16.95** | **0.0007** | 55.19±19.02 | 47.97±25.54 | **0±0***** | **0±0***** |
| **Horizontal (left-right) alterations** | 24H-NBOMe | 1.511 | 0.6797 | 6.688±1.368 | 8.563±1.923 | 5.5±1.092 | 5.875±0.9169 |
| | 24H-NBF | 2.442 | 0.4858 | 9.563±1.281 | 9.5±1.751 | 10.25±1.953 | 16±3.649 |
| | **24H-NBCl** | **14.91** | **0.0019** | 10±1.687 | 9.529±1.768 | 8.941±1.934 | **24.59±4.021*** |

| | | | | | | | |
|---|---|---|---|---|---|---|---|
| | **24H-NBBr** | **31.61** | **<0.0001** | 4.5±1.594 | 9.063±1.916 | **23.81±3.238*** ** | **28.81±4.643*** ** |
| | **24H-NBOMe(F)** | **30.6** | **<0.0001** | 2.5±0.3873 | **10.56±2.16*** | **19.94±3.944*** ** | **27.38±3.979*** ** |
| | 34H-NBOMe | 0.5185 | 0.9148 | 4.938±1.142 | 6.529±1.591 | 5.25±1.349 | 4.25±0.9106 |
| | 34H-NBF | 1.211 | 0.7503 | 3.75±0.7159 | 5.125±1.372 | 5.375±1.224 | 6.438±1.666 |
| | 34H-NBCl | 5.847 | 0.1193 | 9.125±2.196 | 2.75±0.4518 | 7.294±1.921 | 7.375±1.589 |
| | 34H-NBBr | 3.528 | 0.3172 | 7.125±1.408 | 10.44±1.313 | 9.063±1.995 | 8.125±1.821 |
| | **34H-NBOMe(F)** | **11.87** | **0.0078** | 6.563±0.9441 | 10.31±2.25 | 10.69±1.675 | **17.44±2.827** ** |
| **The horizontal shuttling behavior** | 24H-NBOMe | 0.09591 | 0.9923 | 1.834±0.268 | 2.206±0.4661 | 2.096±0.4848 | 1.819±0.2522 |
| | 24H-NBF | 1.698 | 0.6373 | 2.962±0.3668 | 3.201±0.5557 | 2.958±0.581 | 4.061±0.6698 |
| | **24H-NBCl** | **12.8** | **0.0051** | 2.065±0.3385 | 2.247±0.4067 | 2.239±0.3924 | **5.032±0.8101** ** |
| | **24H-NBBr** | **33.7** | **<0.0001** | 1.169±0.3197 | 2.259±0.414 | **4.739±0.5747*** ** | **6.5±0.7136*** ** |
| | **24H-NBOMe(F)** | **26.65** | **<0.0001** | 1.072±0.2572 | 2.447±0.4761 | **3.869±0.6846** ** | **6.461±0.7111*** ** |
| | 34H-NBOMe | 0.9949 | 0.8025 | 1.508±0.2595 | 1.794±0.4402 | 1.592±0.2973 | 1.251±0.241 |
| | 34H-NBF | 2.156 | 0.5407 | 1.411±0.2401 | 1.557±0.3315 | 1.949±0.3783 | 2.014±0.3844 |
| | 34H-NBCl | 7.098 | 0.0688 | 2.741±0.8056 | 0.7975±0.1354 | 2.245±0.5064 | 2.525±0.6578 |
| | 34H-NBBr | 5.984 | 0.1124 | 2.029±0.2683 | 3.481±0.4732 | 3.096±0.5947 | 3.286±0.5298 |
| | **34H-NBOMe(F)** | **9.35** | **0.025** | 1.709±0.2008 | 2.911±0.5969 | 2.703±0.4807 | **4.283±0.66** ** |

**Supplementary Table S2.** Spearman R correlations of Velocity and Horizontal (left-right) alterations or Velocity and Horizontal (left-right) alterations relative to velocity in Novel Tank Test studies among all control groups (n=173, including GZLM). While Horizontal (left-right) alterations correlates with velocity, thus disturbing the endpoint with locomotion data, relative endpoint has no correlation, thus may be used effectively to assess trans/hallucination-like circling behavior in adult zebrafish.

| | **Velocity vs. naïve alterations** | **Velocity vs. relative alterations** |
|---|---|---|
| **r** | 0.4247 | -0.00951 |
| **95% confidence interval** | 0.2901 to 0.5429 | -0.1628 to 0.1442 |
| **P (two-tailed)** | <0.0001 | 0.9012 |

**Supplementary Table S3.** Dose-dependent neurochemical alterations in the adult zebrafish induced by selected *N*-benzylphenethylamine derivatives assessed using Kruskal-Wallis test. Data is represented as mean ± S.E.M. (n=8-10). *p<0.05, **p<0.01, ***p<0.001 vs. control, post-hoc Dunn's test for significant Kruskal-Wallis data. Also see Figure 4 in the main text for graphical representations.

| **Endpoint** | **KWH statistics** | **KWP** | **Control** | **24H-NBF** | **24H-NBBr** | **24H-NBOMe(F)** | **34H-NBOMe** | **34H-NBF** | **34H-NBCl** | **34H-NBBr** | **34H-NBOMe(F)** |
|---|---|---|---|---|---|---|---|---|---|---|---|
| **Norepinephrine, pg/mg** | 10.32 | 0.2432 | 1172±113.9 | 877.9±82.16 | 879.2±107.6 | 836±111.5 | 1219±139.2 | 893.5±88.9 | 857.4±131 | 1134±119.5 | 941.7±198.3 |
| **Serotonin, pg/mg** | 32.29 | <0.0001 | 254.2±28.42 | **140.3±24.47*** | **139.8±11.99*** | **118.7±19.66** ** | 221.4±8.774 | 163.1±21.96 | 139.8±23.8 | 225±18.04 | **114±21.94** ** |
| **5HIAA, pg/mg** | 21.74 | 0.0054 | 82.7±4.863 | 104.6±8.995 | 107.8±5.117 | 117.4±11.94 | 85.58±6.354 | 69.2±7.233 | 89.23±7.672 | 109.5±5.867 | 101.9±15.54 |
| **Dopamine, pg/mg** | 26.65 | 0.0008 | 253.6±25.61 | 155.2±14.78 | **148.5±18.02*** | **138.6±16.9*** | 220±7.773 | 169.3±20.33 | **143.5±21.24*** | 215±21.22 | **139.9±30.27*** |
| **DOPAC, pg/mg** | 14.02 | 0.0813 | 37.52±5.99 | 52.64±7.618 | 56.79±10.27 | 64.33±6.131 | 43.39±4.59 | 38.12±5.146 | 51.65±5.804 | 60.88±8.911 | 62.94±10.61 |
| **HVA, pg/mg** | 11.1 | 0.196 | 12.33±1.831 | 19.36±3.252 | 19.85±4.748 | 27.28±2.995 | 13.59±2.044 | 29.7±11.28 | 20.21±4.161 | 21.05±6.207 | 23.53±5.899 |
| **5HIAA to serotonin ratio** | 47.36 | <0.0001 | 0.3663±0.04798 | **0.8153±0.0756** ** | **0.804±0.06216** ** | **1.134±0.1808*** ** | 0.3859±0.02348 | 0.455±0.05314 | **1.04±0.3493*** | 0.5044±0.0418 | **1.086±0.2175*** ** |

| | | | | | | | | | | |
|---|---|---|---|---|---|---|---|---|---|---|
| **DOPAC to dopamine ratio** | 35.1 | <0.0001 | 0.1708±0.04541 | 0.3594±0.06167 | **0.3723±0.04004*** | **0.4854±0.04476***** | 0.1966±0.0198 | 0.2487±0.0511 | **0.4932±0.1199**** | 0.2953±0.04914 | **0.5087±0.1035**** |
| **HVA to dopamine ratio** | 27.26 | 0.0006 | 0.05678±0.01381 | 0.1326±0.02366 | 0.1394±0.03425 | **0.2128±0.03583***** | 0.06245±0.01046 | 0.175±0.0567 | **0.1605±0.0381*** | 0.1037±0.03459 | **0.1815±0.04801*** |

**Supplementary Table S4.** Results of Generalized Linear Model (GZLM) fits using substitutions in the phenethylamine moiety, *N*-benzyl fragment of molecule and their interaction effects as ‘predictors’, to compare effects of *N*-benzylphenethylamine derivatives. The corrected Akaike information criterion (AICc) was used to choose the ‘best fit’ model among Gaussian distribution (identity link), Poisson distribution (with log link), Gamma distribution (inverse and log links) and Inverse Gaussian distribution (with inverse and log links) for the novel tank test (NTT). Also see Figures 5 and 6 in the main text for graphical representations and Supplementary Tables S5-8 for ANOVA and Tukey tests results.

| Predictor | *b* | 95% CI | t | p.value |
|---|---|---|---|---|
| | | **Velocity, cm/s** | | |
| Intercept | 3.83 | [3.01, 4.65] | 9.14 | < .001 |
| Phenethylamine: 24H | 1.20 | [0.01, 2.38] | 1.98 | .051 |
| Benzyl: NBBr | 0.07 | [-1.09, 1.23] | 0.11 | .909 |
| Benzyl: NBCl | 0.45 | [-0.71, 1.61] | 0.77 | .445 |
| Benzyl: NBF | -0.75 | [-1.91, 0.41] | -1.26 | .211 |
| Benzyl: NBOMe | 0.49 | [-0.67, 1.65] | 0.83 | .410 |
| Phenethylamine: 24H × Benzyl: NBBr | -0.03 | [-1.69, 1.63] | -0.04 | .970 |
| Phenethylamine: 24H × Benzyl: NBCl | -1.30 | [-2.96, 0.36] | -1.53 | .129 |
| Phenethylamine: 24H × Benzyl: NBF | -0.74 | [-2.40, 0.92] | -0.87 | .385 |
| **Phenethylamine: 24H × Benzyl: NBOMe** | **-1.70** | **[-3.36, -0.04]** | **-2.01** | **.047** |
| | | **Time spent top, s** | | |
| Intercept | 170.16 | [129.57, 210.75] | 8.22 | < .001 |
| **Phenethylamine: 24H** | **-117.10** | **[-175.80, -58.40]** | **-3.91** | **< .001** |
| **Benzyl: NBBr** | **-72.14** | **[-129.54, -14.73]** | **-2.46** | **.015** |
| **Benzyl: NBCl** | **-82.78** | **[-140.19, -25.37]** | **-2.83** | **.006** |
| **Benzyl: NBF** | **-115.89** | **[-173.30, -58.48]** | **-3.96** | **< .001** |
| Benzyl: NBOMe | -57.12 | [-114.53, 0.29] | -1.95 | .054 |
| **Phenethylamine: 24H × Benzyl: NBBr** | **84.88** | **[2.77, 166.98]** | **2.03** | **.045** |
| **Phenethylamine: 24H × Benzyl: NBCl** | **85.49** | **[3.39, 167.60]** | **2.04** | **.044** |
| **Phenethylamine: 24H × Benzyl: NBF** | **117.64** | **[35.53, 199.74]** | **2.81** | **.006** |
| Phenethylamine: 24H × Benzyl: NBOMe | 53.38 | [-28.73, 135.48] | 1.27 | .205 |
| | | **Horizontal (left-right) alterations** | | |
| Intercept | 16.92 | [8.86, 24.97] | 4.12 | < .001 |
| **Phenethylamine: 24H** | **14.54** | **[2.89, 26.19]** | **2.45** | **.016** |
| Benzy: lNBBr | -6.83 | [-18.23, 4.56] | -1.18 | .242 |
| Benzyl: NBCl | -8.33 | [-19.73, 3.06] | -1.43 | .155 |
| Benzyl: NBF | -10.67 | [-22.06, 0.73] | -1.83 | .069 |
| Benzyl: NBOMe | -8.50 | [-19.89, 2.89] | -1.46 | .147 |
| Phenethylamine: 24H × Benzyl: NBBr | 5.30 | [-11.00, 21.59] | 0.64 | .525 |
| Phenethylamine: 24H × Benzyl: NBCl | 0.21 | [-16.08, 16.51] | 0.03 | .980 |
| Phenethylamine: 24H × Benzyl: NBF | -13.70 | [-30.00, 2.59] | -1.65 | .102 |
| Phenethylamine: 24H × Benzyl: NBOMe | -15.54 | [-31.83, 0.76] | -1.87 | .064 |
| | | **The horizontal shuttling behavior** | | |

| | | | | |
|---|---|---|---|---|
| Intercept | 4.42 | [3.12, 5.73] | 6.63 | < .001 |
| **Phenethylamine: 24H** | **1.96** | **[0.07, 3.85]** | **2.04** | **.044** |
| Benzyl: NBBr | -1.72 | [-3.57, 0.13] | -1.82 | .071 |
| **Benzyl: NBCl** | **-2.40** | **[-4.25, -0.55]** | **-2.54** | **.012** |
| **Benzyl: NBF** | **-2.39** | **[-4.24, -0.54]** | **-2.54** | **.013** |
| **Benzyl: NBOMe** | **-2.36** | **[-4.21, -0.51]** | **-2.50** | **.014** |
| Phenethylamine: 24H × Benzyl: NBBr | 0.42 | [-2.22, 3.06] | 0.31 | .756 |
| Phenethylamine: 24H × Benzyl: NBCl | 0.47 | [-2.18, 3.11] | 0.35 | .729 |
| Phenethylamine: 24H × Benzyl: NBF | -1.92 | [-4.56, 0.72] | -1.42 | .158 |
| Phenethylamine: 24H × Benzyl: NBOMe | -2.10 | [-4.74, 0.55] | -1.55 | .123 |
| **Number of top entries** | | | | |
| Intercept | 2.00 | [-1.05, 5.05] | 1.29 | .201 |
| Phenethylamine: 24H | 1.27 | [-3.14, 5.68] | 0.57 | .573 |
| **Benzyl: NBBr** | **6.92** | **[2.60, 11.23]** | **3.14** | **.002** |
| **Benzyl: NBCl** | **9.42** | **[5.10, 13.73]** | **4.28** | **< .001** |
| Benzy: lNBF | 3.58 | [-0.73, 7.90] | 1.63 | .106 |
| **Benzy: lNBOMe** | **7.50** | **[3.19, 11.81]** | **3.41** | **.001** |
| **Phenethylamine: 24H × Benzyl: NBBr** | **-6.61** | **[-12.77, -0.44]** | **-2.10** | **.038** |
| **Phenethylamine: 24H × Benzyl: NBCl** | **-10.36** | **[-16.52, -4.19]** | **-3.29** | **.001** |
| Phenethylamine: 24H × Benzyl: NBF | -1.27 | [-7.44, 4.89] | -0.40 | .687 |
| Phenethylamine: 24H × Benzyl: NBOMe | -6.19 | [-12.36, -0.02] | -1.97 | .052 |
| **Latency to enter top, s** | | | | |
| Intercept | 64.74 | [5.97, 123.50] | 2.16 | .033 |
| Phenethylamine: 24H | 73.62 | [-11.36, 158.59] | 1.70 | .092 |
| Benzyl: NBBr | 7.20 | [-75.91, 90.31] | 0.17 | .865 |
| Benzyl: NBCl | 15.19 | [-67.91, 98.30] | 0.36 | .721 |
| **Benzyl: NBF** | **90.84** | **[7.73, 173.95]** | **2.14** | **.034** |
| Benzyl: NBOMe | -4.37 | [-87.48, 78.74] | -0.10 | .918 |
| Phenethylamine: 24H × Benzyl: NBBr | -82.52 | [-201.38, 36.35] | -1.36 | .176 |
| Phenethylamine: 24H × Benzyl: NBCl | 14.01 | [-104.86, 132.87] | 0.23 | .818 |
| Phenethylamine: 24H × Benzyl: NBF | -89.07 | [-207.93, 29.79] | -1.47 | .145 |
| Phenethylamine: 24H × Benzyl: NBOMe | 43.41 | [-75.46, 162.27] | 0.72 | .476 |

**Supplementary Table S5.** ANOVA data pair-wise comparison using the Generalized Linear Model (GZLM) fits using substitutions in the phenethylamine moiety, *N*-benzyl fragment of molecule and their interaction effects as ‘predictors’, to compare effects of *N*-benzylphenethylamine derivatives in the Novel Tank Test. Also see Figures 4 and 5 in the main text for graphical representations and Supplementary Tables S4, and 6-8 for GZLM and Tukey tests results.

| **Factor** | **Df** | **Chisq** | **P** |
|---|---|---|---|
| | | **Velocity, cm/s** | |
| Phenethylamine | 1 | 2.67959841435305 | 0.10164135698357 |
| **Benzyl** | **4** | **9.98055346123705** | **0.040756533502772** |
| Phenethylamine:Benzyl | 4 | 6.44476938048346 | 0.168303794948819 |
| | | **Time spent top, s** | |
| **Phenethylamine** | **1** | **13.4362567832127** | **0.00024680758555913** |
| Benzyl | 4 | 8.27228726078573 | 0.0820980430505072 |
| Phenethylamine:Benzyl | 4 | 8.96052146294373 | 0.0620938671289181 |

| | | | |
|---|---|---|---|
| | **Horizontal (left-right) alterations** | | |
| **Phenethylamine** | **1** | **13.9413684958026** | **0.000188601648671479** |
| **Benzyl** | **4** | **26.5658111764639** | **2.43289835555797e-05** |
| **Phenethylamine:Benzyl** | **4** | **10.1701204391803** | **0.03765748285029** |
| | **The horizontal shuttling behavior** | | |
| **Phenethylamine** | **1** | **9.88588703789612** | **0.00166551223244028** |
| **Benzyl** | **4** | **35.0894245929784** | **4.45295558219627e-07** |
| Phenethylamine:Benzyl | 4 | 7.31020634251079 | 0.120375519013203 |
| | **Number of top entries** | | |
| **Phenethylamine** | **1** | **13.6804709268008** | **0.00021669625812774** |
| **Benzyl** | **4** | **10.8484322951132** | **0.0283212881803089** |
| **Phenethylamine:Benzyl** | **4** | **14.5581456681246** | **0.00571111017083243** |
| | **Latency to enter top, s** | | |
| **Phenethylamine** | **1** | **7.05257617626734** | **0.00791513356296606** |
| Benzyl | 4 | 7.93447377542106 | 0.0940082099707969 |
| Phenethylamine:Benzyl | 4 | 7.89311170127418 | 0.0955730594095111 |

**Supplementary Table S6.** Post-hoc Tukey's test results for significant phenethylamine predictor ANOVA data pair-wise comparison using the Generalized Linear Model (GZLM) fits using substitutions in the phenethylamine moiety, *N*-benzyl fragment of molecule and their interaction effects as 'predictors', to compare effects of *N*-benzylphenethylamine derivatives in the Novel Tank Test. Also see Figures 5 and 6 in the main text for graphical representations and Supplementary Tables S4, 5, 7, and 8 for GZLM, ANOVA and other Tukey tests results.

| **Contrast** | **estimated mean** | **SE** | **z.ratio** | **p.value** |
|---|---|---|---|---|
| | | **Time spent top, s** | | |
| 34H - 24H | 48.8252504757576 | 13.1587213673859 | 3.71048592888149 | 0.000206861795057501 |
| | | **Horizontal (left-right) alterations** | | |
| 34H - 24H | -9.79090909090912 | 2.61148780915206 | -3.74916898198663 | 0.000177421511132918 |
| | | **The horizontal shuttling behavior** | | |
| 34H - 24H | -1.33794191473031 | 0.423850864981386 | -3.15663367771837 | 0.00159601712408099 |
| | | **Number of top entries** | | |
| 34H - 24H | 3.61212121212121 | 0.988330130337898 | 3.6547719241204 | 0.00025741070893214 |
| | | **Latency to enter top, s** | | |
| 34H - 24H | -50.7813249545454 | 19.0494468151706 | -2.66576375929741 | 0.0076813649664175 |

**Supplementary Table S7.** Post-hoc Tukey's test results for significant benzyl predictor ANOVA data pair-wise comparison using the Generalized Linear Model (GZLM) fits using substitutions in the phenethylamine moiety, *N*-benzyl fragment of molecule and their interaction effects as 'predictors', to compare effects of *N*-benzylphenethylamine derivatives in the Novel Tank Test. Also see Figures 5 and 6 in the main text for graphical representations and Supplementary Tables S4-6, and 8 for GZLM, ANOVA and other Tukey tests results.

| **Contrast** | **estimated mean** | **SE** | **z.ratio** | **p.value** |
|---|---|---|---|---|
| | | **Velocity, cm/s** | | |
| NBOMe(F) - NBBr | -0.051705166666667 | 0.423455574648762 | -0.12210293065466 | 0.902817500921662 |
| NBOMe(F) - NBCl | 0.1940675 | 0.423455574648762 | 0.458294828592044 | 0.646740638974382 |
| **NBOMe(F) - NBF** | **1.114628875** | **0.423455574648762** | **2.63222151680146** | **0.00848285343492947** |
| NBOMe(F) - NBOMe | 0.361503708333332 | 0.423455574648762 | 0.853699254362593 | 0.393271647400656 |

| | | | | |
|---|---|---|---|---|
| NBBr - NBCl | 0.245772666666667 | 0.41872407894165 | 0.586956134187152 | 0.557233167861004 |
| **NBBr - NBF** | **1.16633404166667** | **0.41872407894165** | **2.78544774548109** | **0.00534538548423966** |
| NBBr - NBOMe | 0.413208874999999 | 0.41872407894165 | 0.986828548394945 | 0.323726696829956 |
| **NBCl - NBF** | **0.920561374999998** | **0.41872407894165** | **2.19849161129394** | **0.0279140917020914** |
| NBCl - NBOMe | 0.167436208333332 | 0.41872407894165 | 0.399872414207792 | 0.689250491252232 |
| NBF - NBOMe | -0.753125166666667 | 0.41872407894165 | -1.79861919708615 | 0.0720789385481587 |
| | | **Horizontal (left-right) alterations** | | |
| NBOMe(F) - NBBr | 4.18560606060607 | 4.15693064758429 | 1.00689821780848 | 0.313983688525581 |
| **NBOMe(F) - NBCl** | **8.22727272727274** | **4.15693064758429** | **1.97917007156562** | **0.0477968613101093** |
| **NBOMe(F) - NBF** | **17.5189393939394** | **4.15693064758429** | **4.21439299308978** | **2.50450860074529e-05** |
| **NBOMe(F) - NBOMe** | **16.2689393939394** | **4.15693064758429** | **3.91369035790716** | **9.08961814500771e-05** |
| NBBr - NBCl | 4.04166666666667 | 4.11048303727683 | 0.983258325119923 | 0.325480315529613 |
| **NBBr - NBF** | **13.3333333333333** | **4.11048303727683** | **3.24373880451933** | **0.00117971872940261** |
| **NBBr - NBOMe** | **12.0833333333333** | **4.11048303727683** | **2.93963829159565** | **0.00328595597036256** |
| **NBCl - NBF** | **9.29166666666664** | **4.11048303727683** | **2.2604804793994** | **0.0237914466148196** |
| NBCl - NBOMe | 8.04166666666667 | 4.11048303727683 | 1.95637996647572 | 0.0504204103406674 |
| NBF - NBOMe | -1.24999999999997 | 4.11048303727683 | -0.30410051292368 | 0.761051316078089 |
| | | **The horizontal shuttling behavior** | | |
| **NBOMe(F) - NBBr** | **1.51043632807576** | **0.674680021278111** | **2.23874470925402** | **0.0251725316747629** |
| **NBOMe(F) - NBCl** | **2.16213328061743** | **0.674680021278111** | **3.20467957020795** | **0.00135212917384209** |
| **NBOMe(F) - NBF** | **3.35200751461742** | **0.67468002127811** | **4.96829224062007** | **6.75451155593782e-07** |
| **NBOMe(F) - NBOMe** | **3.41163723715909** | **0.674680021278111** | **5.05667446724761** | **4.26630569018105e-07** |
| NBBr - NBCl | 0.651696952541667 | 0.667141460410186 | 0.976849725605386 | 0.328643554192409 |
| **NBBr - NBF** | **1.84157118654166** | **0.667141460410186** | **2.76039085535081** | **0.00577322451322665** |
| **NBBr - NBOMe** | **1.90120090908334** | **0.667141460410186** | **2.84977178290553** | **0.00437506089588123** |
| NBCl - NBF | 1.189874234 | 0.667141460410186 | 1.78354112974543 | 0.0744982589637971 |
| NBCl - NBOMe | 1.24950395654167 | 0.667141460410186 | 1.87292205730014 | 0.0610791488185606 |
| NBF - NBOMe | 0.0596297225416722 | 0.667141460410186 | 0.0893809275547189 | 0.928779180404247 |
| | | **Number of top entries** | | |
| **NBOMe(F) - NBBr** | **-3.61363636363637** | **1.57321041068408** | **-2.29698223396897** | **0.0216197832373683** |
| **NBOMe(F) - NBCl** | **-4.23863636363637** | **1.57321041068408** | **-2.69425903544159** | **0.0070545315286075** |
| NBOMe(F) - NBF | -2.94696969696971 | 1.57321041068408 | -1.87322031239818 | 0.0610379685188806 |
| **NBOMe(F) - NBOMe** | **-4.40530303030304** | **1.57321041068408** | **-2.80019951583429** | **0.00510710302710329** |
| NBBr - NBCl | -0.625000000000001 | 1.55563208901313 | -0.401765947369017 | 0.687856285366683 |
| NBBr - NBF | 0.666666666666662 | 1.55563208901313 | 0.428550343860281 | 0.668250488865895 |
| NBBr - NBOMe | -0.791666666666671 | 1.55563208901313 | -0.50890353333409 | 0.610819843196171 |
| NBCl - NBF | 1.29166666666666 | 1.55563208901313 | 0.830316291229298 | 0.406359979559635 |
| NBCl - NBOMe | -0.16666666666667 | 1.55563208901313 | -0.107137585965073 | 0.914679829469469 |
| NBF - NBOMe | -1.45833333333333 | 1.55563208901313 | -0.93745387719437 | 0.348525138213503 |

**Supplementary Table S8.** Post-hoc Tukey's test results for significant phenethylamine and benzyl interaction predictor ANOVA data pair-wise comparison using the Generalized Linear Model (GZLM) fits using substitutions in the phenethylamine moiety, *N*-benzyl fragment of molecule and their interaction effects as 'predictors', to compare effects of *N*-benzylphenethylamine derivatives in the Novel Tank Test. Also see Figures 5 and 6 in the main text for graphical representations and Supplementary Tables S4-6, and 8 for GZLM, ANOVA and other Tukey tests results.

| Contrast | estimated mean | SE | z.ratio | p.value |
|---|---|---|---|---|

| | | | | |
|---|---|---|---|---|
| | | **Horizontal (left-right) alterations** | | |
| **34H-NBOMe(F) - 24H-NBOMe(F)** | **-14.5378787878788** | **5.94374865180395** | **-2.44591076095833** | **0.0144486758125472** |
| 34H-NBOMe(F) - 34H-NBBr | 6.83333333333334 | 5.81310085922144 | 1.1755057238502 | 0.23979244856145 |
| **34H-NBOMe(F) - 24H-NBBr** | **-13** | **5.81310085922145** | **-2.23632796244673** | **0.0253302960802228** |
| 34H-NBOMe(F) - 34H-NBCl | 8.33333333333334 | 5.81310085922145 | 1.43354356567098 | 0.151702558060722 |
| 34H-NBOMe(F) - 24H-NBCl | -6.41666666666669 | 5.81310085922145 | -1.10382854556666 | 0.269667519603883 |
| 34H-NBOMe(F) - 34H-NBF | 10.6666666666666 | 5.81310085922145 | 1.83493576405885 | 0.0665152077731022 |
| 34H-NBOMe(F) - 24H-NBF | 9.83333333333329 | 5.81310085922145 | 1.69158140749174 | 0.0907258139481679 |
| 34H-NBOMe(F) - 34H-NBOMe | 8.50000000000002 | 5.81310085922144 | 1.4622144369844 | 0.143682456661366 |
| 34H-NBOMe(F) - 24H-NBOMe | 9.49999999999998 | 5.81310085922144 | 1.63423966486491 | 0.102208536947565 |
| **24H-NBOMe(F) - 34H-NBBr** | **21.3712121212122** | **5.94374865180394** | **3.59557803890748** | **0.000323672078895933** |
| 24H-NBOMe(F) - 24H-NBBr | 1.5378787878788 | 5.94374865180394 | 0.258738866323368 | 0.79583672941424 |
| **24H-NBOMe(F) - 34H-NBCl** | **22.8712121212122** | **5.94374865180394** | **3.84794402674997** | **0.000119113258213124** |
| 24H-NBOMe(F) - 24H-NBCl | 8.12121212121214 | 5.94374865180394 | 1.36634514629877 | 0.171830657094428 |
| **24H-NBOMe(F) - 34H-NBF** | **25.2045454545455** | **5.94374865180394** | **4.24051334117163** | **2.23009178952326e-05** |
| **24H-NBOMe(F) - 24H-NBF** | **24.3712121212121** | **5.94374865180393** | **4.10031001459246** | **4.12597041518334e-05** |
| **24H-NBOMe(F) - 34H-NBOMe** | **23.0378787878788** | **5.94374865180394** | **3.87598469206581** | **0.000106194360618718** |
| **24H-NBOMe(F) - 24H-NBOMe** | **24.0378787878788** | **5.94374865180393** | **4.0442286839608** | **5.24956608126435e-05** |
| **34H-NBBr - 24H-NBBr** | **-19.8333333333334** | **5.81310085922144** | **-3.41183368629693** | **0.000645274699446652** |
| 34H-NBBr - 34H-NBCl | 1.5 | 5.81310085922144 | 0.258037841820775 | 0.796377702321007 |
| **34H-NBBr - 24H-NBCl** | **-13.25** | **5.81310085922144** | **-2.27933426941686** | **0.0226472023831299** |
| 34H-NBBr - 34H-NBF | 3.8333333333333 | 5.81310085922144 | 0.659430040208644 | 0.509619657059195 |
| 34H-NBBr - 24H-NBF | 2.99999999999995 | 5.81310085922144 | 0.516075683641543 | 0.605801550474247 |
| 34H-NBBr - 34H-NBOMe | 1.66666666666668 | 5.81310085922144 | 0.286708713134198 | 0.774335365448777 |
| 34H-NBBr - 24H-NBOMe | 2.66666666666663 | 5.81310085922143 | 0.458733941014708 | 0.646425237487486 |
| **24H-NBBr - 34H-NBCl** | **21.3333333333334** | **5.81310085922144** | **3.66987152811771** | **0.000242672400111277** |
| 24H-NBBr - 24H-NBCl | 6.58333333333334 | 5.81310085922144 | 1.13249941688007 | 0.257424529398933 |
| **24H-NBBr - 34H-NBF** | **23.6666666666667** | **5.81310085922144** | **4.07126372650558** | **4.6758774513906e-05** |
| **24H-NBBr - 24H-NBF** | **22.8333333333333** | **5.81310085922144** | **3.92790936993848** | **8.56874874207336e-05** |
| **24H-NBBr - 34H-NBOMe** | **21.5** | **5.81310085922144** | **3.69854239943114** | **0.000216841131658809** |
| **24H-NBBr - 24H-NBOMe** | **22.5** | **5.81310085922144** | **3.87056762731165** | **0.000108582228940984** |
| **34H-NBCl - 24H-NBCl** | **-14.75** | **5.81310085922144** | **-2.53737211123764** | **0.0111688167371114** |
| 34H-NBCl - 34H-NBF | 2.3333333333333 | 5.81310085922144 | 0.401392198387868 | 0.688131392381088 |
| 34H-NBCl - 24H-NBF | 1.49999999999995 | 5.81310085922144 | 0.258037841820767 | 0.796377702321013 |
| 34H-NBCl - 34H-NBOMe | 0.16666666666668 | 5.81310085922144 | 0.0286708713134219 | 0.977127088138007 |
| 34H-NBCl - 24H-NBOMe | 1.16666666666664 | 5.81310085922143 | 0.200696099193932 | 0.840936210052125 |
| **24H-NBCl - 34H-NBF** | **17.0833333333333** | **5.81310085922144** | **2.9387643096255** | **0.00329523555228239** |
| **24H-NBCl - 24H-NBF** | **16.25** | **5.81310085922144** | **2.7954099530584** | **0.00518339394617743** |
| **24H-NBCl - 34H-NBOMe** | **14.9166666666667** | **5.81310085922144** | **2.56604298255106** | **0.0102866080434138** |
| **24H-NBCl - 24H-NBOMe** | **15.9166666666667** | **5.81310085922144** | **2.73806821043157** | **0.00618012563148508** |
| 34H-NBF - 24H-NBF | -0.833333333333352 | 5.81310085922144 | -0.143354356567101 | 0.886010328757602 |
| 34H-NBF - 34H-NBOMe | -2.16666666666662 | 5.81310085922144 | -0.372721327074447 | 0.709355861713312 |
| 34H-NBF - 24H-NBOMe | -1.16666666666666 | 5.81310085922144 | -0.200696099193937 | 0.840936210052122 |
| 24H-NBF - 34H-NBOMe | -1.33333333333327 | 5.81310085922143 | -0.229366970507346 | 0.818583705600895 |
| 24H-NBF - 24H-NBOMe | -0.333333333333313 | 5.81310085922144 | -0.0573417426268356 | 0.954272969228459 |
| 34H-NBOMe - 24H-NBOMe | 0.999999999999957 | 5.81310085922142 | 0.172025227880511 | 0.863417695660923 |
| | | **Number of top entries** | | |
| 34H-NBOMe(F) - 24H-NBOMe(F) | -1.27272727272727 | 2.2494402842495 | -0.565797314842658 | 0.571531572038086 |
| **34H-NBOMe(F) - 34H-NBBr** | **-6.91666666666667** | **2.19999599834515** | **-3.14394511256812** | **0.0016668670104111** |

| | | | | |
|---|---|---|---|---|
| 34H-NBOMe(F) - 24H-NBBr | -1.58333333333334 | 2.19999599834516 | -0.719698278780656 | 0.471710786250842 |
| **34H-NBOMe(F) - 34H-NBCl** | **-9.41666666666667** | **2.19999599834515** | **-4.280310815906** | **1.86632479264106e-05** |
| 34H-NBOMe(F) - 24H-NBCl | -0.333333333333339 | 2.19999599834516 | -0.151515427111719 | 0.879569142250034 |
| 34H-NBOMe(F) - 34H-NBF | -3.58333333333334 | 2.19999599834516 | -1.62879084145096 | 0.103357304174182 |
| 34H-NBOMe(F) - 24H-NBF | -3.58333333333334 | 2.19999599834516 | -1.62879084145096 | 0.103357304174182 |
| **34H-NBOMe(F) - 34H-NBOMe** | **-7.5** | **2.19999599834515** | **-3.40909711001362** | **0.000651782735538499** |
| 34H-NBOMe(F) - 24H-NBOMe | -2.58333333333335 | 2.19999599834515 | -1.17424456011581 | 0.240297082375395 |
| **24H-NBOMe(F) - 34H-NBBr** | **-5.6439393939394** | **2.2494402842495** | **-2.50904166403441** | **0.0121059204691516** |
| 24H-NBOMe(F) - 24H-NBBr | -0.310606060606069 | 2.2494402842495 | -0.138081487550891 | 0.890176016410835 |
| **24H-NBOMe(F) - 34H-NBCl** | **-8.1439393939394** | **2.2494402842495** | **-3.62042924676107** | **0.000294114672333446** |
| 24H-NBOMe(F) - 24H-NBCl | 0.939393939393932 | 2.2494402842495 | 0.417612303812436 | 0.676230599518599 |
| 24H-NBOMe(F) - 34H-NBF | -2.31060606060607 | 2.24944028424949 | -1.02719155373221 | 0.304330274480864 |
| 24H-NBOMe(F) - 24H-NBF | -2.31060606060607 | 2.24944028424949 | -1.02719155373222 | 0.304330274480863 |
| **24H-NBOMe(F) - 34H-NBOMe** | **-6.22727272727273** | **2.24944028424949** | **-2.7683654333373** | **0.00563382458366751** |
| 24H-NBOMe(F) - 24H-NBOMe | -1.31060606060608 | 2.24944028424949 | -0.582636520641559 | 0.560138014813394 |
| **34H-NBBr - 24H-NBBr** | **5.33333333333333** | **2.19999599834515** | **2.42424683378747** | **0.0153401749608269** |
| 34H-NBBr - 34H-NBCl | -2.5 | 2.19999599834515 | -1.13636570333788 | 0.255803543272477 |
| **34H-NBBr - 24H-NBCl** | **6.58333333333333** | **2.19999599834515** | **2.9924296854564** | **0.0027676641487113** |
| 34H-NBBr - 34H-NBF | 3.33333333333333 | 2.19999599834515 | 1.51515427111716 | 0.12973334052597 |
| 34H-NBBr - 24H-NBF | 3.33333333333333 | 2.19999599834515 | 1.51515427111716 | 0.12973334052597 |
| 34H-NBBr - 34H-NBOMe | -0.583333333333327 | 2.19999599834515 | -0.265151997445501 | 0.79089237218386 |
| **34H-NBBr - 24H-NBOMe** | **4.33333333333332** | **2.19999599834515** | **1.96970055245231** | **0.0488726996426298** |
| **24H-NBBr - 34H-NBCl** | **-7.83333333333334** | **2.19999599834515** | **-3.56061253712534** | **0.000369990698362956** |
| 24H-NBBr - 24H-NBCl | 1.25 | 2.19999599834515 | 0.568182851668938 | 0.569910814278654 |
| 24H-NBBr - 34H-NBF | -2 | 2.19999599834515 | -0.9090925626703 | 0.363301268107962 |
| 24H-NBBr - 24H-NBF | -2 | 2.19999599834515 | -0.909092562670302 | 0.363301268107961 |
| **24H-NBBr - 34H-NBOMe** | **-5.91666666666666** | **2.19999599834515** | **-2.68939883123297** | **0.00715808405919645** |
| 24H-NBBr - 24H-NBOMe | -1.00000000000002 | 2.19999599834515 | -0.454546281335157 | 0.649435688791224 |
| **34H-NBCl - 24H-NBCl** | **9.08333333333334** | **2.19999599834515** | **4.12879538879428** | **3.64668733892553e-05** |
| **34H-NBCl - 34H-NBF** | **5.83333333333333** | **2.19999599834515** | **2.65151997445504** | **0.0080130373205998** |
| **34H-NBCl - 24H-NBF** | **5.83333333333333** | **2.19999599834515** | **2.65151997445504** | **0.00801303732059983** |
| 34H-NBCl - 34H-NBOMe | 1.91666666666668 | 2.19999599834515 | 0.871213705892375 | 0.383637477680229 |
| **34H-NBCl - 24H-NBOMe** | **6.83333333333332** | **2.19999599834515** | **3.10606625579019** | **0.00189594229418652** |
| 24H-NBCl - 34H-NBF | -3.25 | 2.19999599834515 | -1.47727541433924 | 0.139601825566451 |
| 24H-NBCl - 24H-NBF | -3.25000000000001 | 2.19999599834515 | -1.47727541433924 | 0.139601825566451 |
| **24H-NBCl - 34H-NBOMe** | **-7.16666666666666** | **2.19999599834515** | **-3.25758168290191** | **0.00112365914100504** |
| 24H-NBCl - 24H-NBOMe | -2.25000000000002 | 2.19999599834515 | -1.02272913300409 | 0.306435932927541 |
| 34H-NBF - 24H-NBF | -1.55431223447522e-15 | 2.19999599834515 | -7.06506846214438e-16 | 0.999999999999999 |
| 34H-NBF - 34H-NBOMe | -3.91666666666666 | 2.19999599834515 | -1.78030626856267 | 0.0750258518070118 |
| 34H-NBF - 24H-NBOMe | 0.999999999999988 | 2.19999599834515 | 0.454546281335144 | 0.649435688791233 |
| 24H-NBF - 34H-NBOMe | -3.91666666666665 | 2.19999599834515 | -1.78030626856267 | 0.0750258518070115 |
| 24H-NBF - 24H-NBOMe | 0.999999999999989 | 2.19999599834515 | 0.454546281335145 | 0.649435688791233 |
| **34H-NBOMe - 24H-NBOMe** | **4.91666666666664** | **2.19999599834515** | **2.23485254989782** | **0.0254270305729397** |

**Supplementary Table S9.** Behavioral alterations of adult zebrafish in the Zebrafish Tail Immobilization test induced by anxiolytic *N*-benzylphenethylamine deriatives 24H- and 34H-NBOMe(F) assessed using Mann-Whitney test. Data is represented as mean ± S.E.M. (n=13-15). Also see Figure 7 in the main text for graphical representation.

| Endpoint | MW statistics | p | Control | Experimental group |
|---|---|---|---|---|
| | | 24H-NBOMe(F) | | |
| Distance, cm | 51 | 0,0908 | 46,78±9,718 | 65,65±10,45 |
| **Time spent mobile, s** | **45** | **0,0441** | **10,11±1,621** | **22,91±4,687** |
| | | 34H-NBOMe(F) | | |
| **Distance, cm** | **57** | **0,0209** | **70,21±20,59** | **127±27,23** |
| **Time spent mobile, s** | **49** | **0,0075** | **26,6±12,68** | **54,82±12,7** |

**Supplementary Table S10.** Results of k-means clustering analyses (4 centers) on standardized means of velocity, time spent top and horizontal alterations relative to velocity of all groups (including control reference group) in GZLM experiment. Also see Figure 6 in the main text for graphical representation.

| Cluster name | Velocity, cm/s | Time spent top, s | The horizontal shuttling behavior | Within cluster sum of squares by cluster | Cluster members |
|---|---|---|---|---|---|
| **A** | -1.10214363156356 | -0.771653546720937 | -0.687803077734166 | 0.795781981955976 | 24H-NBF, 24H-NBOMe, 34H-NBF |
| **B** | 0.158128889284927 | 0.439603860661689 | -0.664406034442939 | 1.02724218143932 | Control, 34H-NBBr, 34H-NBCl, 34H-NBOMe |
| **C** | 1.06099372354873 | -0.624057515394331 | 1.31492188652732 | 2.19554600340698 | 24H-NBBr, 24H-NBCl, 24H-NBOMe(F) |
| **D** | -0.50906583309522 | 2.42871774369905 | 0.776267711392304 | 0 | 34H-NBOMe(F) |

**Analytical data**

**24H-NBOMe (F) imine (23).**

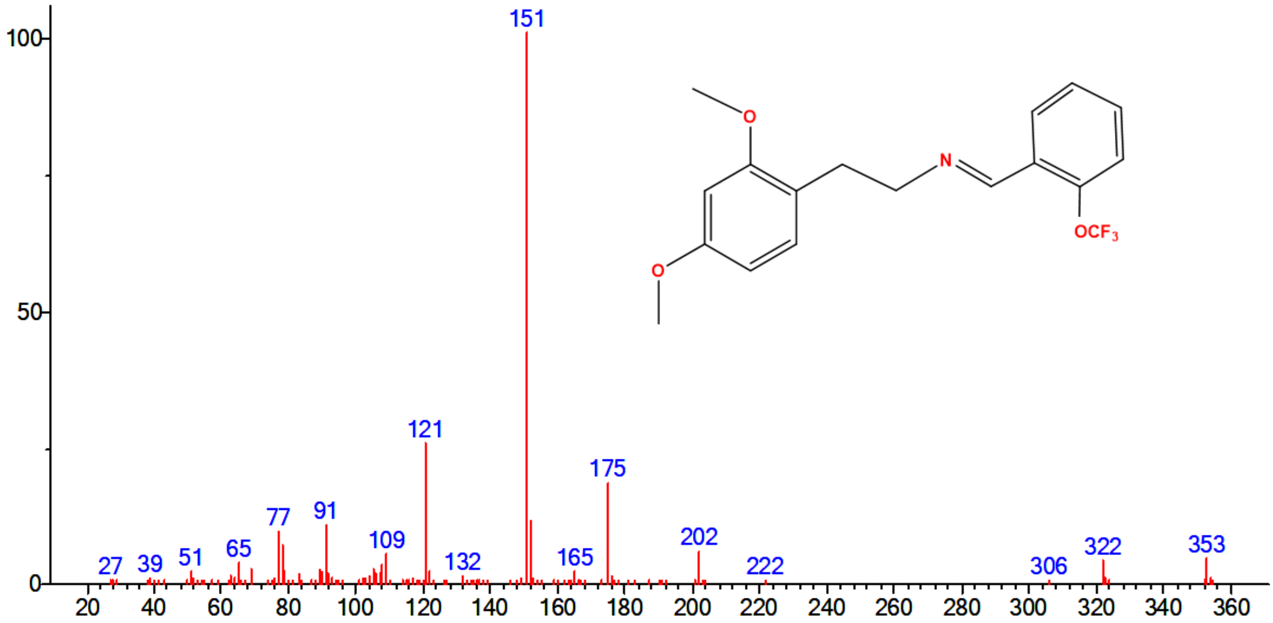


**Figure S1:** EI mass spectrum of **23**

**24H-NBF imine (24).**

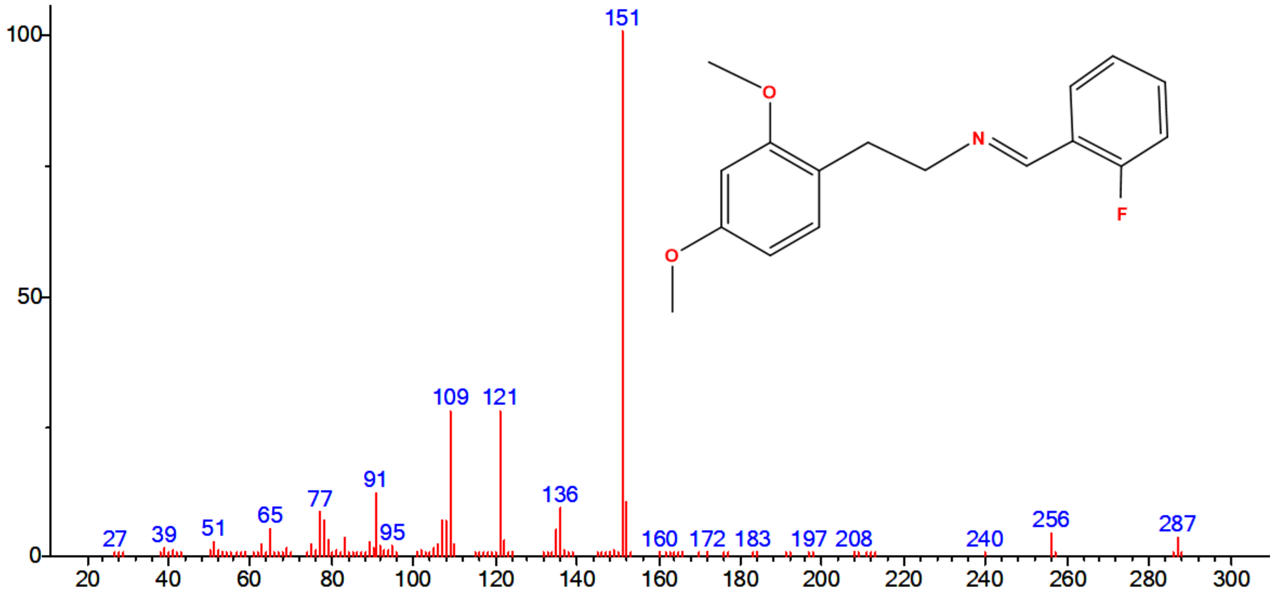


**Figure S2:** EI mass spectrum of **24**

**24H-NBCl imine (25).**

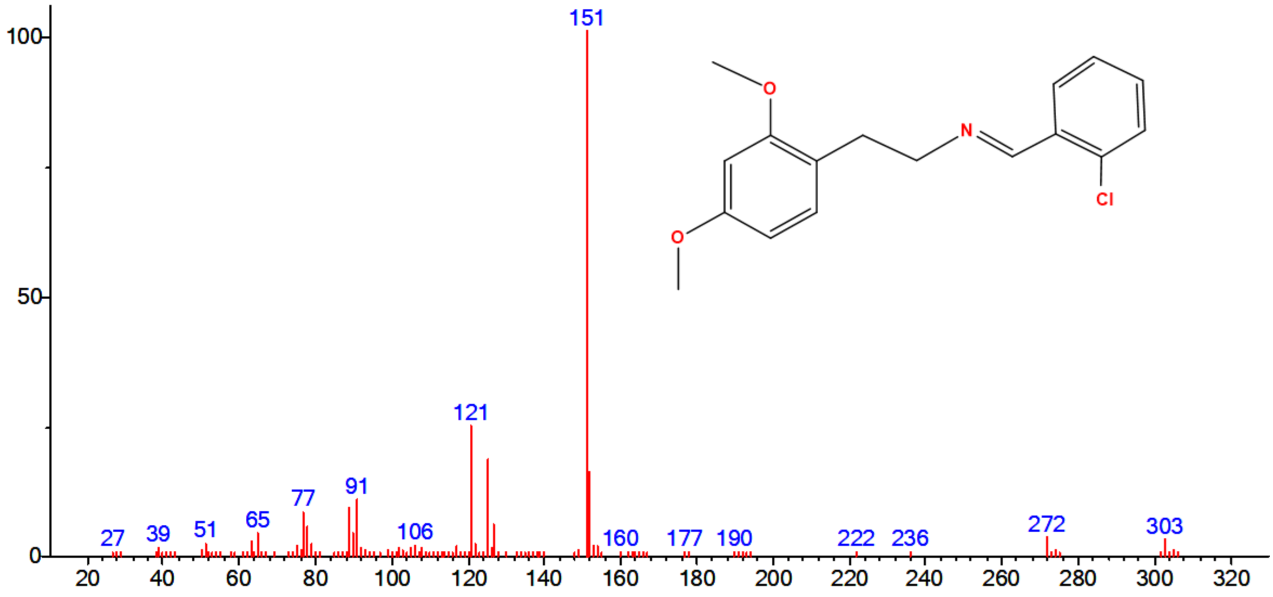


**Figure S3:** EI mass spectrum of **25**

**24H-NBBr imine (26).**

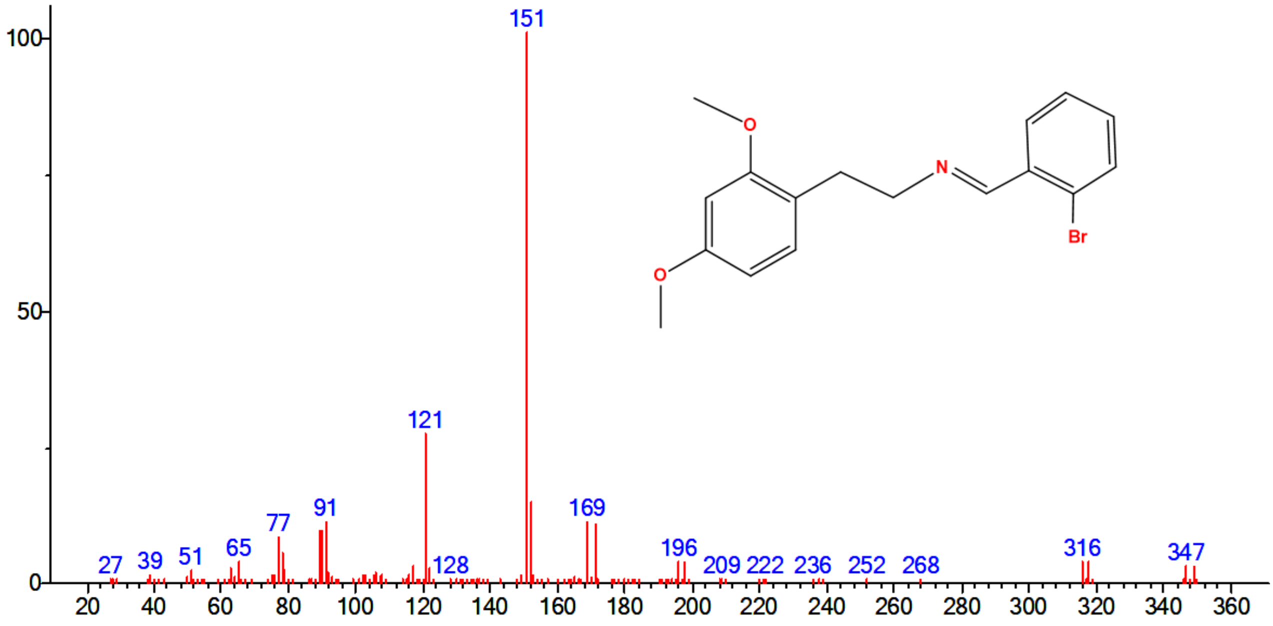


**Figure S4:** EI mass spectrum of **26**

**34H-NBOMe (F) imine (28).**

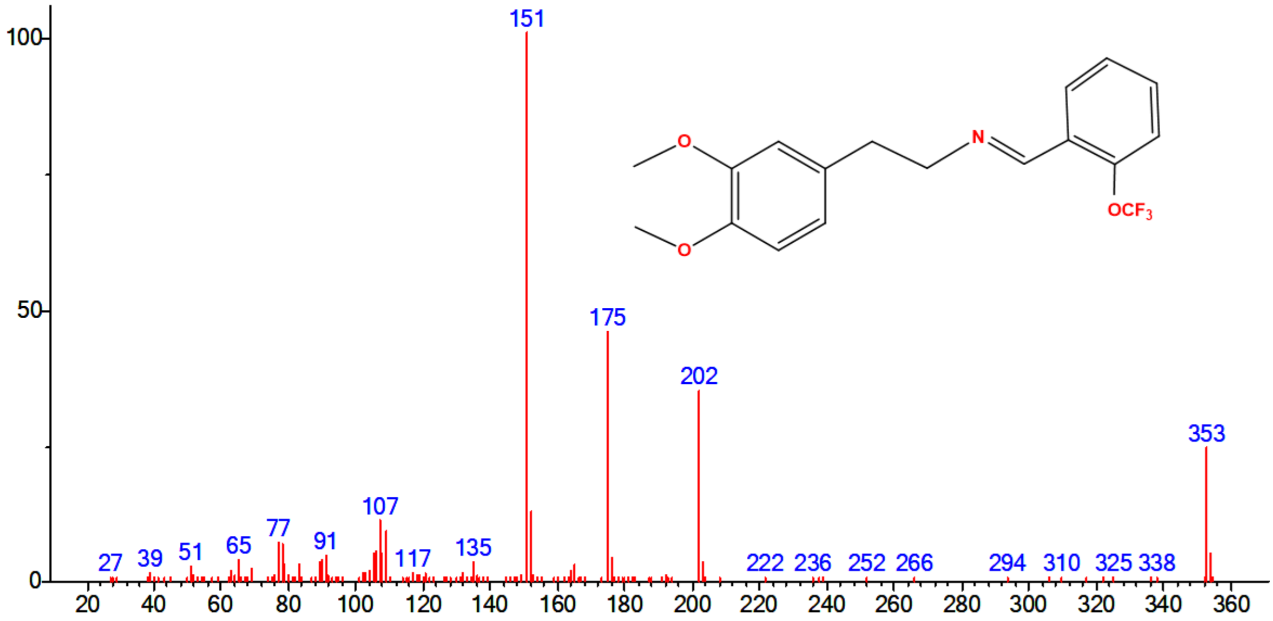


**Figure S5:** EI mass spectrum of **28**

**34H-NBF imine (29).**

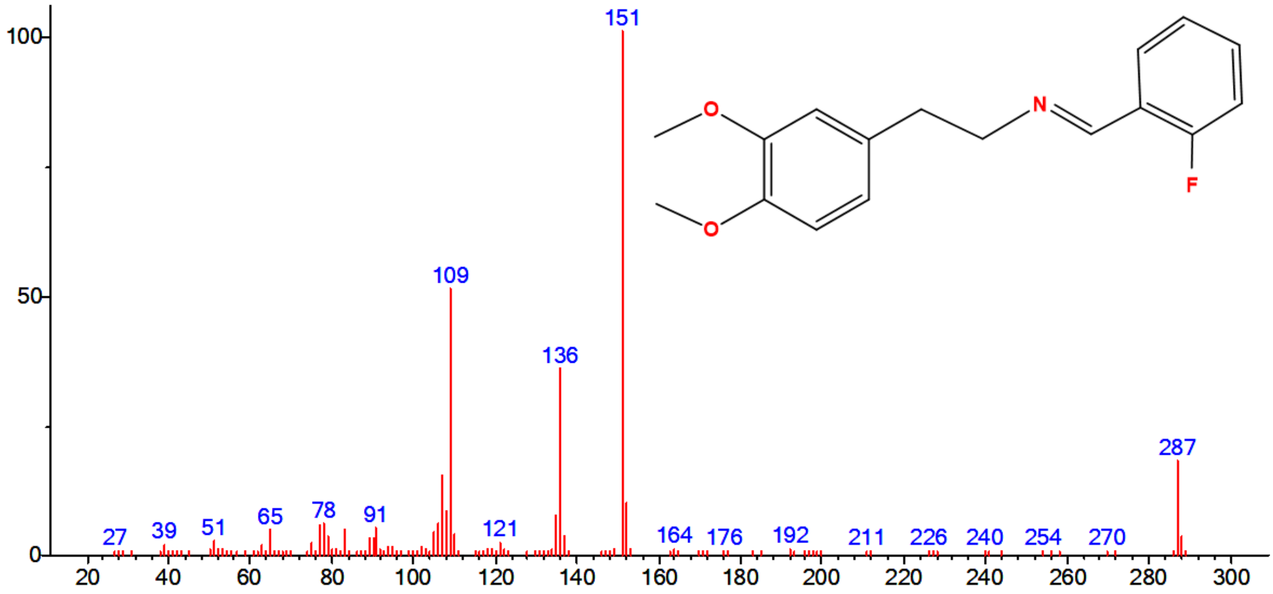


**Figure S6:** EI mass spectrum of **29**

**34H-NBCl imine (30).**

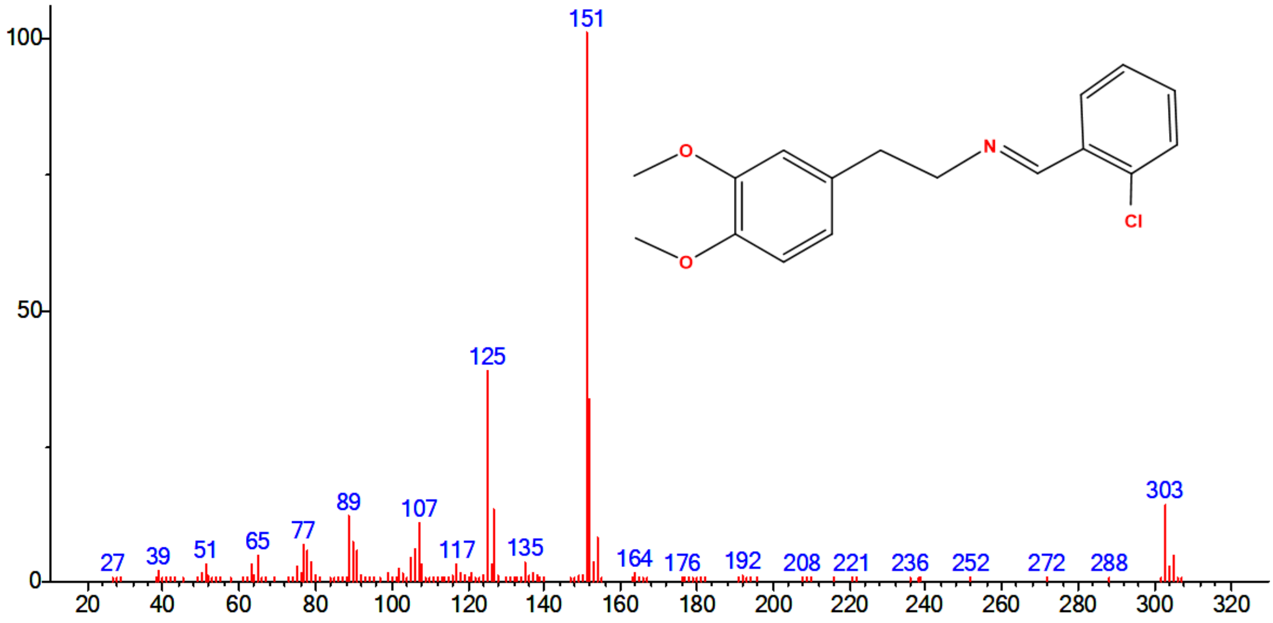


**Figure S7:** EI mass spectrum of **30**

**34H-NBBr imine (31).**

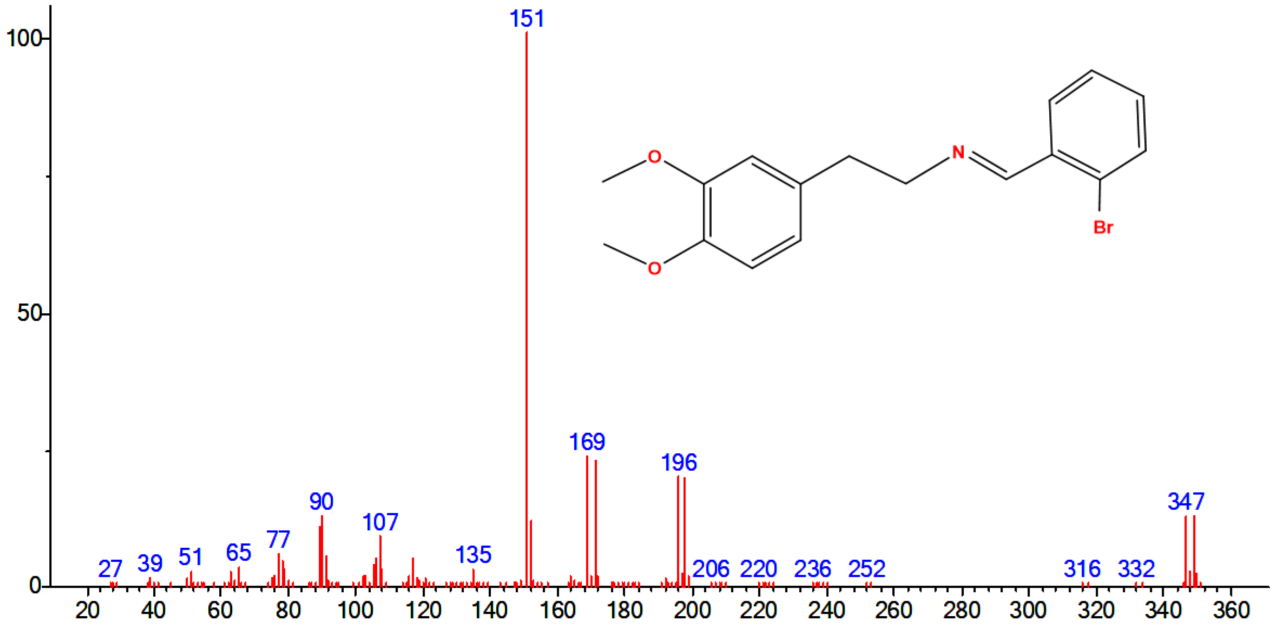


**Figure S8:** EI mass spectrum of **31**

***N*-(2-Trifluoromethoxybenzyl)-2-(2,4-dimethoxyphenyl)ethanamine hydrochloride (24H-NBOMe(F), (2)).**

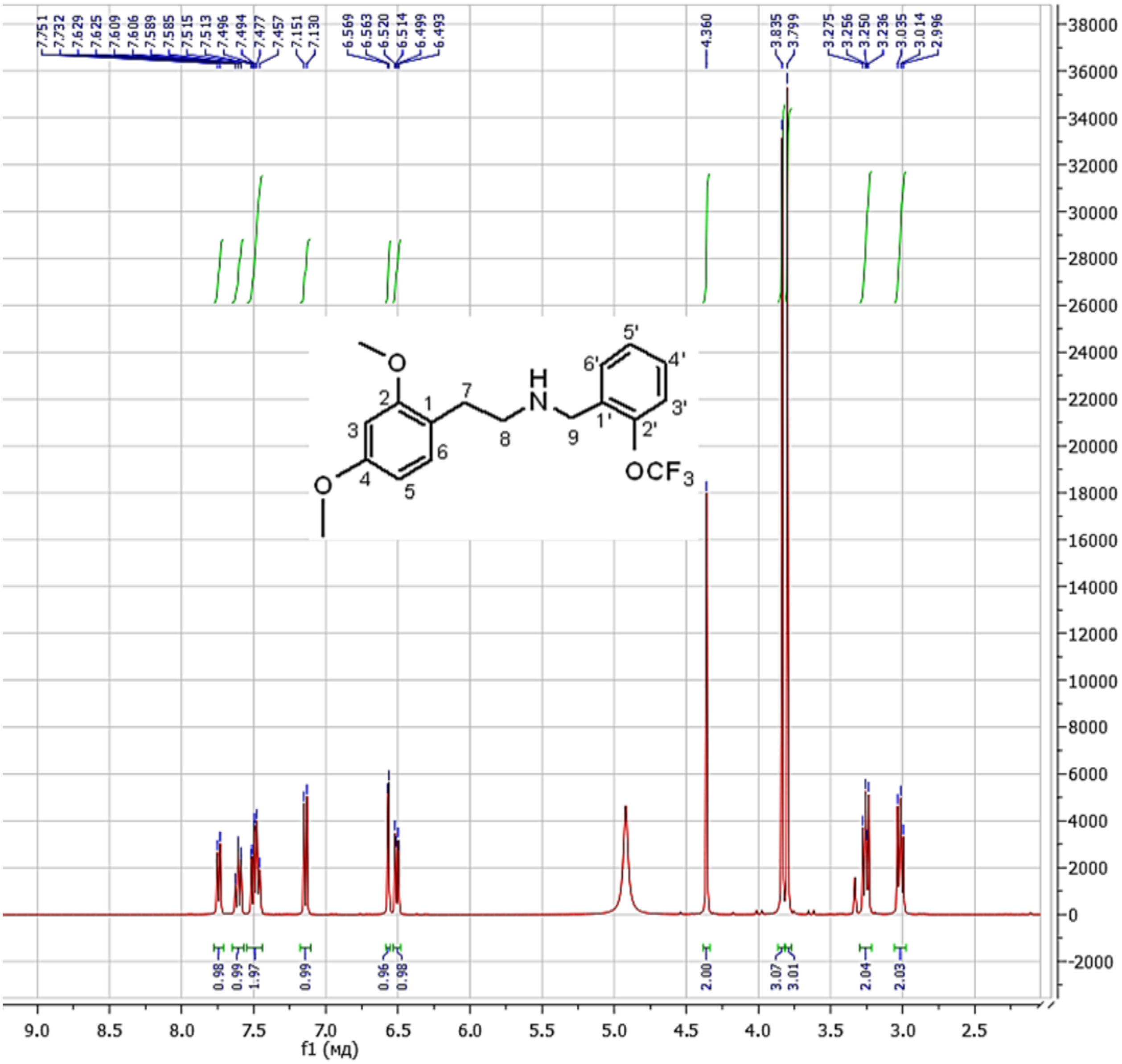


**Figure S9.** $^{1}$H NMR spectrum of **2** (400 MHz, $CD_3OD$)

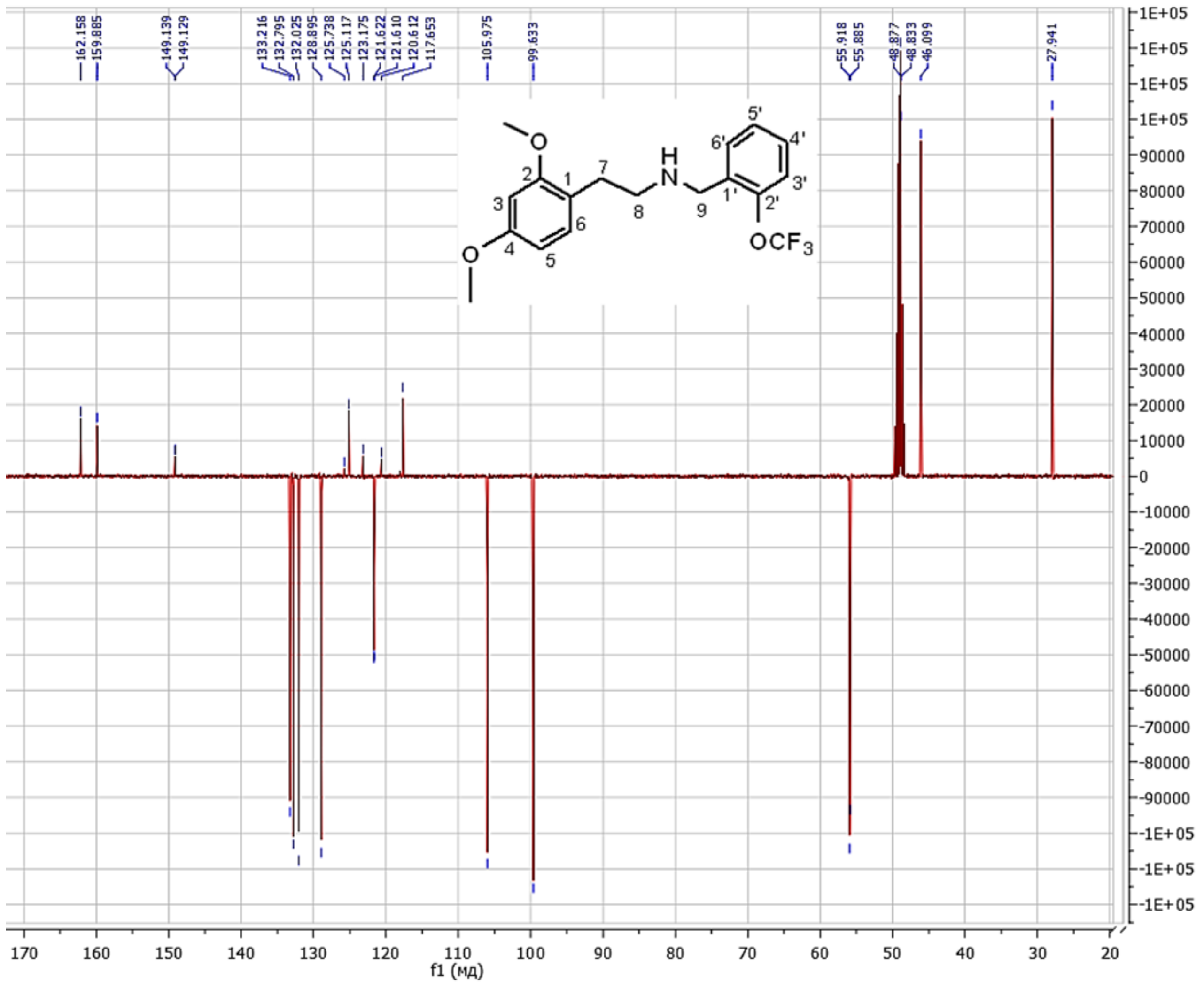


**Figure S10.** [13]C NMR spectrum of **2** (101 MHz, $CD_3OD$)

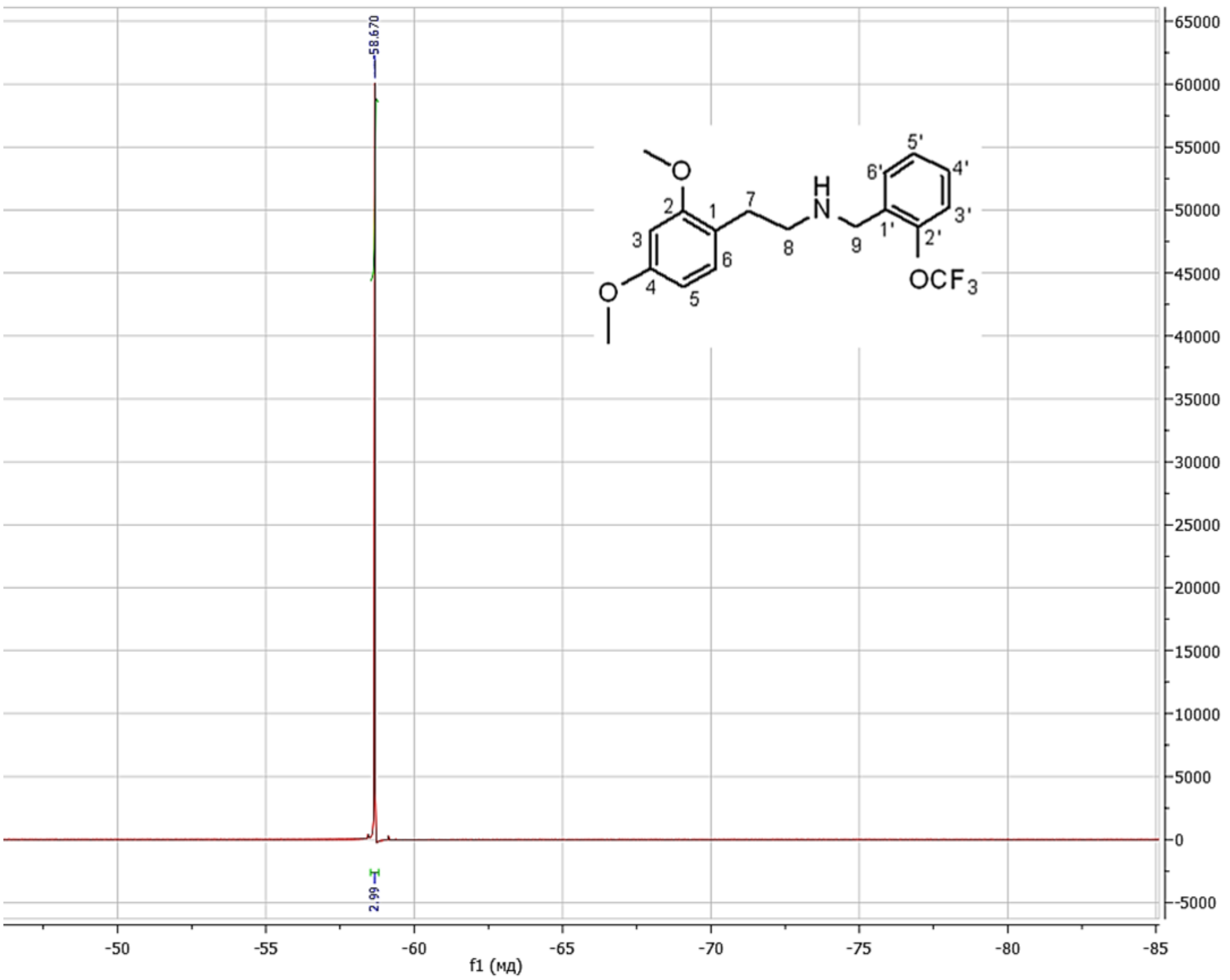


**Figure S11.** $^{19}$F NMR spectrum of **2** (376 MHz, $CD_3OD$)

**N-(2-Fluorobenzyl)-2-(2,4-dimethoxyphenyl)ethanamine hydrochloride (24H-NBF, (3)).**

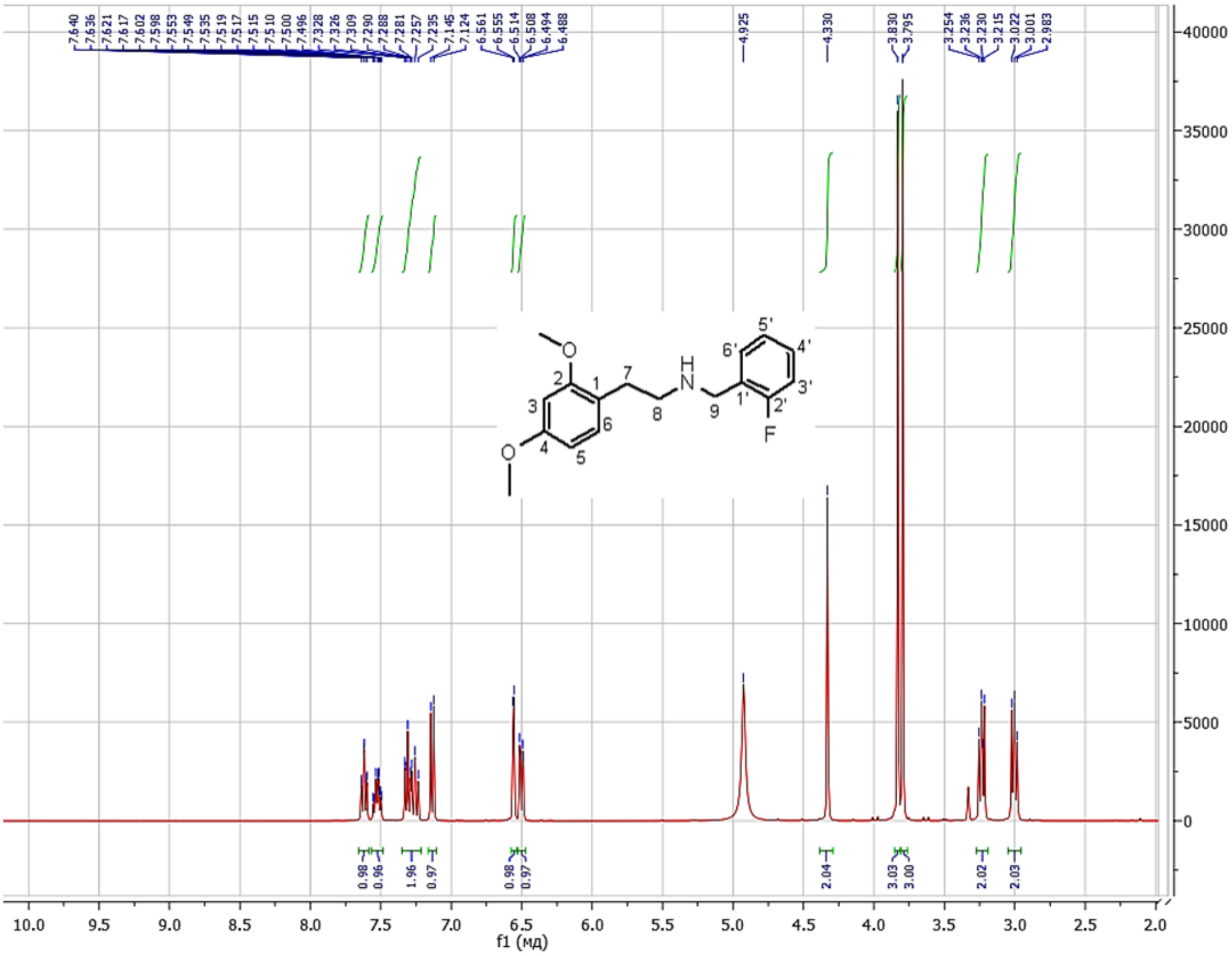

**Figure S12.** $^{1}H$ NMR spectrum of **3** (400 MHz, $CD_3OD$)

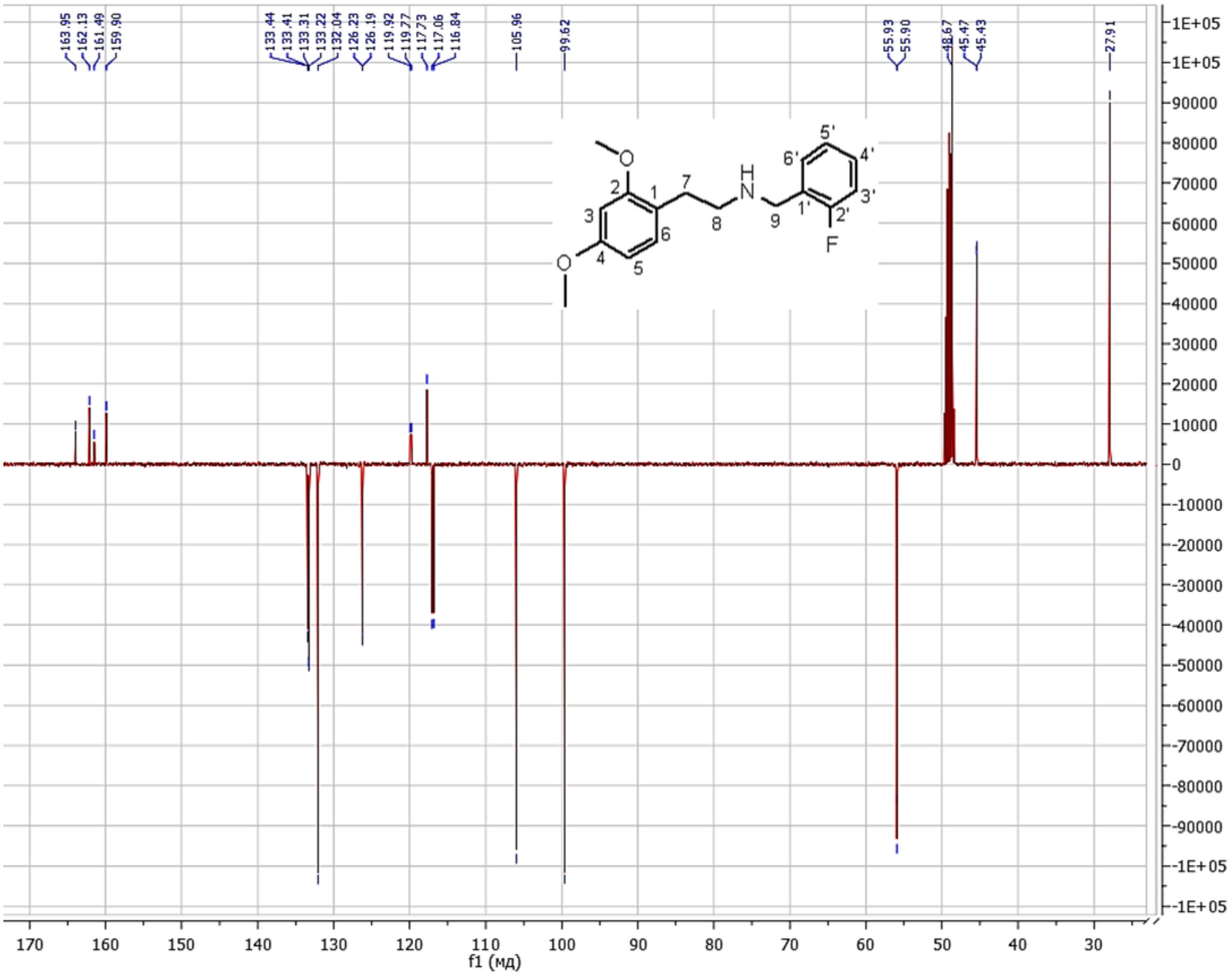


**Figure S13.** $^{13}$C NMR spectrum of **3** (101 MHz, $CD_3OD$)

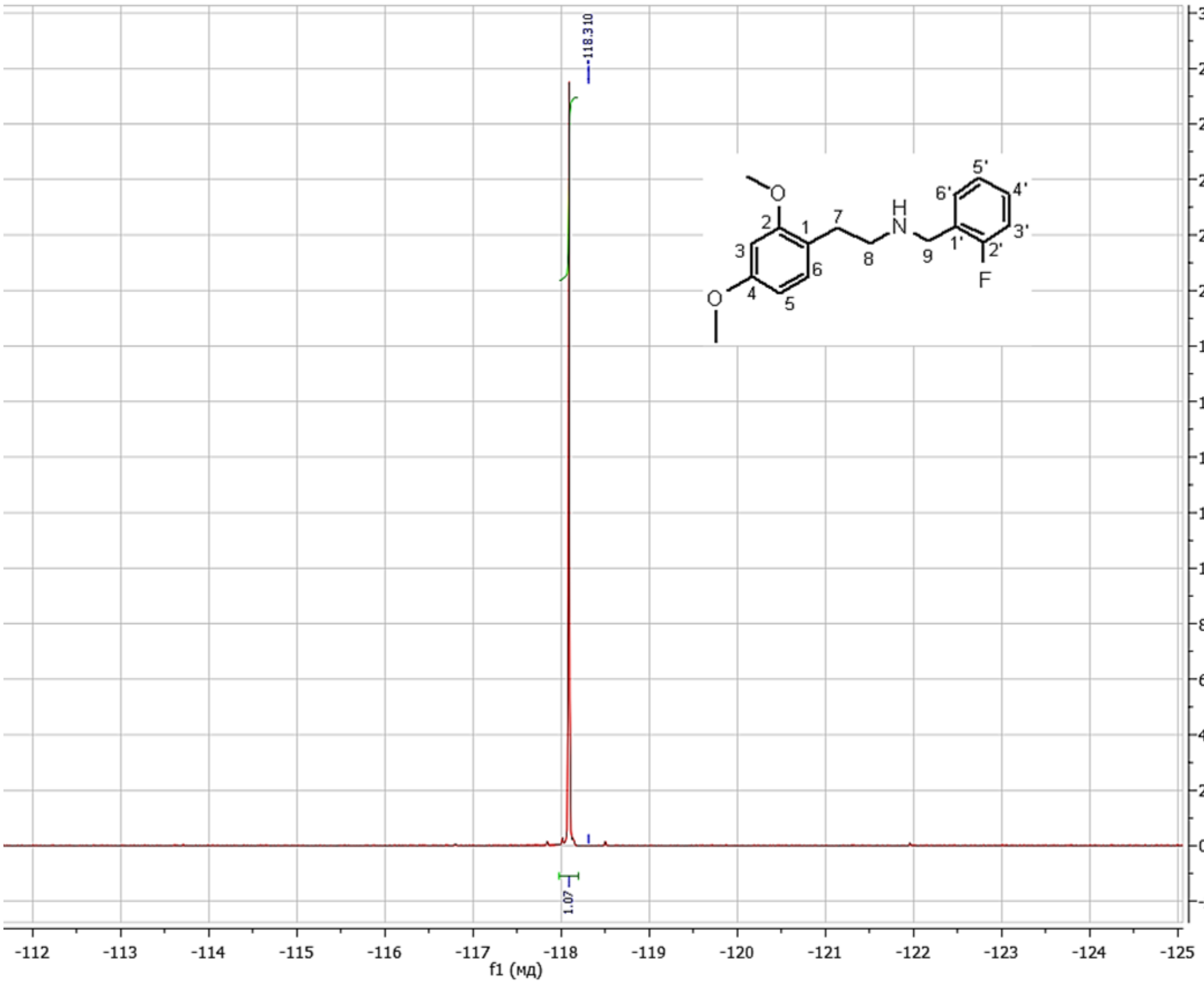


**Figure S14.** $^{19}$F NMR spectrum of **3** (376 MHz, $CD_3OD$)

**_N_-(2-Chlorobenzyl)-2-(2,4-dimethoxyphenyl)ethanamine hydrochloride (24H-NBCl, (4)).**

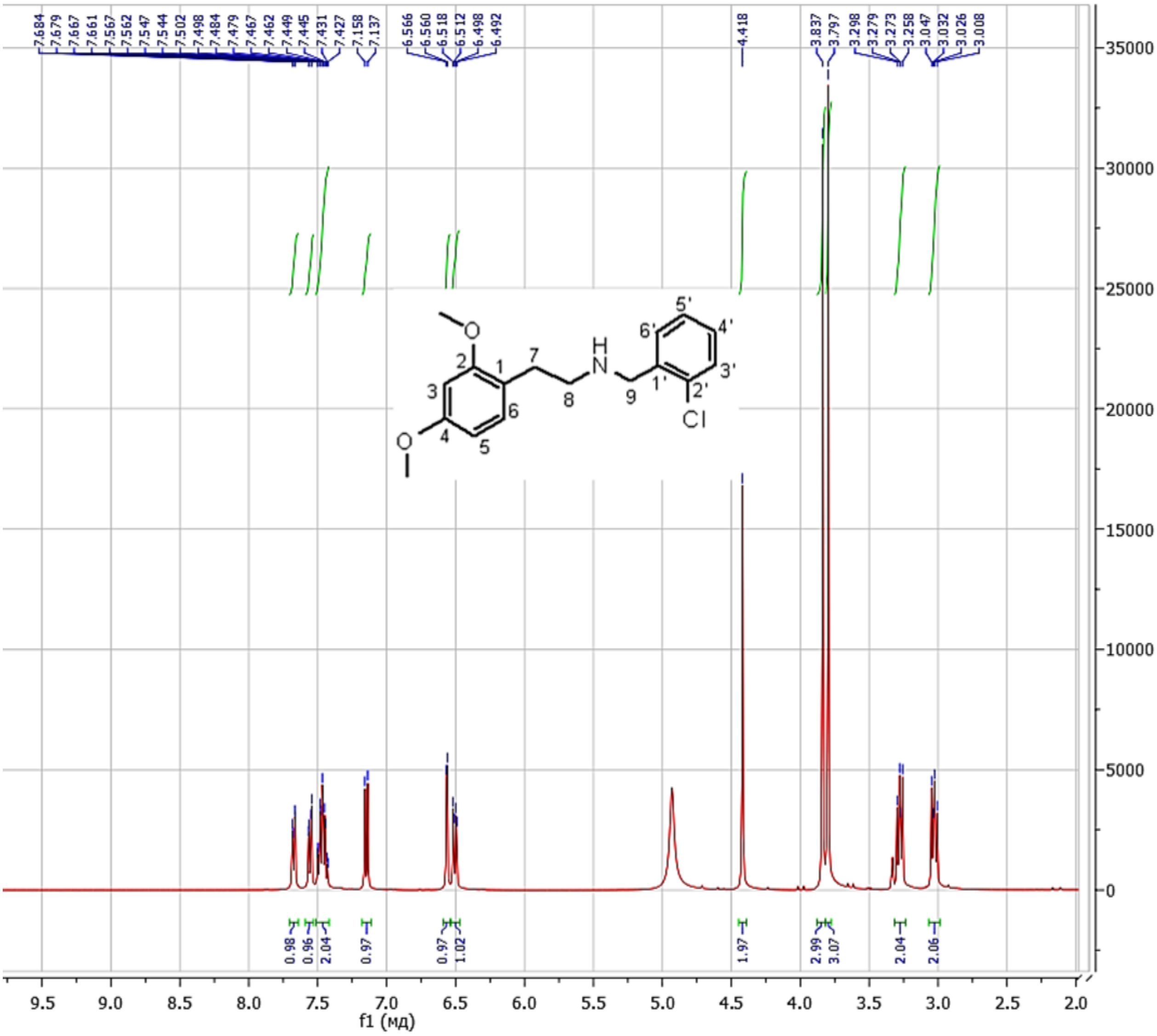


**Figure S15.** $^{1}$H NMR spectrum of **4** (400 MHz, $CD_3OD$)

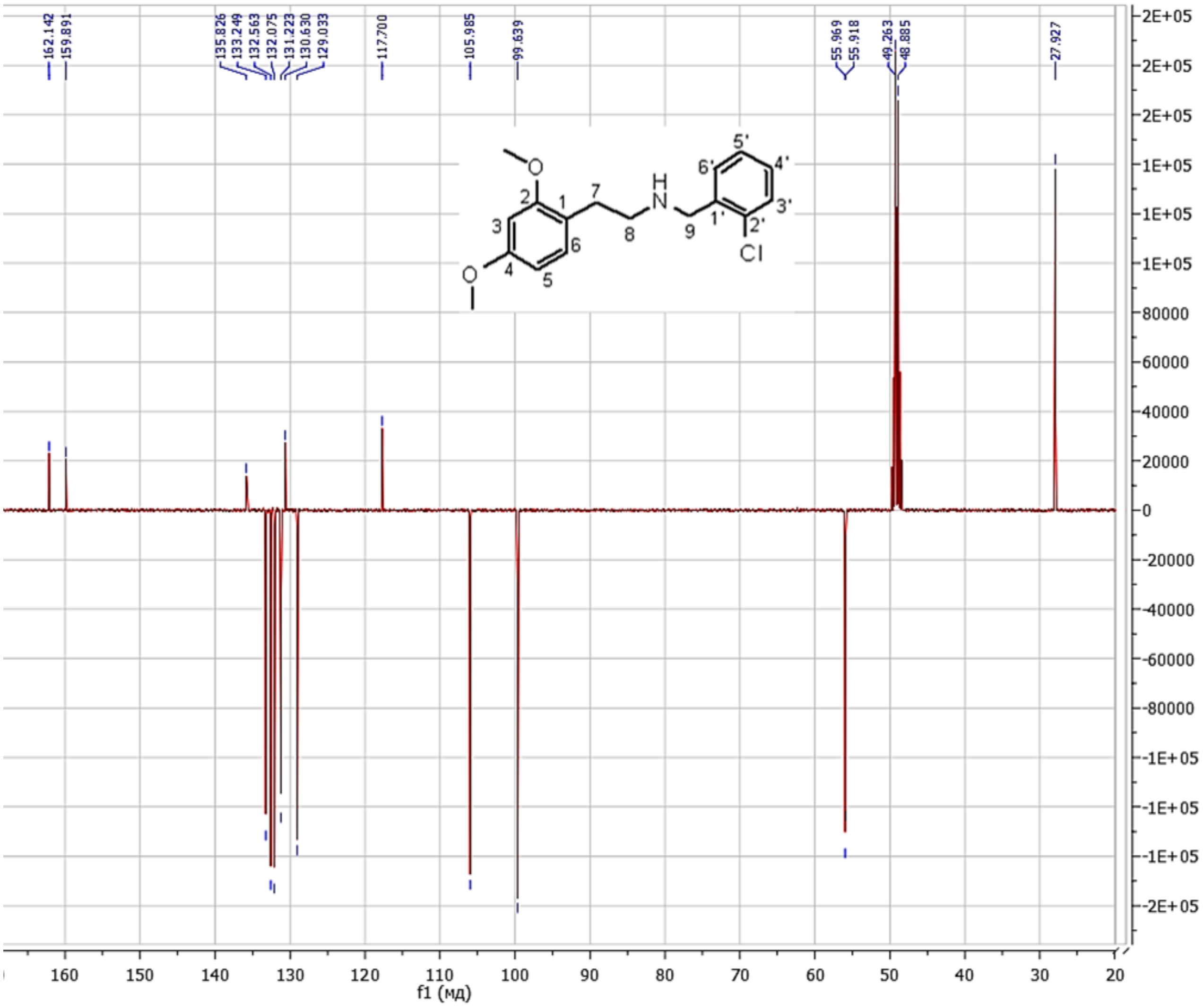


**Figure S16.** $^{13}C$ NMR spectrum of **4** (101 MHz, $CD_3OD$)

**N-(2-Bromobenzyl)-2-(2,4-dimethoxyphenyl)ethanamine hydrochloride (24H-NBBr, (5)).**

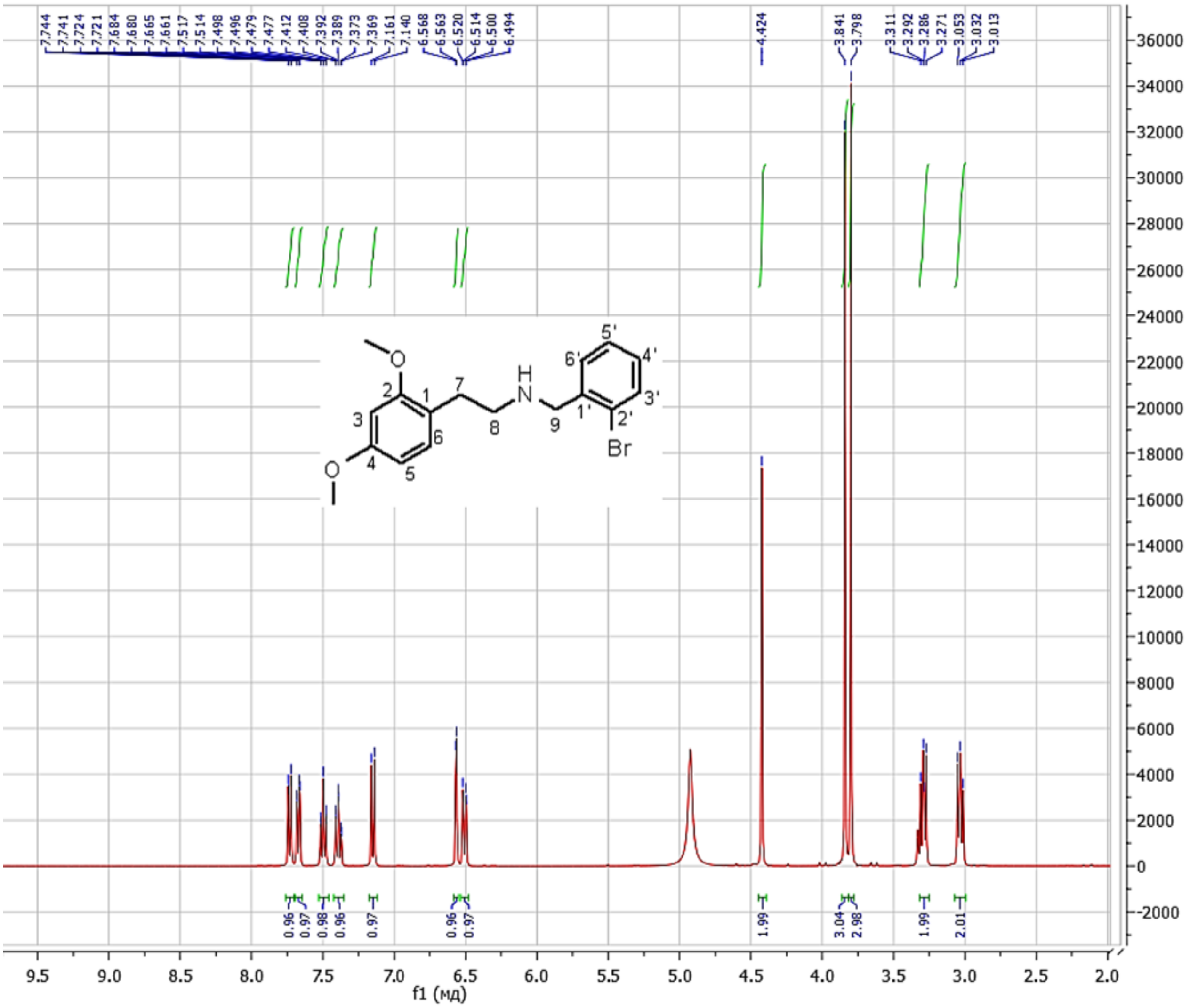


**Figure S17.** $^{1}$H NMR spectrum of **5** (400 MHz, $CD_3OD$)

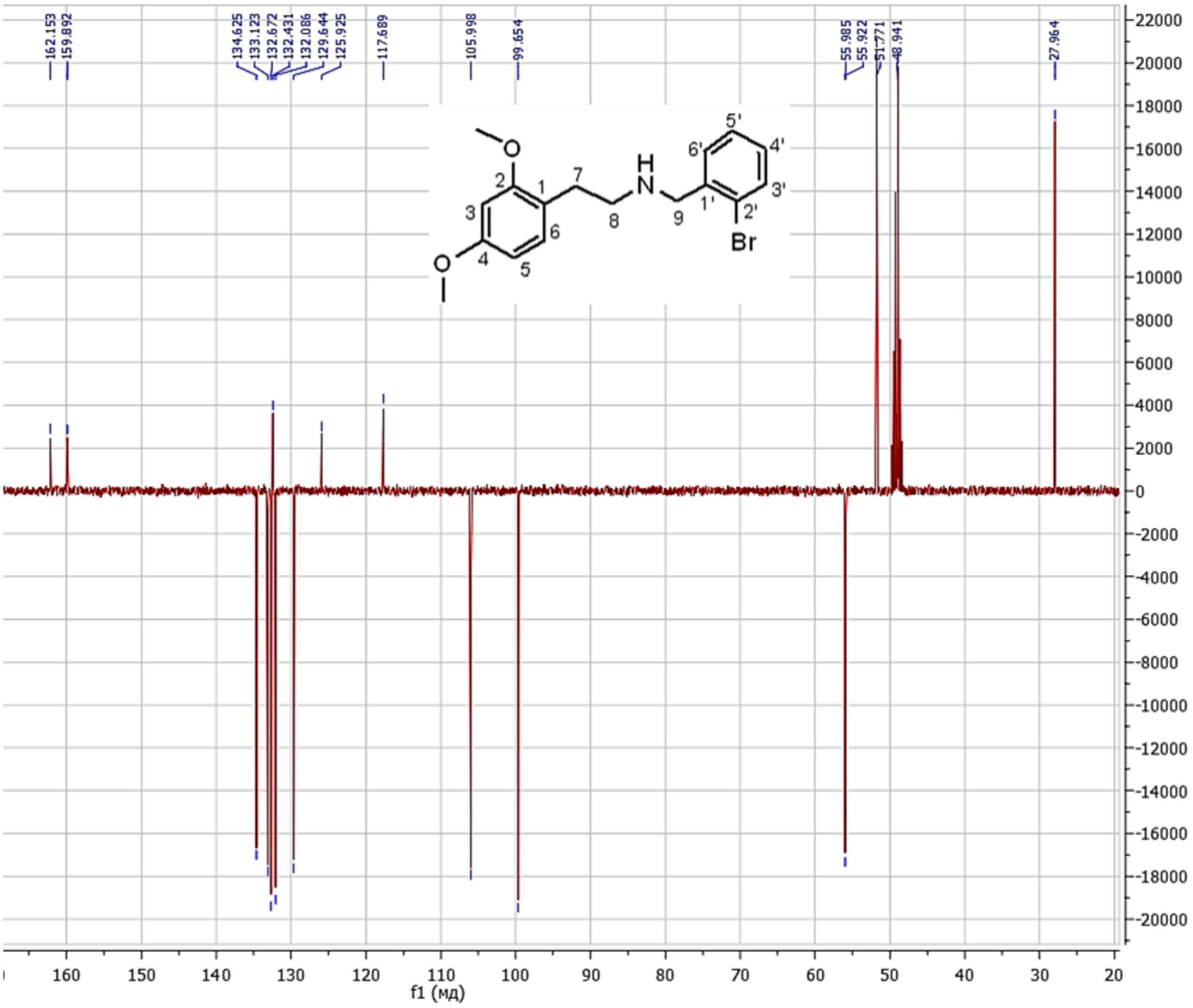


**Figure S18.** $^{13}$C NMR spectrum of **5** (101 MHz, $CD_3OD$)

***N*-(2-Trifluoromethoxybenzyl)-2-(3,4-dimethoxyphenyl)ethanamine hydrochloride (34H-NBOMe(F), (7)).**

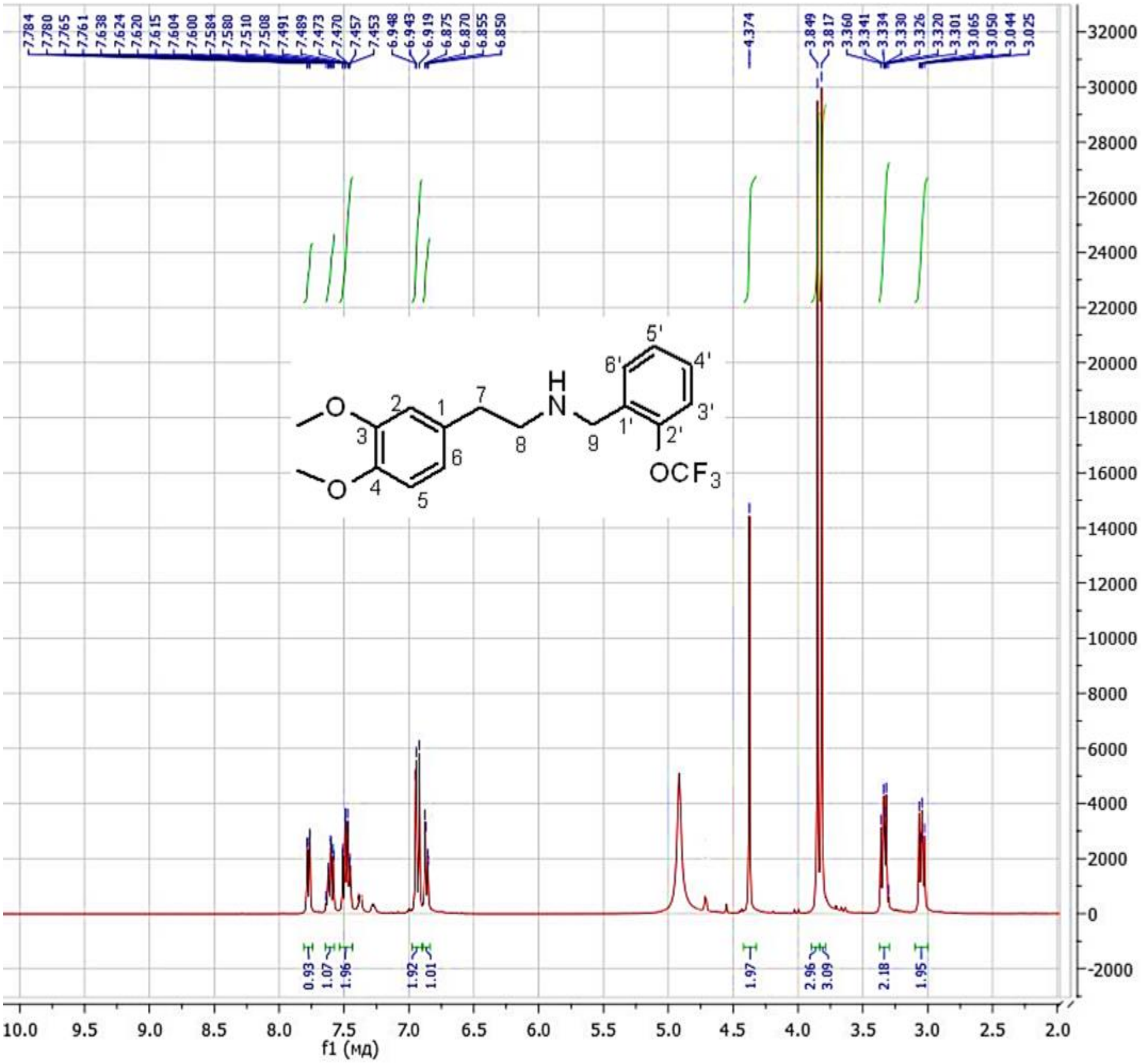


**Figure S19.** $^{1}$H NMR spectrum of **7** (400 MHz, $CD_3OD$)

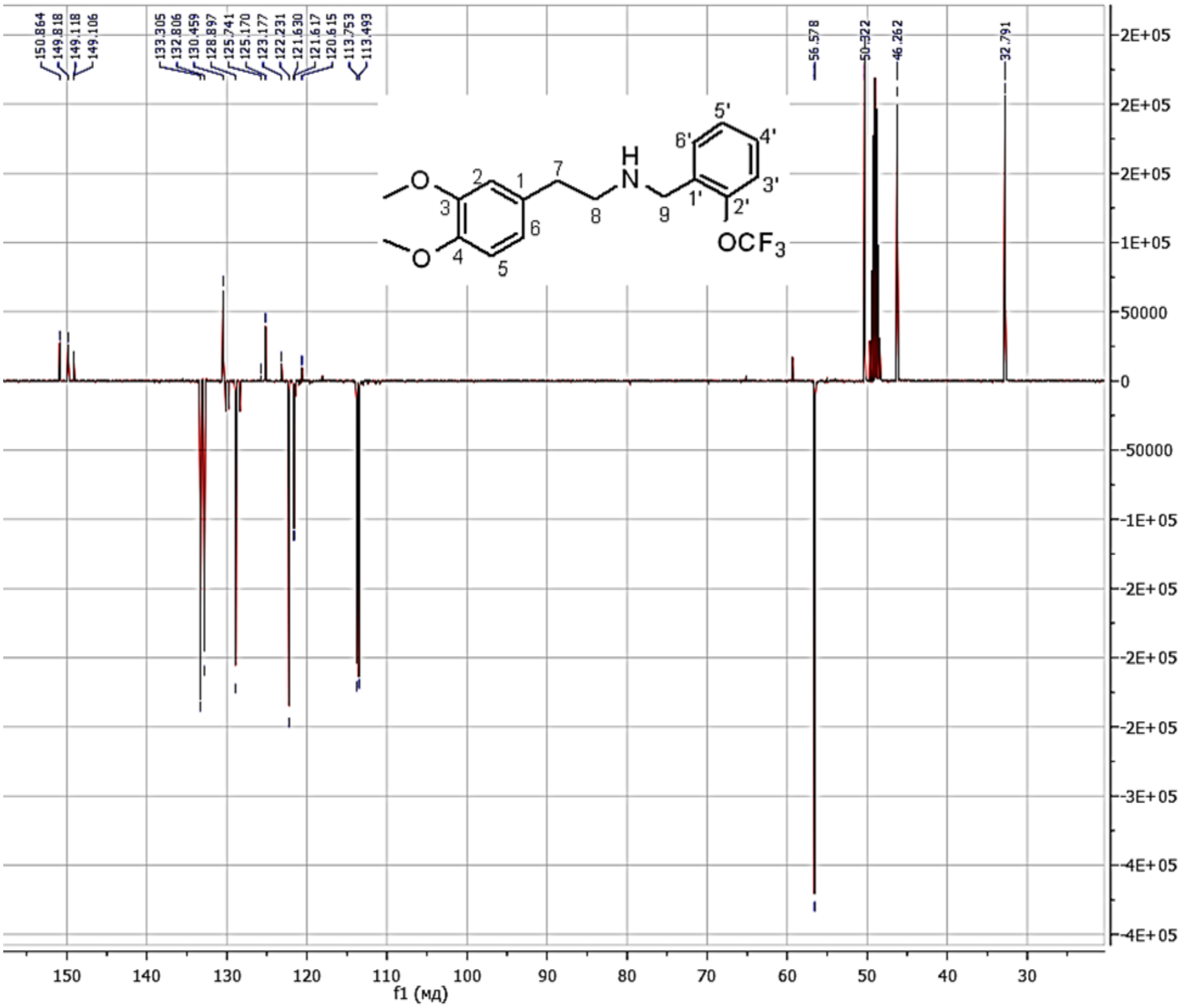

**Figure S20.** $^{13}C$ NMR spectrum of **7** (101 MHz, $CD_3OD$)

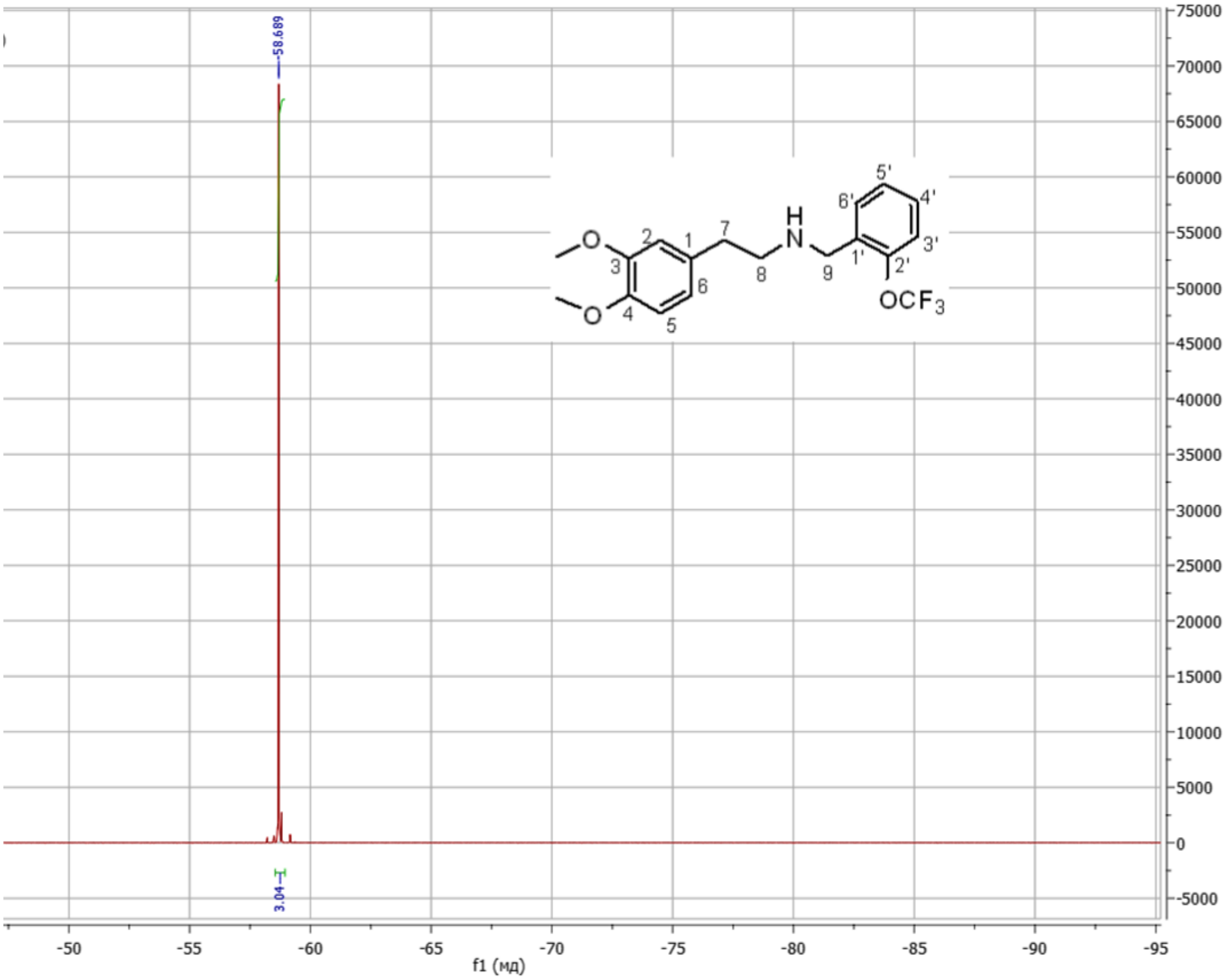


**Figure S21.** [19]F NMR spectrum of **7** (376 MHz, $CD_3OD$)

***N*-(2-Fluorobenzyl)-2-(3,4-dimethoxyphenyl)ethanamine hydrochloride (34H-NBF, (8)).**

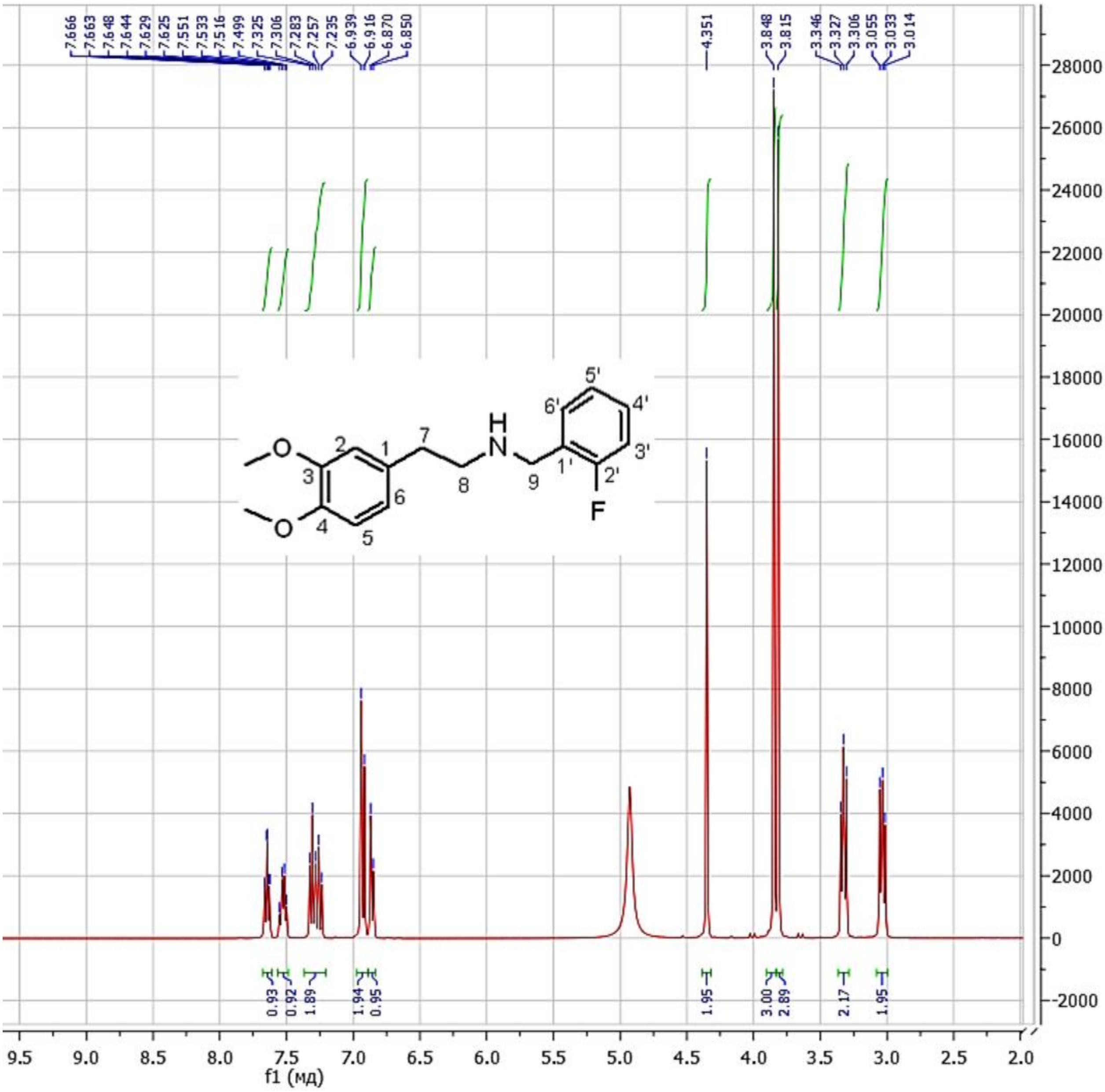

**Figure S22.** [1]H NMR spectrum of **8** (400 MHz, $CD_3OD$)

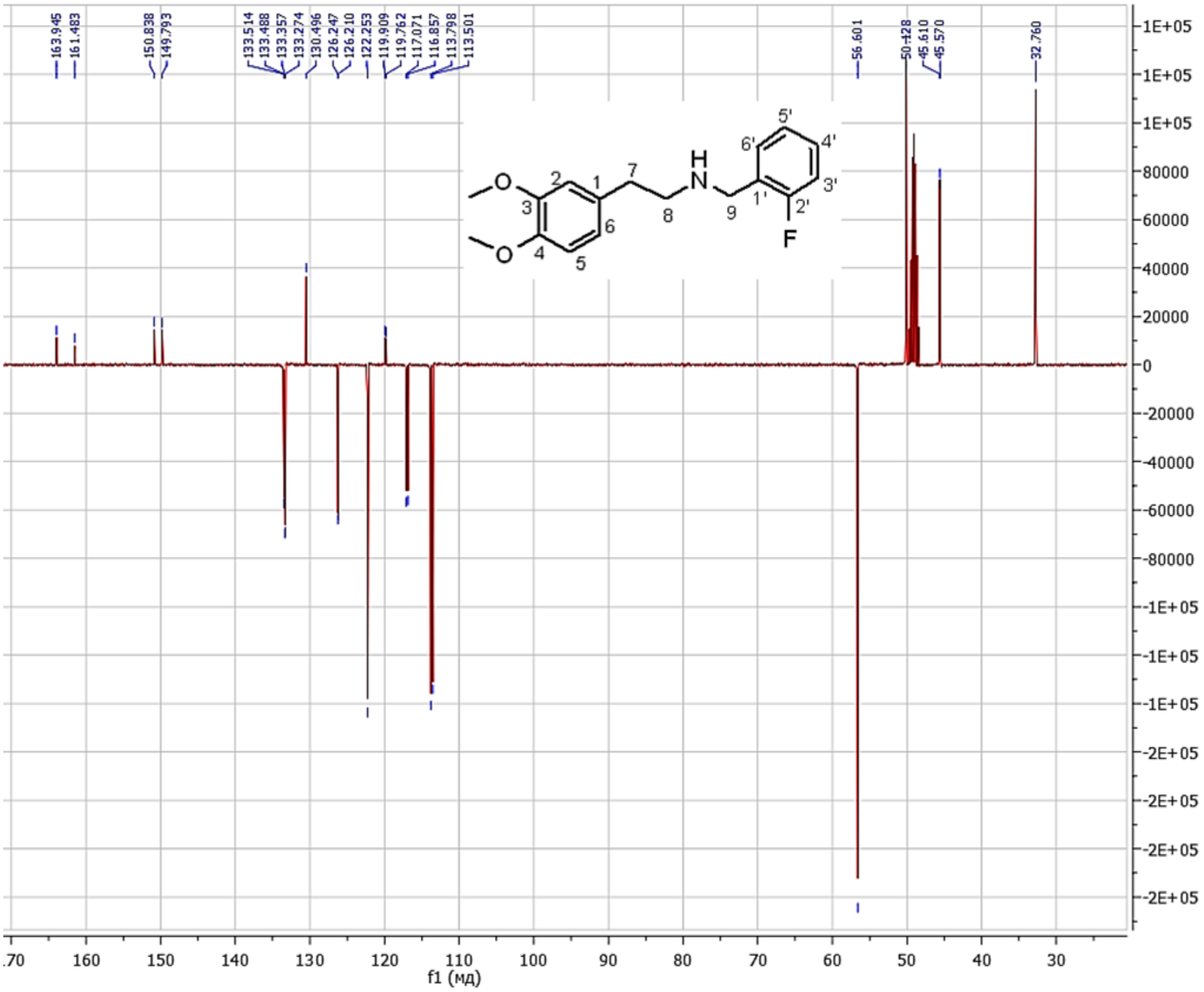


**Figure S23.** $^{13}$C NMR spectrum of **8** (101 MHz, $CD_3OD$)

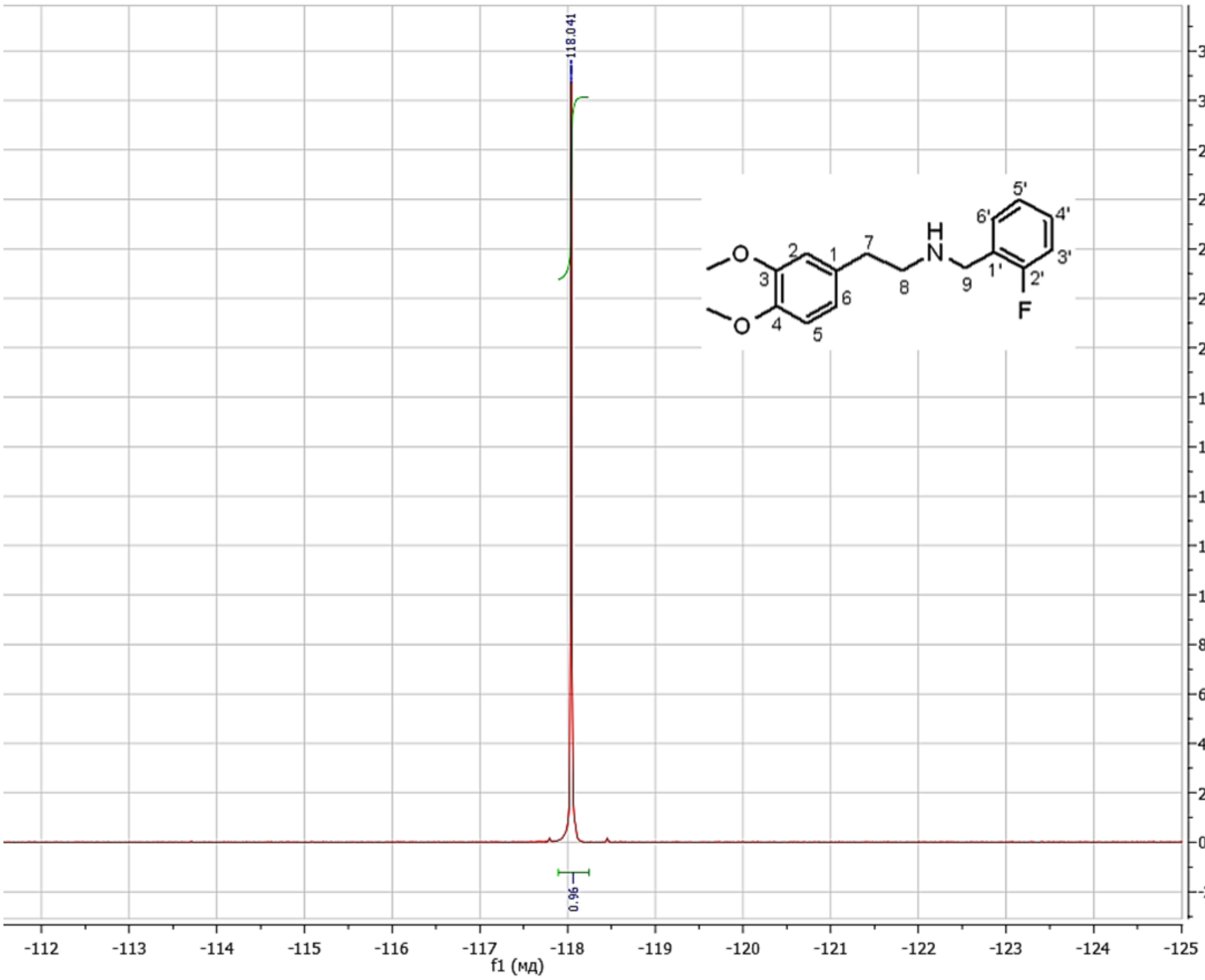


**Figure S24.** $^{19}$F NMR spectrum of **8** (376 MHz, $CD_3OD$)

**N-(2-Chlorobenzyl)-2-(3,4-dimethoxyphenyl)ethanamine hydrochloride (34H-NBCl, (9)).**

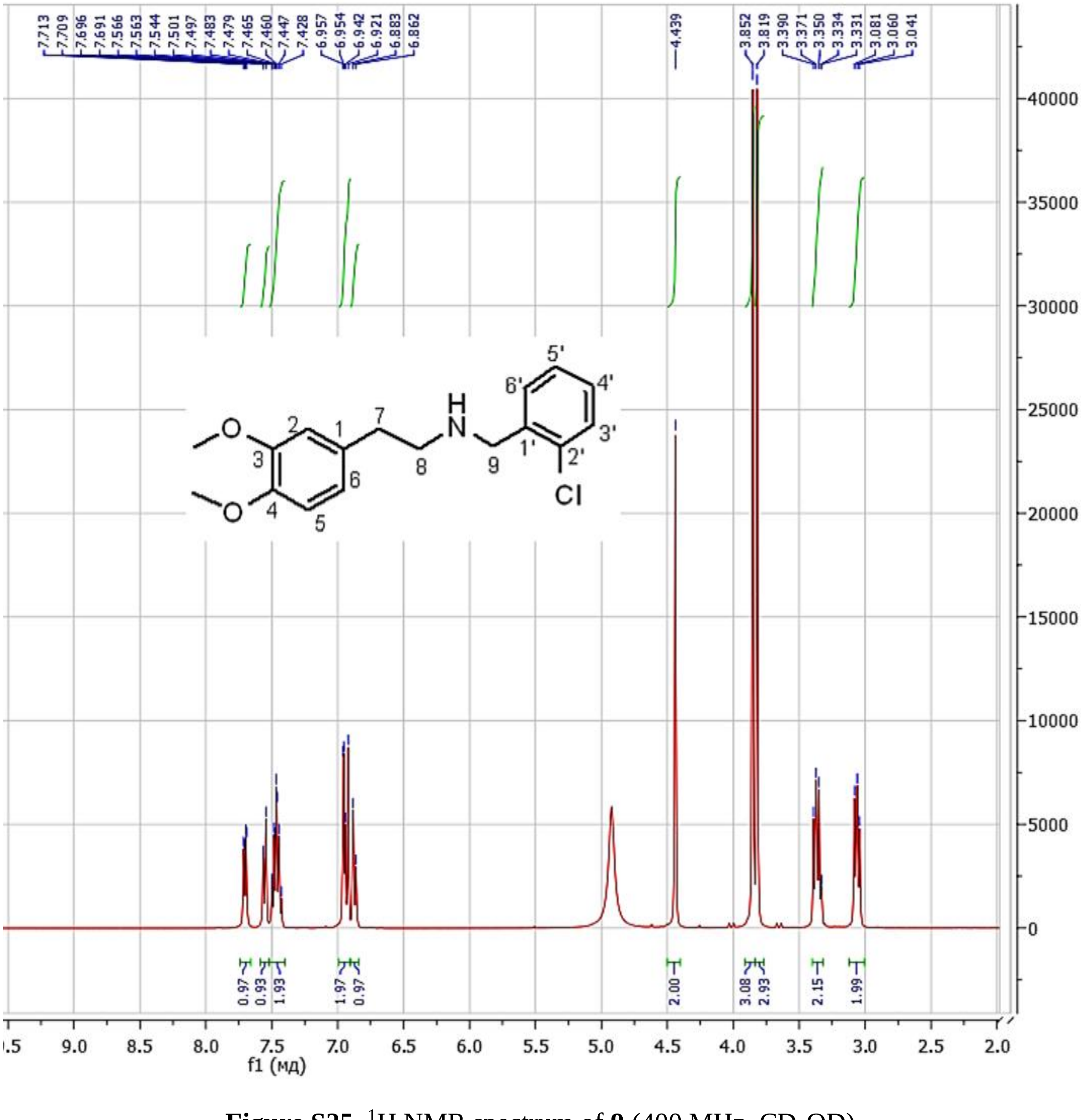

**Figure S25.** $^{1}H$ NMR spectrum of **9** (400 MHz, $CD_3OD$)

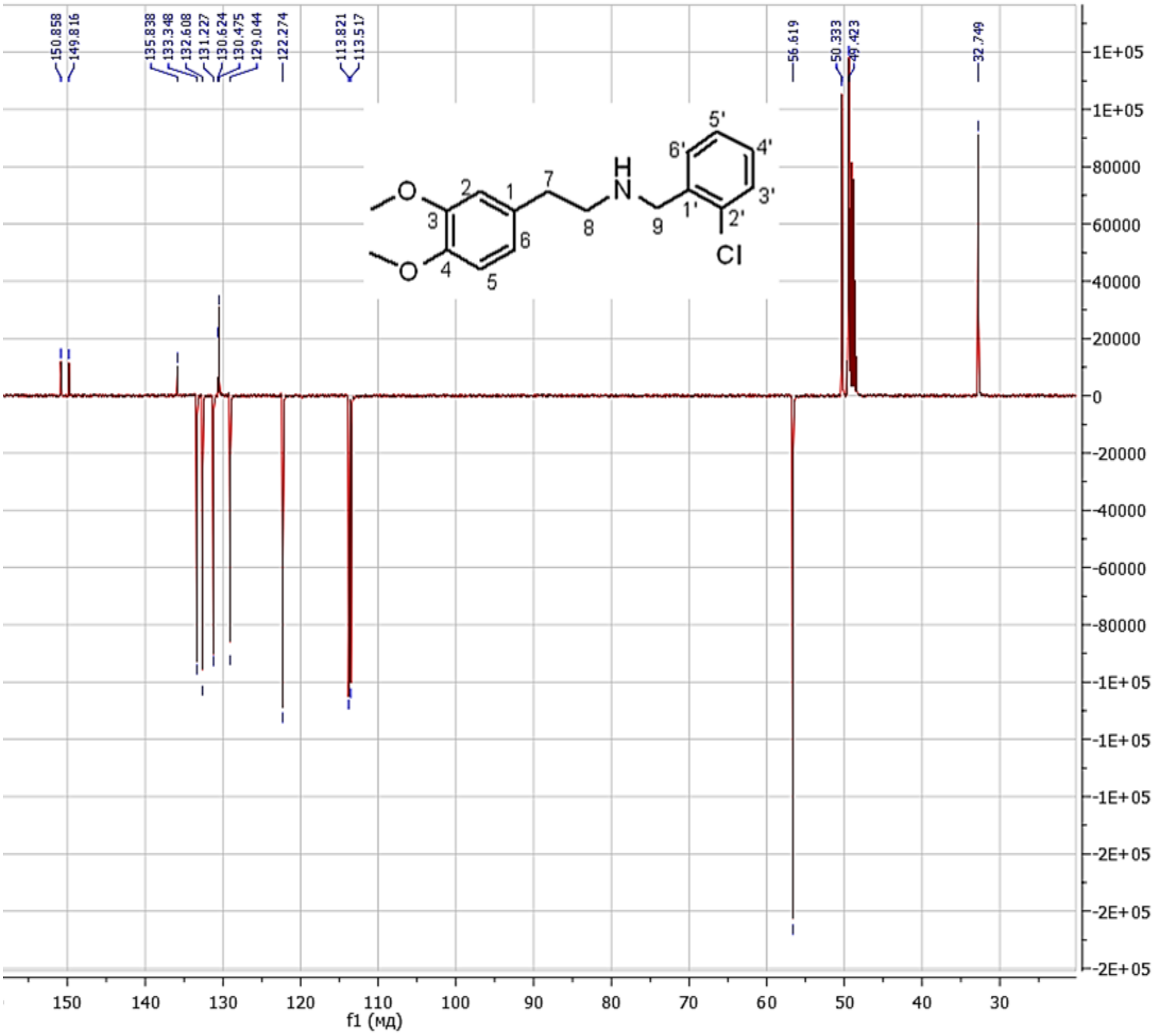


**Figure S26.** $^{13}C$ NMR spectrum of **9** (101 MHz, $CD_3OD$)

***N*-(2-Bromobenzyl)-2-(3,4-dimethoxyphenyl)ethanamine hydrochloride (34H-NBBr, (10)).**

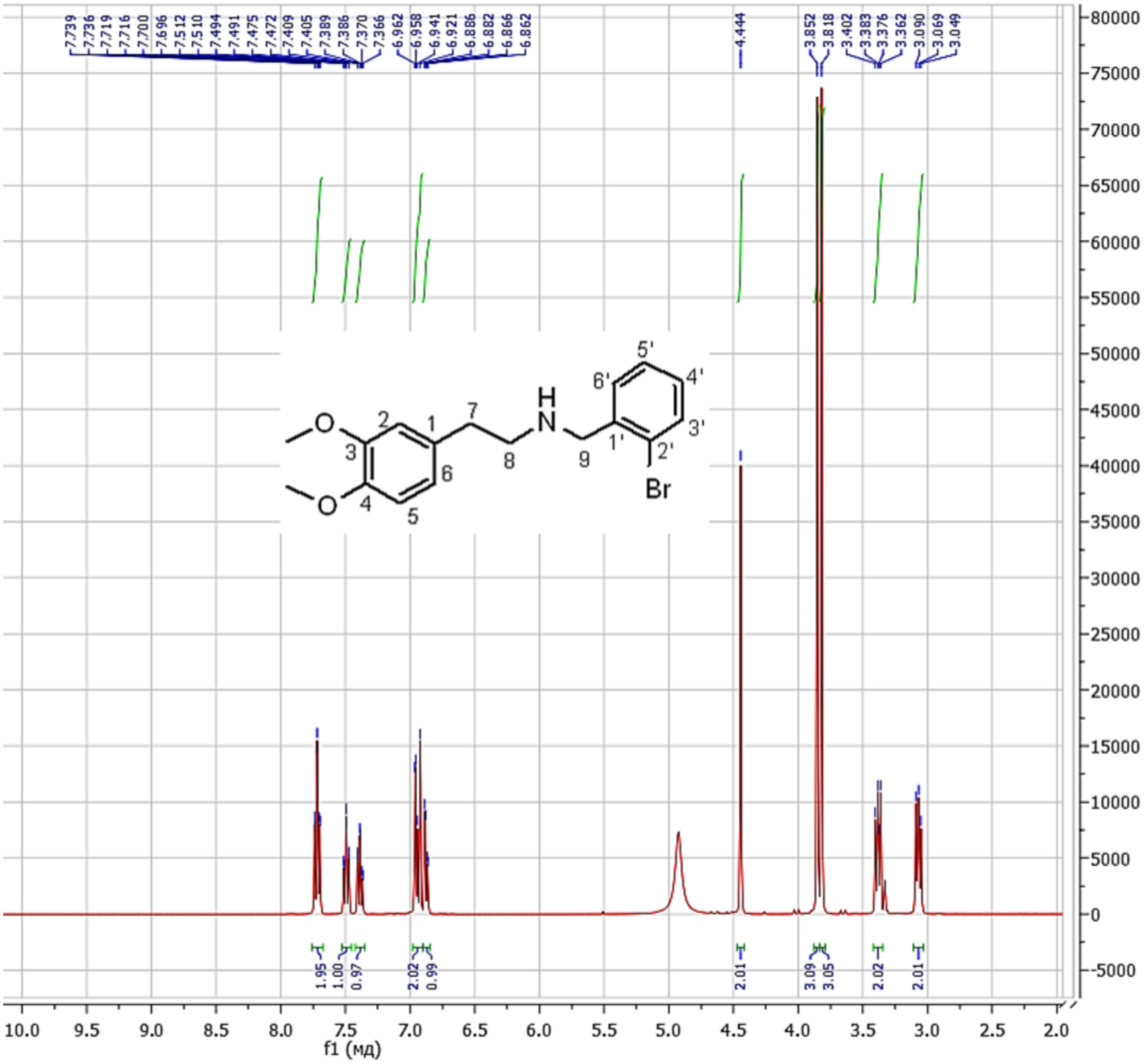

**Figure S27.** [1]H NMR spectrum of **10** (400 MHz, $CD_3OD$)

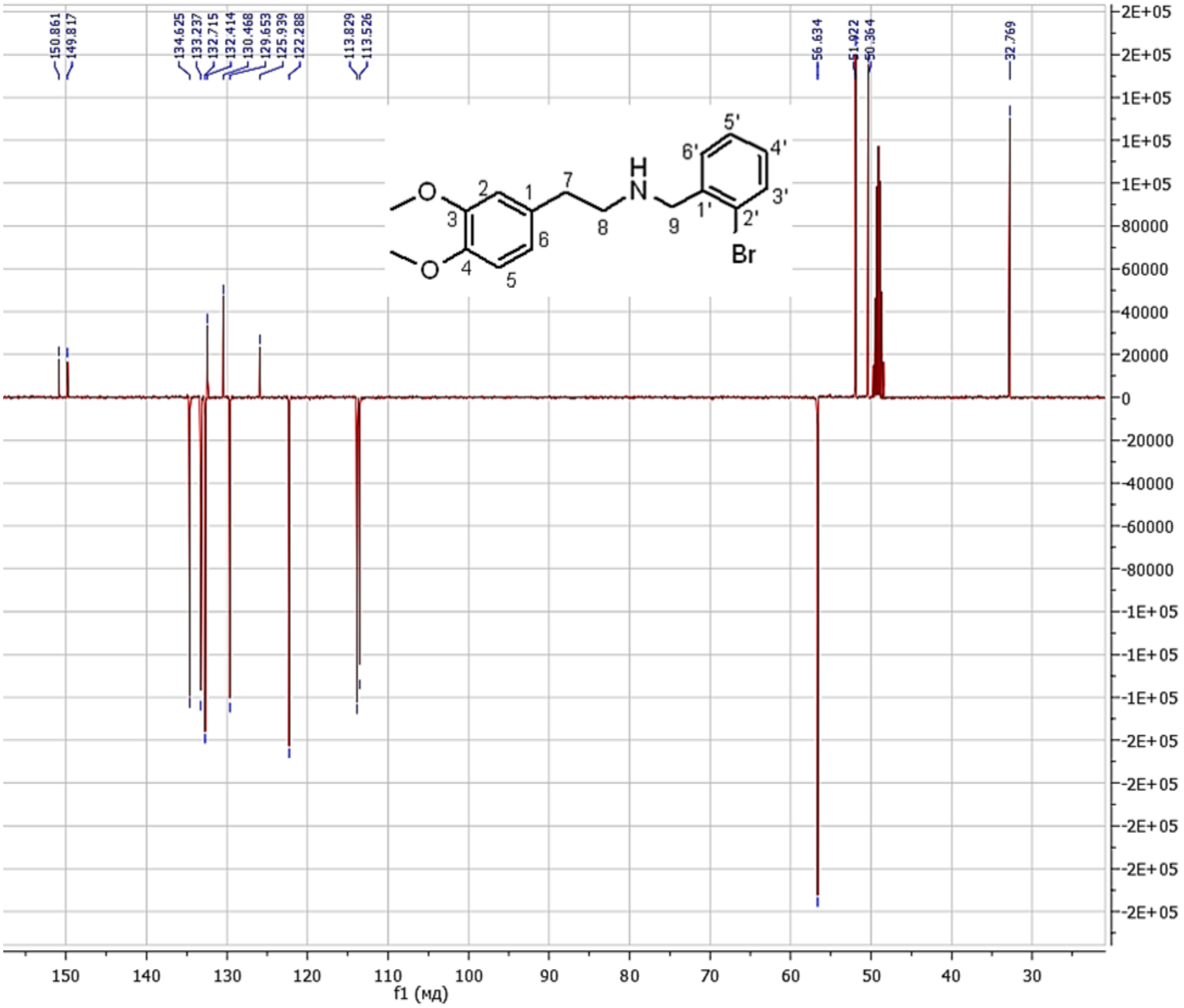


**Figure S28.** $^{13}$C NMR spectrum of **10** (101 MHz, $CD_3OD$)

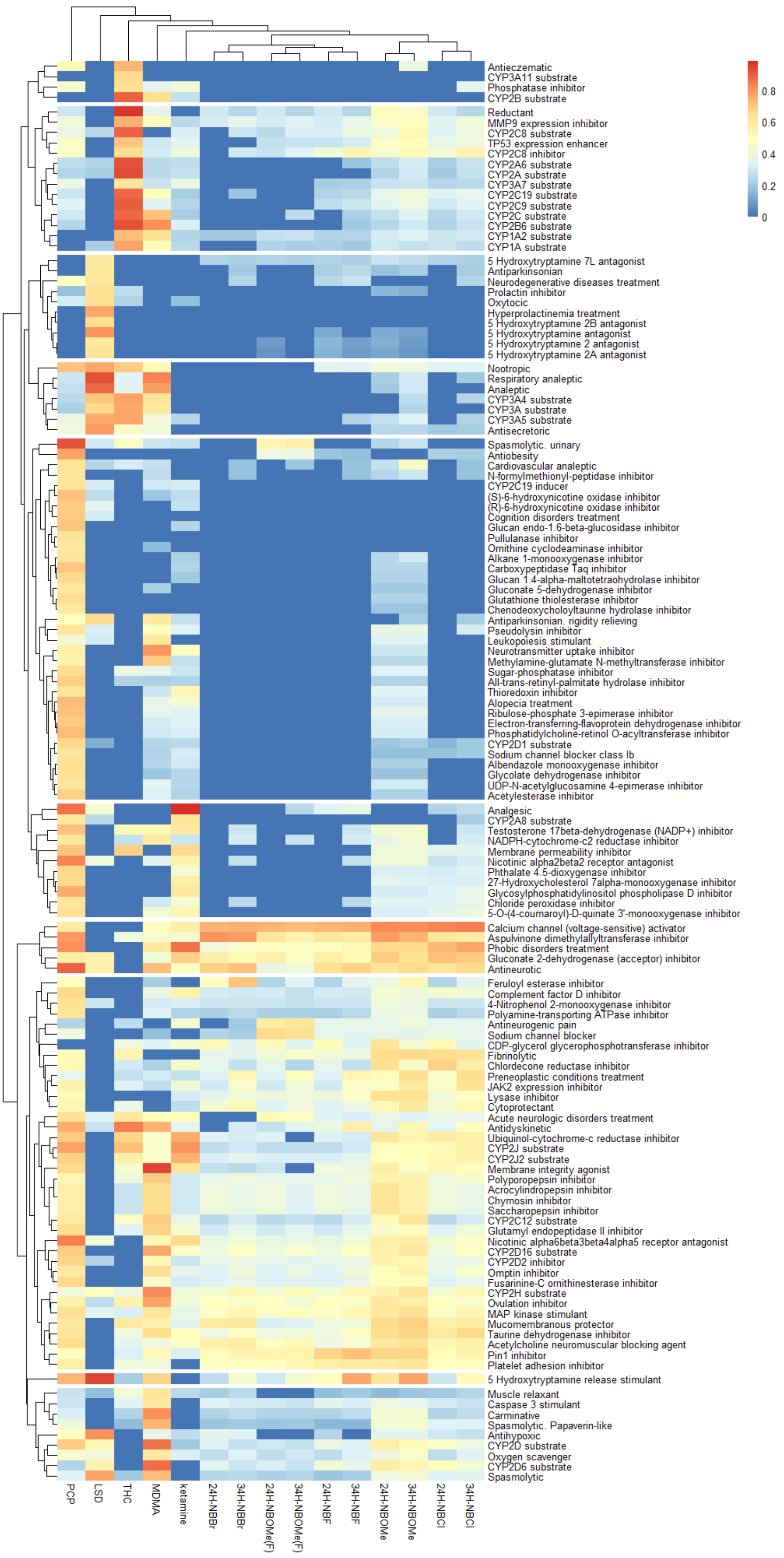


**Figure S29.** Heatmap representing probability of compounds activities (Pa, %) assessed using the PASSonline software with hierarchical clustering constructed using PCP, that was excluded from the final analyses.

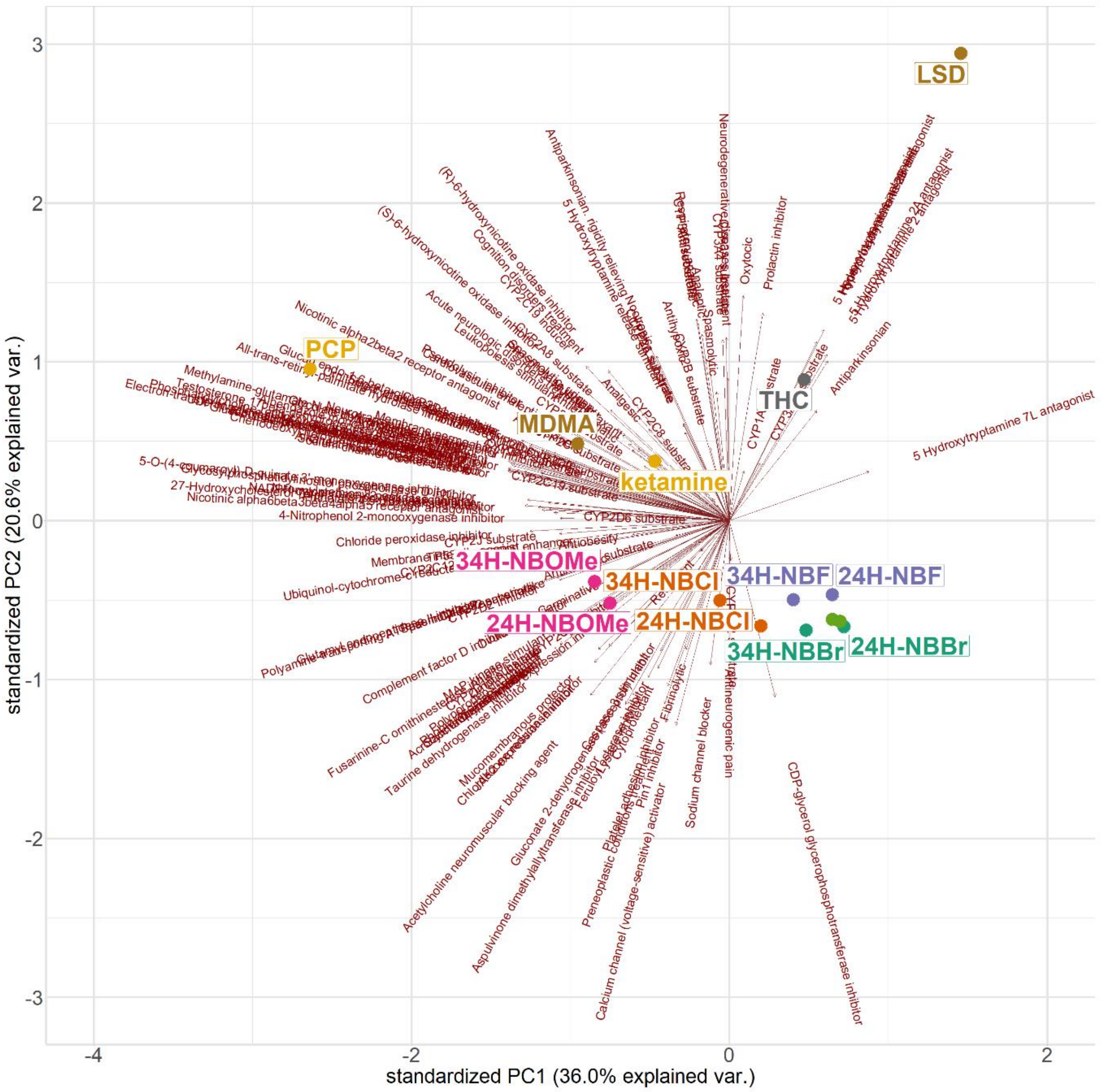


**Figure S30.** The principal component analysis results represented using the principal components with the most % of explained variance based on the compounds probability of activity, constructed using PCP, that was excluded from the final analyses. The arrows correspond to an activity loading.

**CRediT authorship contribution statement**

Konstantin A. Demin (Investigation) (Methodology) (Project administration) (Validation) (Writing - original draft) (Writing - review and editing), Olga V. Kupriyanova (Investigation) (Methodology) (Formal analysis) (Writing - Original Draft) (Writing - review and editing), Vadim A. Shevyrin (Investigation) (Methodology) (Formal analysis) (Writing - Original Draft) (Writing - review and editing), Ksenia A. Derzhavina (Investigation) (Formal analysis) (Writing - original draft) (Writing - review and editing), Nataliya A. Krotova (Investigation) (Formal analysis) (Writing - review and editing), Nikita P. Ilyin (Investigation) (Formal analysis) (Writing - review and editing), Tatiana O. Kolesnikova (Investigation) (Writing - Original Draft) (Writing - review and editing), David S. Galstyan (Investigation) (Writing - Original Draft) (Writing - review and editing), Iurii M. Kositsyn (Investigation) (Writing - Original Draft) (Writing - review and editing), Abubakar-Askhab S. Khaybaev (Investigation) (Writing - review and editing), Maria V. Seredinskaya (Investigation) (Writing - review and editing), Yaroslav Dubrovskii (Investigation) (Methodology) (Writing - review and editing), Raziya G. Sadykova (Investigation) (Writing - review and editing), Maria O. Nerush (Investigation) (Writing - review and editing), Mikael S. Mor (Investigation) (Writing - review and editing), Elena V. Petersen (Methodology) (Writing - review and editing), Tatyana Strekalova (Methodology) (Writing - review and editing), Evgeniya V. Efimova (Investigation) (Methodology) (Writing - review and editing), Dmitrii V. Bozhko (Investigation) (Methodology) (Writing - review and editing), Vladislav O. Myrov (Investigation) (Methodology) (Writing - review and editing), Sofia M. Kolchanova (Investigation) (Methodology) (Writing - review and editing), Aleksander I. Polovian (Investigation) (Methodology) (Writing - review and editing), Georgii K. Galumov (Investigation) (Methodology) (Writing - review and editing), and Allan V. Kalueff (Conceptualization) (Funding acquisition) (Methodology) (Project administration) (Resources) (Supervision) (Validation) (Writing - review and editing)
